# Meiotic cohesin Rec8 imposes fitness costs on fission yeast gametes favoring the evolution of parental bias in gene expression

**DOI:** 10.64898/2026.03.07.710300

**Authors:** Celso Martins, Harry Booth, Clàudia Salat-Canela, Zena Hadjivasiliou, Aleksandar Vještica

## Abstract

Differences between partner gametes, which evolved repeatedly in eukaryotes, can contribute to the evolution of the sexes, sexual selection and non-Mendelian inheritance^1– 4^. Yet, the empirical evidence for how functional asymmetries arise between initially equivalent gametes is limited. Here, we combine theoretical and experimental approaches in the fission yeast *Schizosaccharomyces pombe* to show how selective pressures acting concurrently on gametes and zygotes drive the evolution of gamete differences. We find that despite being morphologically identical, P- and M-type partner gametes invest asymmetrically in zygotic development by contributing different amounts of conserved meiotic cohesins. P-gametes preferentially produce thve Rec8^5^ cohesin that increases zygotic fitness but reduces gamete viability, revealing a trade-off between reproductive success and gamete survival. We demonstrate that this asymmetry is mediated by partner-specific communication and model its evolutionary dynamics using empirically determined parameters. Our results support classical theoretical predictions for the evolution of gamete differences and provide a mechanistic understanding of how molecular asymmetries between partners can originate from opposing selection pressures acting in species that lack morphologically distinct gametes.

---

During sexual reproduction, partner gametes fuse to produce a zygote. Zygotic development and fitness are often determined by factors asymmetrically provided by the two parents. In oogamous species, such as animals and flowering plants, large oocytes provision most of the zygote resources, whereas small and numerous sperm are specialized for motility and genome delivery. In contrast, in isogamous species, zygotes rely on similar contributions from the two morphologically identical parents, or isogametes, that differentially express only a handful of mating-type-specific genes^6^. How gamete asymmetries arose in isogamous ancestors is a fundamental but open question in biology. The prevailing theoretical models for the evolution of anisogamy — first formalized by Parker, Baker and Smith^7^ — propose that asymmetries between isogametes arise under opposing pressures to 1) increase zygote provisioning, which selects for large, high-investment gametes, and to 2) proliferate gamete numbers, which favors small, low-investment gametes. However, empirical measurements of the proposed selective pressures are largely lacking, as is evidence of molecular mechanisms by which such evolutionary forces can act concomitantly in ancestral isogamous species. In this work, we use fission yeast to show that functional asymmetries operate during sex between isogametes, and we identify molecular basis for the selection pressures that drive the evolution of these asymmetries.

## RESULTS

### Partner gametes asymmetrically produce meiotic cohesins

During sexual reproduction, the fission yeast produces morphologically indistinguishable partner P- and M-gametes (**Fig.1A**), yet the P-parental genome is preferentially utilized to initiate zygotic gene expression^8^. Whether gametes differentially invest into zygotic fitness — for example, by depositing zygotic factors before partners fuse — has not been reported. Previous studies suggested that meiotic recombination genes, which have strictly zygotic functions, may become induced already during mating^9–11^. To directly test this hypothesis, we compared the transcriptomes of non-mating populations, comprised of a single gamete type, with those of mating populations where gametes formed pairs but failed to fuse due to deletion of the formin *fus1*^12,13^ (**Fig.S1A**). Mated gametes showed >2-fold upregulation of 255 protein coding genes (**Fig.1B**), which included early meiotic genes^9^ and genes functionally associated with the meiotic cell cycle and recombination (**Fig.S1B-S1E**). Transcripts for the meiotic cohesin subunits Rec8 and Rec11, and the meiotic recombination factor Rec25, were as abundant as transcripts encoding key mating regulators and housekeeping genes (**Fig.S1F**). These results suggest that transcription of meiotic genes may start before gametes fuse.

**Figure 1.**
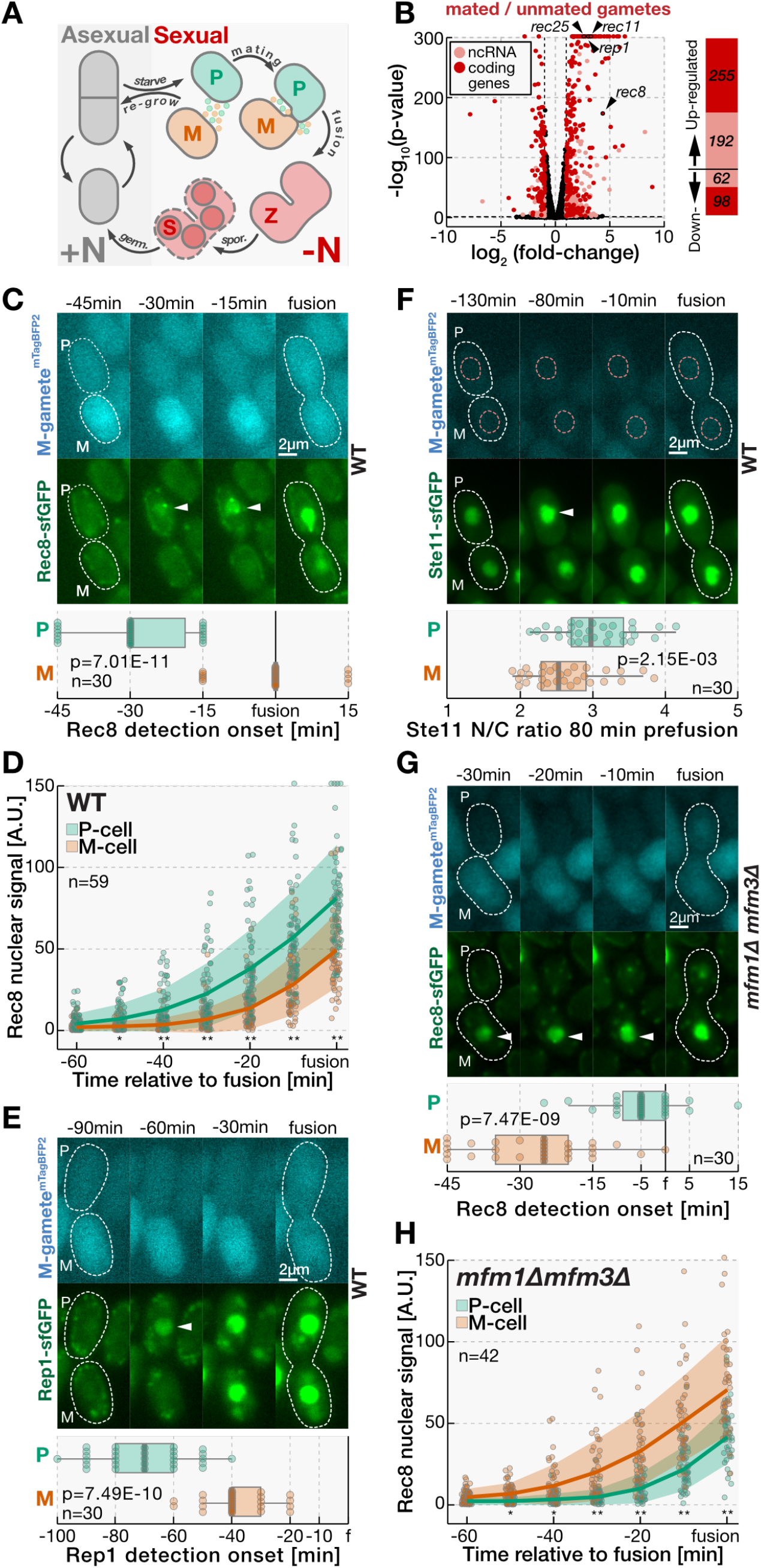
Pheromone-induced signaling drives early expression of meiotic genes in P-gametes. **(A)** Schematic of fission yeast lifecycle. Mitotic proliferation occurs in the presence of nutrients (+N, gray cells) and depletion of a nitrogen source (-N) induces cells to arrest in the G_1_ cell-cycle phase and to differentiate into partner P- and M-gametes (*P* and *M*, respectively). Mating type switching in homothallic *h90* strains^50^ produces both types of gametes, whereas heterothallic *h+* and *h-* strains produce only P- and M-gametes, respectively^51^. Partners communicate using a pair of diffusible pheromones (green and orange dots) and cognate receptors to orient growth that brings them into physical contact. Upon gamete fusion, zygotes (*Z*) rapidly enter the pre-meiotic S-phase^23^, followed by meiotic divisions that produce four haploid spores (*S*) within an ascus. When nitrogen source is restored, spores, as well as unfused gametes, resume mitotic cycling^18,52^.**(B)** Volcano plot shows protein-coding (red) and non-coding (pink) polyA+ genes that are differentially expressed (|log_2_(fold change)|≥ 1, p ≤ 0.05) between non-mated and mated gamete populations 15 h after removal of a nitrogen source (**Fig.S1A**). Black dots indicate genes that are not differentially expressed. Arrowheads indicated meiotic genes upregulated in mating. **(C)** Micrographs show Rec8-sfGFP (green) during homothallic wild-type cells mating. The M-gamete-specific *p*^*mam1*^ promoter regulates expression of mTagBFP2 (blue), which we used to determine the gametes mating type and time of fusion. Boxplots report the onset of Rec8-sfGFP detection in P- and M-gametes. Note that at -45 min cells show only the faint background fluorescence in the form of dots and strings, most likely corresponding to mitochondria. Note that Rec8-sfGFP produces a diffuse nuclear signal in addition to prominent nuclear foci that are consistent with reported Rec8 enrichment at centromeres^5^. Note Rec8-sfGFP expression prior to cell fusion and first in the centrally positioned nucleus of the P-gametes (arrowheads, boxplots), followed by M-gametes, and continued production in zygotes (**Fig.S2A**). We note that Rec8 expression asymmetry between partners was not due to the mTagBFP2 since heterothalic P-gametes carrying the red nuclear marker initiated Rec8-sfGFP production before their unlabeled partners (**Fig.S6B**). **(D)** The graph reports Rec8-sfGFP nuclear signal during wild-type matings in P- and M-gametes, as indicated. Note that P-gametes induce Rec8 earlier and to higher levels than M-gametes. **(E)** Micrographs show Rep1-sfGFP (green) during wild-type matings. The mTagBFP2 (blue) as in (C). Boxplots report the onset of Rec8-sfGFP signal detection in P- and M-gametes. Note that Rep1-sfGFP produces a diffuse nuclear signal which appears first in P-gametes (arrowheads) and subsequently in M-gametes, with continued presence in zygotes (**Fig.S2I**).**(F)** Micrographs show Ste11-sfGFP (green) during wild-type matings. The mTagBFP2 (blue) as in (C). Nuclear boundaries, which we visualized with the nucleus-targeted ^NLS^mCherry fluorophore (**Mov. 1**), are indicated with orange dashed lines. Boxplots report the Ste11-sfGFP nucleocytoplasmic (N/C) ratio in P- and M-gametes 80 min before cell fusion. Note that Ste11-sfGFP signal increases during mating and that the P-gamete shows higher nuclear Ste11 levels than the M-gamete at 80 min before fusion (arrowhead, boxplots). **(G)** Micrographs show Rec8-sfGFP (green) during mating of homothallic strains lacking the *mfm1* and *mfm3* genes. The mTagBFP2 (blue) as in (C). Boxplots report the onset of Rec8-sfGFP signal detection in P- and M-gametes. Note that Rec8-sfGFP signal is produced first in M-gametes (arrowheads, boxplots) during mating of *mfm1Δ mfm3Δ* partners. **(H)** The graph reports Rec8-sfGFP nuclear signal during mating between P- and M-gametes lacking *mfm1* and *mfm3* genes. Note that M-gametes induce Rec8 earlier and to higher levels than M-gametes. All timepoints are relative to partner fusion and gamete mating types are indicated (P/M). Scale bars are indicated in micrographs, white dashed lines indicate cell boundaries visualized from brightfield images. Dots represent individual datapoints and we indicate the number of analysed mating events (n). We show Kruskal-Wallis test p-values either as exact values or indicate value range with asterisks as 1.00E-3 > * > 1.00E-6 > **. In boxplots, box edges indicate 25th and 75th percentiles, the central line denotes the median, and whiskers extend 1.5 times the interquartile range. In line graphs, lines connect mean values and shaded areas report standard deviation.

To characterize the production of meiotic proteins in the two gametes during wild-type matings, we fused the green fluorescent protein sfGFP^14^ with the endogenous Rec8 in homothallic cells, which undergo mating-type switching to produce both P- and M-gametes (**Fig.1C**). We tracked cytosolic mixing as partners fuse using the blue fluorescent protein mTagBFP2^15^ expressed in M-gametes (**Fig.1C**). Consistent with transcriptomic analyses, Rec8-sfGFP was produced during mating and before cell fusions (**Fig.1C, Mov.1A**). We detected Rec8-sfGFP also in wild-type gametes that mated but ultimately failed to fuse with a partner (6.94±1.65% of n>1400 mating cells; detailed below). Strikingly, Rec8-sfGFP signal appeared 29±9 min earlier in P- than M-gametes (**Fig.1C-1D**). As partners produced Rec8-sfGFP at comparable rates, the signal in the two gametes remained asymmetric, with M-cells expressing a third of Rec8 of their partners throughout mating (**Fig.1D, S2B**). Tagging endogenous Rec11 or Rec25 with sfGFP revealed similar expression dynamics – both proteins were produced prefusion and first in P-gametes (**Fig.S2C-S2D**). We conclude that key meiotic proteins are produced in a mating-type-specific manner prior to partner fusion and predominantly by P-gametes.

### Pheromone-driven partner communication generates the parental bias in Rec8 expression

How do partners bias the Rec8 production? Imaging of nascent transcription sites showed that during mating, *rec8* transcription is initiated first in P-gametes (**Fig.S2E-S2F**). Rec8 expression in gametes was not regulated by the P-cell-specific transcription factor Pi^5^ (**Fig.S2G**). Instead, Rec8-sfGFP induction dynamics (**Fig.S2B**) suggested that both gametes regulate Rec8 expression with the same transcriptional machinery that is first activated in P-cells. Consistent with this hypothesis, deleting the Rep1 transcription factor^16,17^ severely reduced Rec8-sfGFP levels in both gametes (**Fig.S2H**). Importantly, the endogenous Rep1 tagged with sfGFP appeared in P-gametes 32±16 min earlier than M-gametes (**Fig.1E, S2I, Mov.1B**). Rep1 preceded Rec8 production by comparable periods of time in both parents (41±19 and 37±14 min in P- and M-gametes, respectively; p=0.11). The *rep1* promoter activation, and the *rep1* transcription, were also asymmetric and occurred more robustly in P-cells (**Fig.S2J-S2M**). Since these results suggest that the parental bias in Rec8 expression is caused by an earlier asymmetry in *rep1* transcription, we proceeded to investigate *rep1* regulation. We monitored the dynamics of Ste11 HMG family transcription factor that induces Rep1 and numerous mating genes^10,11,18,19^. Expectedly^20^, in unmated gametes, the sfGFP-tagged endogenous Ste11 localized to the cytosol and the nucleus (**Fig.1F, Mov.1C**). As gametes engaged partners, Ste11-sfGFP nuclear targeting was significantly higher for P-than M-gametes in the period preceding the Rep1 production (**Fig.1F, S2N**). Thus, Ste11 localization, which correlates with its ability to activate transcription of target genes^20^, is differentially regulated in partner gametes, and likely responsible for the parental bias in Rep1 and Rec8 expression.

To investigate Ste11 regulation, we tested the roles of diffusible pheromones that control its localization and activity^10,11,19,20^. Since P-gametes perceive the partner’s pheromone^19^, we deleted *mfm1* and *mfm3* paralogs of the three M-pheromone genes, which did not prevent sexual reproduction (**Fig.S3A**). Remarkably, in *mfm1Δ mfm3Δ* mutant pairs, Ste11-sfGFP nuclear targeting was stronger in M-than P-gametes (**Fig.S3B-S3C, Mov.2A**) and Rep1-sfGFP appeared in M-gametes 37±15 min before P-gametes, effectively reversing the asymmetry observed in wild-type pairs (**Fig.S3D, Mov.2B**). Furthermore, *mfm1Δ mfm3Δ* mutant M-cells induced Rec8-sfGFP 22±15 min ahead of P-cells (**Fig.1G-H, S3E, Mov.2C**). We conclude that distinct properties of pheromone signaling in P- and M-gametes lead to parental bias in the dynamics of Ste11, and expression of Rep1 and Rec8. Collectively, our results support a model where differences in pheromone signaling between gametes lead to distinct Ste11 nuclear targeting and activity that generate parental bias in the expression of Rep1 and its target genes, including meiotic segregation and recombination factors Rec8, Rec11 and Rec25.

### Rec8 produced in gametes promotes zygotic success and offspring survival

Having observed differential Rec8 provisioning by the two parents, we proceeded to test whether *rec8* P- and M-alleles make distinct contributions to zygote development. We measured the viability of spores produced in crosses of *rec8* mutant and wild-type heterothallic strains, which do not undergo mating-type switching and produce gametes of a single mating type. Strikingly, deleting *rec8* only in the P-gamete reduced progeny viability by 47±8% — nearly twice the 24±7% reduction observed upon deleting *rec8* in M-gametes only (**Fig.2A**). In contrast, loss of a single *rec8* copy did not reduce the viability of spores produced by azygotic meiosis^21^ in diploid strains since heterozygous *rec8Δ/rec8*^*WT*^ and homozygous *rec8*^*WT*^*/rec8*^*WT*^ diploids produced spores of similar viability (**Fig. S4A**). Two lines of evidence show that during sex, *rec8* parental alleles make quantitatively distinct contributions to accuracy of meiosis^5,22^; First, timelapse imaging of the fluorophore tagged Hht1 histone H3^23^ revealed that meiotic divisions failed to segregate chromatin into four even masses in 30±5% of zygotes produced by *rec8*Δ P-parent, compared to 14±6% of zygotes with *rec8Δ* M-parent (**Fig.S4B-S4C, Mov.3**). Second, we monitored the inheritance of two parental reporter alleles of the same genomic locus (**Fig.S4D**) and found that meiotic allele segregation was perturbed more strongly upon deleting *rec8* in P-gametes compared to M-gametes (23±3% and 10±2% of mis-segregation events, respectively; **Fig.2B**). Consistent with meiotic defects, the frequency of asci with aberrant spore number was 13±3% and 5±2% for crosses where *rec8* was deleted in either P- or M-gametes, respectively (**Fig.2B**). Thus, during sex, parent-of-origin effects govern Rec8 contributions to allele segregation and spore viability, with the P-allele exerting significantly greater impact.

Even though M-pheromone defective mutants (*mfm1Δ mfm3Δ*) strongly delayed Rec8 expression in P-gametes (**Fig.1G-1H**), we detected only mild defects in sporulation and meiotic allele segregation (**Fig.S3A, S4E**). These findings suggested that during sex between *mfm1Δ mfm3Δ* mutants, *rec8* P-allele has a limited role. Indeed, *mfm1Δ mfm3Δ* mutant zygotes lacking the *rec8* P-allele showed 10±2% fewer meiotic defects than those lacking the M-allele (**Fig.S4E**). Thus, in contrast to wild-type, meiosis in *mfm1Δ mfm3Δ* zygotes predominantly relies on *rec8* supplied by M-gametes. This suggested that, in principle, Rec8 provisioned by either gamete supports its functions. Consistent with this idea, expressing Rec8 from the *p*^*mam1*^ promoter, which drives expression of the pheromone receptor in M-gametes early in mating, fully rescued meiotic defects in zygotes that lacked native *rec8* loci (**Fig.S4F-S4G, Mov.4A**). The loss of native *rec8* alleles was also rescued when Rec8 was induced from the *p*^*pak2*^ promoter, which drives expression of the Pak2 polarity kinase during mating and in both gametes (**Fig.S4G-S4H, Mov.4B**). However, when Rec8 was placed under the *p*^*mei3*^ promoter, which is rapidly and exclusively activated following gamete fusion to express the Mei3 meiotic inducer^24,25^, it failed to efficiently complement the native *rec8* loci and mating produced 30±3% of defective asci (**Fig.S4G, S4I, Mov.4C**). These findings suggest that the roles of *rec8* parental alleles are not strictly determined by the parent of origin, and instead, relate to the time of Rec8 induction relative to gamete fusion.

Since gamete fusion is closely followed by pre-meiotic S-phase^24^ – and because sister chromatid cohesion requires cohesin loading onto chromosomes before DNA replication^26–28^ – we hypothesized that Rec8 production in gametes ensures sufficient levels at zygotic S-phase. To quantify Rec8 amounts produced from each parental allele at S-phase, we monitored crosses where one or both parents carried *rec8-sfGFP* and co-visualized the mCherry-tagged DNA replication fork component Pcn1^29^, which forms nuclear foci only during DNA replication^29^ (**Fig.2C, Mov.5**). Compared to expression from both parental copies, P- and M-alleles produced 75±18% and 29±13% of the Rec8-sfGFP at pre-meiotic S-phase (**Fig.2D**). DNA replication was initiated at comparable times post-fusion in all crosses (**Fig. S4J**). These findings suggested that the *rec8* M-allele cannot compensate for the loss of the P-allele because Rec8 production in M-gametes starts too close to S-phase. To test this idea, we performed crosses where P-gametes lacked the *rec8* gene and we delayed the premeiotic S-phase by 35±13 min by deleting one copy of the zygotic inducer *mei3* ^8,24^ (**Fig.S4K, Mov.6**). This delay allowed a 38±22% increase in Rec8-sfGFP production from the M-allele at S-phase (**Fig.2E-2F**) and, importantly, largely rescued meiotic defects caused by *rec8* P-allele deletion (**Fig.2G**). Mutating *mei3* did not suppress defects in control zygotes that lacked both *rec8* alleles (**Fig.2G**). We conclude that *rec8* induction in P-gametes ensures sufficient Rec8 levels at premeiotic S-phase to support sister chromatid cohesion.

To further understand how reduced Rec8 levels affect meiotic chromosomes, we used the *parS/ParB* reporter system^30^ to visualize the centromere-distal *lys3* and the centromere-proximal *rec6* loci (see Methods), which are located ∼3Mb and ∼4kb from the centromeres of chromosomes I and II, respectively (**Fig.2H, S5A-S5B, Mov.7-8**). In wild-type zygotes, replicated loci residing on sister chromatids appeared almost exclusively as a single dot indicative of tight cohesion (**Fig.2H-2I, S5A**). Rec8 from both parents was required for efficient sister chromatid cohesion at both the centromere-distal and centromere-proximal loci in zygotes (**Fig.2HI-2I, S5A**, ref.^31^). Importantly, in zygotes lacking one parental *rec8* allele — and more often in those lacking the P-than the M-allele — we observed that centromere-linked *rec6* loci on sister chromatids segregated to opposite poles (**Fig. 2H, 2J**), which is indicative of aberrant, equational first meiotic division. This is in contrasts to wild-type zygotes that carried out reductional meiosis I, where *rec6* loci residing on sister chromatids co-segregated together (**Fig. 2H, 2J**). Interestingly, deletion of the *rec8* P-allele had a more pronounced effect on the chromosome inherited from the P-parent (**Fig.2J**). The observed equational meiosis I suggested that ablation of one *rec8* parental allele reduces the ability of meiotic centromeres to orchestrate reductional meiosis^31^. Even though removal of one *rec8* parental allele reduced cohesion on chromosome arms (**Fig.2I, S5A**; ref.^31,32^), non-disjunction of homologs at meiosis I — where four *rec6* loci on replicated homologues segregate together^31^ — occurred very rarely and at rates comparable to wild-type zygotes (**Fig.S5B-S5C**). However, when we specifically impaired chromosome arm cohesion by deleting the meiotic cohesin Rec11^31,32^, homologue non-disjunction rates increased and the spore viability was reduced (ref.^31,33^ and **Fig. S5B-S5D, Mov.9**). Consistent with our observations that P-gametes are first to induce Rec11 (**Fig. S2C**), we find that the *rec11* P-allele contributed more to spore viability than the M-allele (**Fig.S5D-S5E**). Deleting both *rec11* and *rec8* in P-gametes did not show additive effects (**Fig.S5E**; ref.^33^). These results suggest that meiotic cohesins produced in P-gametes promote spore viability primarily through Rec8 functions at centromeres with a smaller role for Rec11 and cohesion at chromosome arms.

**Figure 2.**
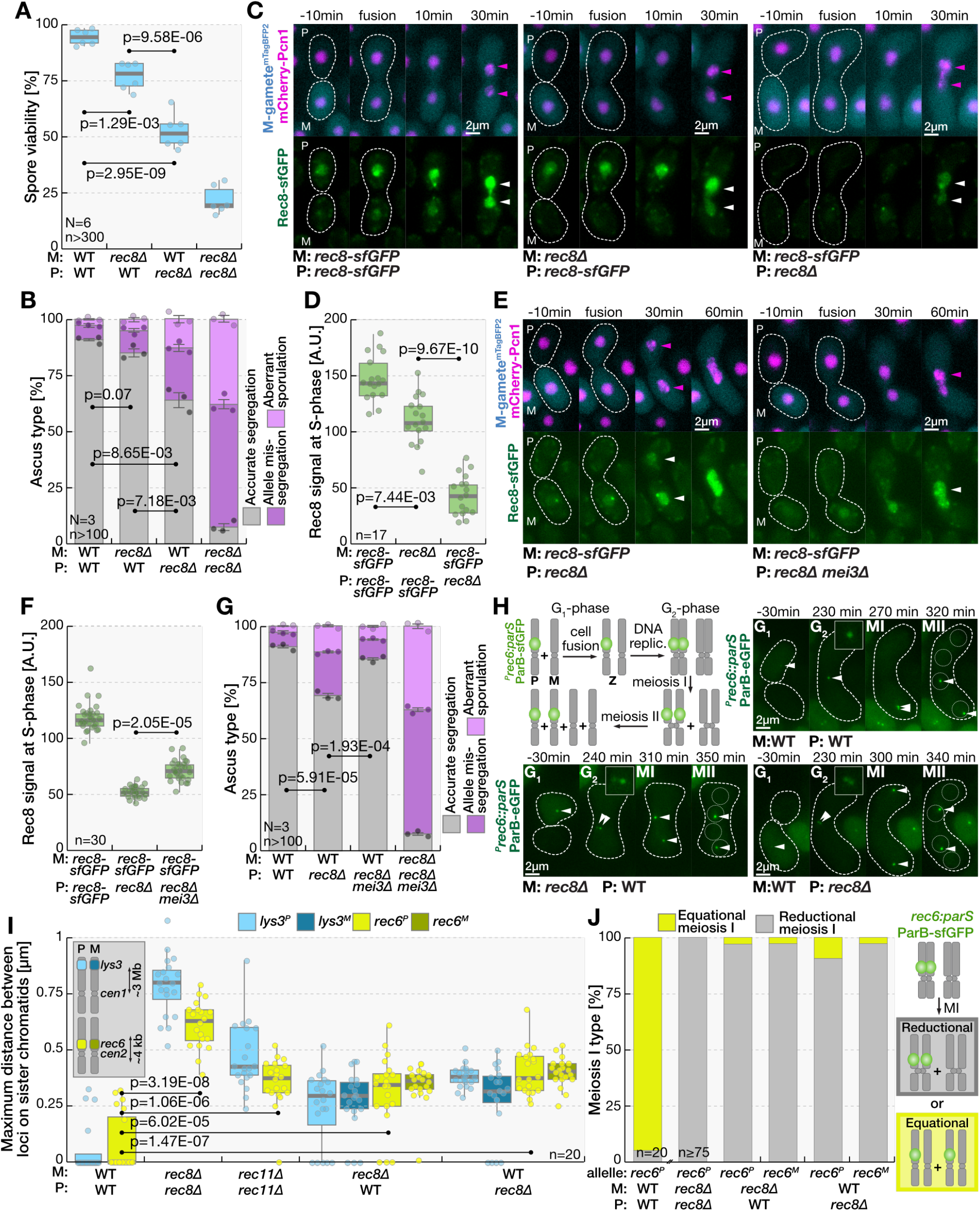
The early induction of Rec8 in P-gametes is required for successful meiosis. **(A)** Boxplots report the viability of spores produced in indicated crosses of wild-type and *rec8Δ* mutant gametes. Tukey test p-values are shown. Note that deleting *rec8* in the P-gamete reduces spore viability more than deleting *rec8* in the M-gamete. **(B)** Stacked histograms report the frequency of asci with accurate meiotic segregation of reporter alleles (grey; detailed in **Fig.S4D**), asci with mis-segregate alleles (purple) and asci containing aberrant spore number (pink) for indicated crosses between wild-type and *rec8Δ* mutant gametes. Welch test p-values are shown. Note that meiotic defects are more pronounced when *rec8* is deleted in P-than M-gametes only. **(C)** Micrographs show Rec8-sfGFP (green) expressed from either one or both parental alleles, as indicated. Cells co-express mCherry-Pcn1 (magenta), which during DNA-replication forms foci (magenta arrowheads) and a uniform nuclear signal outside S-phase. The M-gamete-specific *p*^*mam1*^ promoter regulates expression of mTagBFP2 (blue), which we used to determine the mating type of gametes and the time of fusion. Note that the Rec8-sfGFP signal at the onset of S-phase (white arrowheads) is lower when produced by the M-allele alone (right) than that produced by the P-allele alone (middle) or both parental alleles (left). **(D)** The boxplot reports Rec8-sfGFP levels at the onset of S-phase in zygotes produced in indicated crosses shown in (C). Dunn test p-values are shown. Note that Rec8 levels at S-phase are largely produced by the P-allele. **(E)** Micrographs show Rec8-sfGFP (green) expressed from the M-allele in indicated crosses where P-gametes lack the *rec8* gene and the *mei3* gene is either intact or deleted. The S-phase marker mCherry-Pcn1 and mTagBFP2 as in (C). Note that at the premeiotic S-phase onset (magenta arrowheads) is delayed when P-gametes lack the *mei3* gene, and that this allows for increased Rec8-sfGFP production from the M-allele (white arrowheads). **(F)** The boxplot reports Rec8-sfGFP levels at the onset of S-phase in zygotes produced in indicated crosses shown in (E). Dunn test p-values are shown. Note that deleting the *mei3* in P-gametes, which delays pre-meiotic S-phase, allows for higher Rec8 production from the M-allele by the time S-phase starts. **(G)** As in (B), stacked histograms report meiotic segregation accuracy for indicted crosses between wild-type and *rec8Δ* mutant gametes where the *mei3* gene in P-gametes is either intact or deleted. Welch test p-values are shown. Note that the deletion of the *mei3* P-allele, which delays pre-meiotic S-phase, largely suppresses meiotic segregation defects caused by the deletion of the *rec8* P-allele. **(H)** The schematic illustrates the wild-type meiotic dynamics for chromosomes (gray) inherited by the zygote (Z) from P- and M-parents, where the *rec6* P-allele located ∼4kb from the centromere (gray circle) is labelled (green). Micrographs show mating between indicated wild-type, *rec8Δ* and *rec11Δ* partners where P-gametes carry the *rec6* locus labeled with the *parS*^*c3*^/ParB^c3^-eGFP (green) and M-gametes carry the *rec6* locus labeled with the *parS*^*c2*^/ ParB^c2^-mCherry-NLS (**Mov.8**). Cell cycle phases are indicated (G_1_, G_2_, MI and post-MII). Note that the *rec6* locus-recruited *parS*^*c3*^/ParB^c3^-eGFP (arrowheads, insets) produces a single dot prior to meiosis in wild-type zygotes but two dots in zygotes that lack either parental *rec8* allele. Also note that at meiosis I, in wild-type zygotes the *rec6* loci co-segregate and in *rec8* mutants the foci separate. **(I)** As indicated, boxplots report the maximum separation between genomic loci (*lys3* or *rec6*, see inset) residing on sister chromatids of indicated parental chromosome (P or M) prior to meiosis in zygotes produced in indicated crosses of wild-type, *rec8Δ* and *rec11Δ* gametes. The genomic loci were visualized using the *parS*^*c3*^/ParB^c3^-eGFP reporters shown in (C) and **Fig.S5A**. Note that the removal of parental *rec8* alleles reduces loci cohesion in zygotes. **(J)** For indicated crosses between wildtype and *rec8Δ* mutant gametes, stacked histograms report the frequency of *rec6* loci residing on sister chromatids of indicated parental chromosome (P or M) co-segregating together (reductional division) or separating (equational division) at meiosis I. The visualized *rec6* loci are shown in panel (H). Note that the zygotes lacking the *rec8* P-allele show higher rates of aberrant, equational meiosis I than zygotes lacking the M-allele. All timepoints are relative to cell fusion and gamete mating types are indicated (P/M). Scale bars are indicated in micrographs, white dashed lines indicate cell boundaries visualized from brightfield images. Dots represent individual datapoints and we indicate the number of analyzed replicates (N) and relevant cell types (n). In stacked histograms, bars report means and error bars denote standard deviation. In boxplots, box edges indicate 25th and 75th percentiles, the central line denotes the median, and whiskers extend 1.5 times the interquartile range.

Collectively, our results are consistent with a model that by producing distinct amounts of meiotic cohesins ahead of S-phase, P- and M-parental alleles differentially contribute to the meiotic chromosome cohesion and centromere function needed for efficient meiotic divisions and production of viable spores.

### Rec8 causes genomic instabilities and reduced viability in gametes

While preceding results show that Rec8 production in gametes promotes zygotic fitness, we find that it comes at a fitness cost to gametes. In mating populations of Rec8-sfGFP homothallic strain, we observed gametes that induced Rec8 during mating but then failed to fuse with a partner. These gametes typically arose when two P-gametes competed for a mate and only one fused with a partner, leaving the unfused P-gamete with induced Rec8 (6.33±1.38% of mating events; **Fig.3A, Mov.10**) Additionally, when we interrupted mating by transferring mating mixtures to growth media, Rec8-expressing unfused gametes constituted 7.1 ± 3.4% of the population. These gametes were predominantly P-cells, as indicated by their lower expression from the M-gamete-specific promoter compared to Rec8-negative gametes (**Fig.S6A**), a conclusion we confirmed using labeled heterothallic partners (**Fig.S6B-S6C**). Thus, wild-type mating mixtures contain a sub-population of Rec8-expressing haploid gametes that fail to fuse and are predominantly P-mating type. Importantly, when the supplied nutrients triggered mitotic re-entry, 6.7±1.3% of Rec8-expressing gametes — or their daughters produced in first division — either lysed or arrested growth (**Fig.3B-3C, Mov.11**). In contrast, such defects arose in only 0.3±0.1% of gametes that did not induce Rec8 (**Fig.3C, Mov.11**). These results suggested that Rec8 expressed during mating compromises cell viability, which may explain why Rec8 is supplied in a mating-type-specific manner despite its crucial role for meiosis.

**Figure 3.**
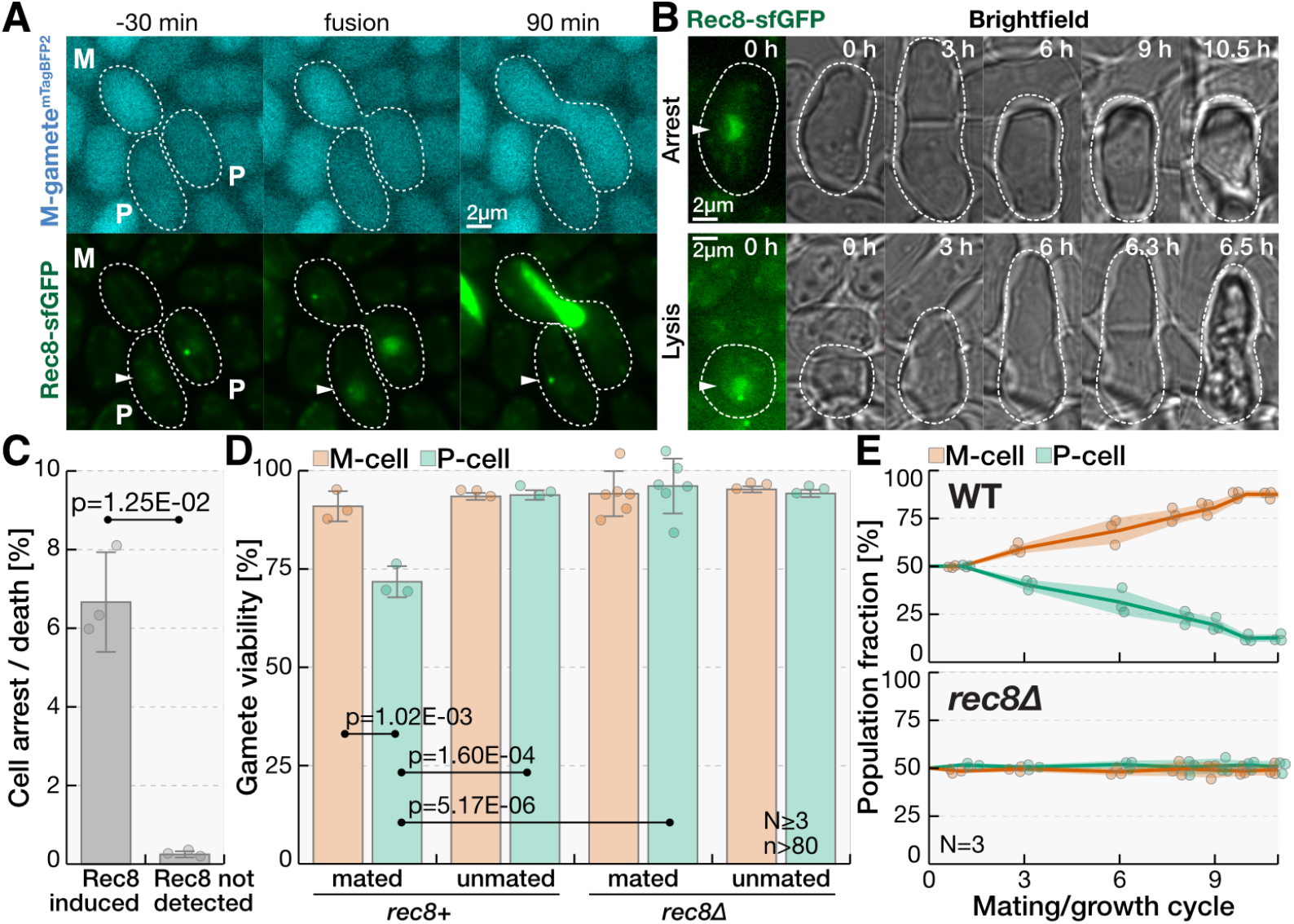
Rec8 accumulated during mating increases genomic instabilities and reduces fitness of P-gametes. **(A)** Micrographs show Rec8-sfGFP (green) in homothallic mating mixtures where two P-gametes compete for one M-partner. The M-gamete-specific *p*^*mam1*^ promoter regulates expression of mTagBFP2 (blue), which we used to determine the gametes mating type and time of fusion. Indicated times are relative to cell fusion. Note that Rec8-sfGFP is expressed pre-fusion in both outlined P-gametes (arrowheads) and that one of them fails to fuse with a partner. **(B)** The timelapses of brightfield images (gray) show homothallic cells returned to growth media after being induced to mate. Rec8-sfGFP (green) is shown at the timelapse onset. Note that for gametes that induce Rec8, mitosis leads to cell lysis (bottom) or the daughter cells that arrest growth (top). **(C)** The bar chart reports the frequency of growth arrest and cell death for mitotic progeny of gametes that either induced or did not show detectable Rec8 signal at the time homothallic mating mixtures were placed on growth media. Welch test p-values is shown. Note that the gametes with notable Rec8 expression have a ∼10-fold increase in cell death. **(D)** The bar chart reports viability (**Fig.S6D**) of mated and unmated P- and M-gametes with indicated genotypes. Tukey test p-values are shown. Note that mating reduces viability of P-gametes, and that this is dependent on the *rec8* gene. **(E)** Graphs report the fraction of P- and M-gametes after the indicated number of experimental cycles of mating and regrowth (**Fig.S6H**) for evolving populations of heterothallic *fus1Δ* cells that either carry wild-type *rec8* (top) or lack the *rec8* gene (bottom). Note that M-gametes outcompete P-gametes in populations with wild-type *rec8* allele but not in populations where *rec8* is deleted. Scale bars are indicated in micrographs, white dashed lines indicate cell boundaries visualized from brightfield images and gamete mating types are indicated (P/M). Dots represent individual datapoints and we indicate the number of analyzed replicates (N) and relevant cell types (n). In bar plots, bars report means and error bars denote standard deviation. In line graphs, lines connect means, and shaded areas report standard deviation.

The correlation between Rec8 induction and cell viability (**Fig.3C** and ref.^32^), prompted us to directly test the role of Rec8 in survival of unfused gametes. We prepared mating mixtures of *rec8+* and *rec8Δ* cells — where we deleted the *fus1* gene to abolish cell fusion — and we measured the viability of unmated gametes and gametes that formed mating pairs (**Fig.S6D**). Remarkably, mating reduced viability of P-gametes by 22±4%, and this viability loss was dependent on the intact *rec8* gene (**Fig.3D**). Deleting the *rec11* gene also increased viability of mated P-gametes albeit with a smaller effect (**Fig.S6E**), suggesting that recruitment of the meiotic cohesin complex to both centromeres and chromosome arms reduces gametes viability. In contrast, mating and *rec8* or *rec11* genes did not strongly impact the viability of M-gametes (**Fig.3D, S6E**). The loss-of-heterozygosity assay^32^ (**Fig.S6F**), which reports mitotic mis-segregation of alleles, showed a Rec8-dependent, 17±5-fold increase in mitotic defects for mated P-cells as compared to unmated cells with the same genotype (**Fig.S6G**). Taken together, our results show that Rec8 accumulated during mating of P-gametes that fail to fuse with a partner increases genomic instabilities and reduces their fitness.

Our results suggest that the asymmetry in *rec8* expression causes distinct fitness costs on P- and M-gametes and, thus, may affect how gamete populations evolve. To test this idea, we mated equal number of *fus1Δ* heterothallic P- and M-gametes, regrew the mated population and used this population to repeat the cycle (**Fig. S6H**). Strikingly, the fraction of P-gametes in the population decreased to 41±3% after only three experimental cycles and continued to decrease in subsequent cycles reaching 13±2% of the recovered cells after eleven mating rounds (**Fig.3E**). In parallel, we subjected *rec8Δ fus1Δ* populations of P- and M-gametes to the same cycles of mating and regrowth and found no evidence for selection of either gamete type (**Fig.3E**). We conclude that *rec8* differentially affects the fitness of the two gamete types during sexual and asexual reproduction, which may contribute to the evolution of asymmetric functions in gametes.

### Rec8-dependent costs and benefits drive the evolution of parentally biased Rec8 expression

Collectively, our experiments show that Rec8 expression prefusion produces opposing effects on the fitness of gametes and zygotes. This prompted us to hypothesize that, consistent with theoretical models for the evolution of anisogamy^6,7,34^, such conflicting Rec8 roles drive the evolution of asymmetric Rec8 production between gametes. To rigorously test this idea — and because existing models are rooted in the life cycles of animal-like, multicellular organisms — we developed an evolutionary model based on the fission yeast life cycle (**Fig.4A**, see Supplemental Notes S2). We consider a population where cells proliferate mitotically followed by a mating step. Each cell is associated with an evolvable genotype (*r*_*p*_,*r*_*m*_) which represents the quantity of Rec8 contributed at the point of fusion by the P- and M-gamete, respectively. Although our model is motivated, and subsequently parametrized, by the costs and benefits Rec8 elicits on gametes and zygotes, the framework we present is general and can be applied to the combined effect of meiotic factors beyond Rec8 and Rec11. We introduce probability *f* that partners fuse during mating. When fusion is successful, the cumulative amount of the Rec8 from both parents, *n=r*_*p*_*+r*_*m*_, specifies zygote fitness, which in our model defines the probability of successful meiosis, *P*(*n*) (**Fig.4A**). Alternatively, partners fail to fuse with probability *1-f* and the amount of Rec8 in gametes, *r*_*p/m*_, imposes a fitness cost. We model this cost with the probability that a gamete successfully returns to mitosis to produce daughter cells, *q*(*r*_*p/m*_) (**Fig.4A**). By subjecting the population to cycles of *r*_*p*_,*r*_*m*_ mutation, asexual and sexual reproduction, we find that populations initially set to a symmetric state (*r*_*p*_ =*r*_*m*_) can evolve to an asymmetric state (*r*_*p*_ ≠*r*_*m*_ ; **Fig.4B**; **Mov.12**). Thus, the fission yeast life cycle properties captured by our model allow evolution of asymmetric gamete investment.

**Figure 4.**
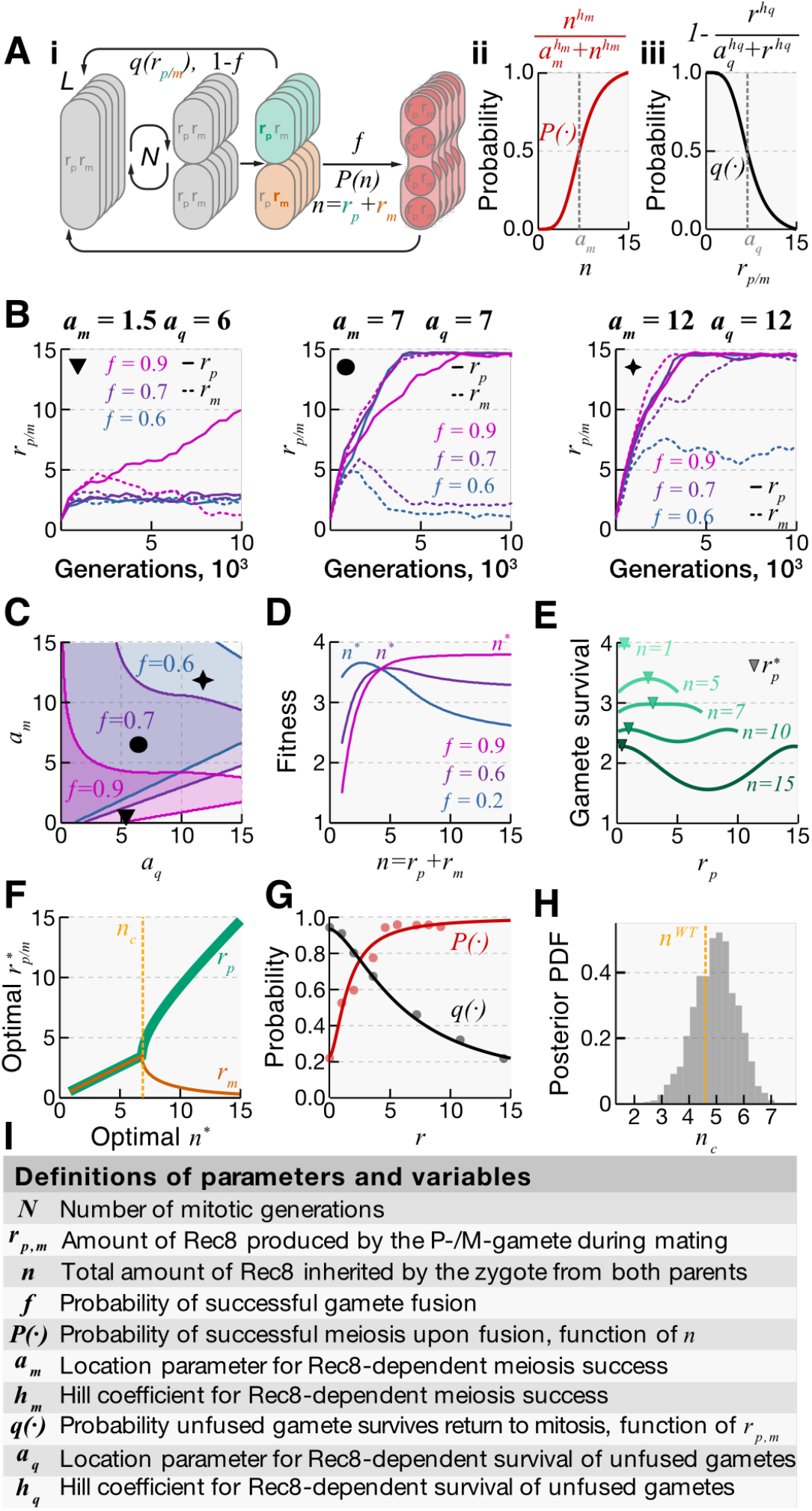
The selective pressures Rec8 places on gametes and zygotes drive asymmetric parental Rec8 contributions. **(A)** The schematic shows the lifecycle stages represented in the model (i). Each cell in a populationof size *L* is associated with an evolving genotype *r*_*p*_,*r*_*m*_ that defines the amount of the meiotic factor produced by the P- and M-parent during mating. Cells proliferate mitotically for *N* generations followed by sister cells mating step. We introduce a probability, *f*, that partners fuse during mating. If fusion is successful, zygotes receive *n=r*_*p*_*+r*_*m*_ meiotic factor investment and sporulate with probability *P*(*n*), which we assume takes a sigmoidal form (ii, red line). Non-fused gametes return to mitosis with probability *q*(*r*_*p/m*_), which we assume takes the sigmoidal form (iii, black line), and die otherwise. Parameter and variable definitions are given in panel **I. (B)** The average allocation of the meiotic factor in the P and M parents, *r*_*p*_ and *r*_*m*_ respectively, in populations evolving for 10,000 generations. The gamete that evolves the higher *r* value is designated as the P-parent (*r*_*p*_, solid lines) and its partner as the M-parent (*r*_*m*_, dashed lines). Depending on parameter choices the population evolves to a symmetric state where *r*_*p*_*=r*_*m*_ (when dashed and solid lines converge) or an asymmetric state where *r*_*m*_<*r*_*p*_ (dashed and solid lines diverge), suggesting that in theory asymmetry in meiotic factor contribution can theoretically be an evolutionary stable strategy under some conditions. The simulations were performed for three different fusion probabilities *f* (indicated by color) and for three different combinations of *a*_*q*_ and *a*_*m*_ parameters (indicated with ▾, •, ★ markers and values on the top) that define the shape of *P*(·) and *q*(·). For simulation details see Supplemental Notes S2. Parameter and variable definitions are given in panel **I. (C)** The regions in parameter space *a*_*q*_ versus *a*_*m*_ that result in evolution of asymmetric parental investment as the optimal reproductive strategy. Here, the optimal genotype 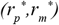 was defined as asymmetric if 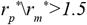 . Shaded regions correspond to those combinations of *a*_*q*_ and *a*_*m*_ which satisfy this condition, for given values of fusion probability *f* indicated by color. Markers (▾, •, ★) indicate the parameter sets used in the evolutionary simulations in panel **B**. Parameter and variable definitions are given in panel **I. (D)** The graph shows the overall fitness measured as expected number of offspring defined by 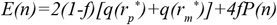 (**Eq.1** in the main text) versus total parental investment of meiotic factor *n*, for a given fusion efficiency *f* . We first determine the optimal genotype 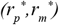 given the requirement that 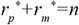. These values are then used to compute the fitness as a function of *n* using 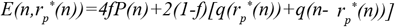, as detailed in Supplemental Notes S2. The plots suggest that as the probability of fusion increases the optimal total investment for zygotes increases. Parameter and variable definitions are given in panel **I. (E)** Gamete survival quantified as the expected number of offspring produced by unfused gametes, given by *2[q*(*r*_*p*_)*+q*(*n-r*_*p*_)], versus the P-parent investment *r*_*p*_ for given values of total parental investment of meiotic factor for the zygote set at indicated values of *n=r*_*p*_ *+r*_*m*_ . The triangle represents the optimum 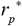 for each fixed value of *n*. When total zygote investment is low, gamete survival is high and is optimal at 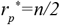 indicating that a symmetric investment of the meiotic factor by the two parents is optimal. As *n* increases, the overall fitness of unfused gametes, 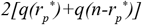, decreases and, importantly, the optimal 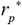 bifurcates away from the equal parental contribution at the curve middle, indicating that an asymmetric investment of the meiotic factor by the two parents becomes optimal. Parameter and variable definitions are given in panel **I. (F)** The optimal meiotic factor contribution by each parent 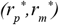 versus the optimal total parental investment *n*^*\**^. This is a summary plot of panel **E**, that allows visualization of the bifurcation point where gamete investment becomes asymmetric. Once the total investment is driven past a critical value *n*_*c*_ (orange dashed line), asymmetric parental investment emerges as an optimal strategy (the green and orange lines branch off). Parameter and variable definitions are given in panel **I. (G)** The functional forms of the probability of zygote survival as a function of total Rec8 investment *P*(·) (red line) and of gamete survival as a function of the amount of Rec8 produced by gametes *q*(·) (black line) fitted to the experimental data (dots; **Fig.S7C-S7E**). Parametrization of the functions using experimental data is detailed in the Supplemental Notes S2. Parameter and variable definitions are given in panel **I. (H)** The histogram shows the distribution of critical values *n*_*c*_ derived for 10,000 parameter sets sampled from the inferred posterior distribution for the parameters of *q*(·). The critical *n*_c_ value was defined to be the smallest *n* for which 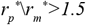. The dashed line indicates the total parental investment inferred for wild type matings (*n*_*WT*_*=4*.*6 A*.*U*.), which falls within the center of the predicted distribution suggesting that the costs and benefits of Rec8 expression identified experimentally are consistent with asymmetric gamete expression of Rec8. For details, see Supplemental Notes S2. **(I)** The definitions of parameters and variables in the theoretical model.

Numerical simulations indicated that the evolution of asymmetric gametes depended on the probability *f* that partners fuse, and the Rec8-dependent fitness benefits and costs, specified by the *a*_*m*_ and *a*_*q*_ parameters that define the shape of the functions *P*(·) and *q*(·), respectively (**Fig.4B**). We corroborated these results by noting that the expected number of offspring which originate from a genotype (*r*_*p*_,*r*_*m*_) after one life cycle, effectively defining its fitness, is given by (see Supplemental Notes S2)

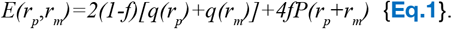

Using **Eq.1** we defined regions in the parameter space where gametes investment ratio, *r*_*p*_/*r*_*m*_, exceeded 1.5 (**Fig.4C**, shaded areas), confirming that the fusion efficiency *f* and the parameters that determine the shape of *P*(·) and *q*(·) collectively determine the optimal ratio between the two parental Rec8 contributions.

What sets the evolutionary optimum for the total amount of Rec8 provided by both parents? We evaluated fitness *E*(*r*_*p*_,*r*_*m*_) in terms of the total parental investment, *n=r*_*p*_*+r*_*m*_, which provides benefits if partners fuse (*P*(*n*) increases with *n*) and produces costs if fusion is unsuccessful (*2[q*(*r*_*p*_)*+q*(*n-r*_*p*_)] decreases with increasing *n*). Balancing these opposing effects, the overall fitness *E*(*r*_*p*_,*r*_*m*_) is maximized for a given value of *n*, which we term *n\** (**Fig.4D**). The fusion efficiency *f* acts as a weighing factor between benefits and costs of Rec8 investment (**Eq.1**) and is thus crucial in determining the optimal total investment *n\** (**Fig.4D**). For example, at low *f*, unfused gametes are numerous and the optimal total parental investment decreases because smaller *n* reduces the costs of returning to mitosis. As *f* increases, meiotic success of increasingly abundant zygotes dominates, and the overall fitness *E*(*r*_*p*_,*r*_*m*_) is maximal for higher total investment *n*, increasing *n\** (**Fig.4D**). In this way, the fusion efficiency *f* gates the contribution that the zygotic success *P*(·) has towards increasing the optimal total parental investment *n\** (**Fig.4D**). Importantly, when selection drives the optimal investment *n\** above a critical value, *n*_*c*_, the optimal gamete investments *r*_*p/m*_ bifurcate and *r*_*p*_ ≠*r*_*m*_ (**Fig.4E-4F**). The bifurcation occurs when splitting the optimal total investment *n\** evenly is more costly than distributing it unevenly between unfused partners (**Fig.4E**) — that is, when 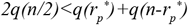, as determined by the shape of the *q*(*r*_*p/m*_) function and the value of *n* (see Supplemental Notes S2). Taken together, this analysis shows how the evolution of asymmetric Rec8 production in gametes critically depends on the fusion efficiency and quantitative effects of Rec8 on fitness of gametes and zygotes.

Are the Rec8 costs and benefits we identified experimentally sufficient to drive the evolution of asymmetric Rec8 provisioning? To address this question, we experimentally quantified *P*(*n*) and *q*(*r*_*p/m*_). Specifically, we quantified the effect of Rec8 levels on fitness of gametes and zygotes by measuring the spore viability and the unfused gamete survival for cells carrying different copy number of the *rec8* gene (**Fig.S7A-S7E**). We used Bayesian inference to parametrize *P*(·) and *q*(·) from this experimental data (**Fig. 4G, S7A-S7E**; see Supplemental Notes S2). Using this parametrization, and the high fusion efficiency observed earlier (*f=0*.*9*; **Fig.3A, S3A**), we produced a posterior distribution for the critical parental investment *n*_*c*_ above which we expect a substantial asymmetry in Rec8 investment to evolve (**Fig.4H)**. Crucially, the posterior likelihood of *n*_*c*_ being lower than the total Rec8 investment quantified from wild-type matings was 33.06% (**Fig.4H**; *n*_*WT*_=4.6a.u., **Fig.S7A**), consistent with a high likelihood of asymmetry arising at the experimentally quantified Rec8 levels and fitness effects. Taken together, our theoretical model, experimental measurements and statistical analysis show that an asymmetry in Rec8 investment is likely instigated by the conflicting selective pressures Rec8 places on gametes and zygotes.

## DISCUSSION

Unlike asymmetrically expressed mating factors such as pheromone receptors, Rec8 expression is under opposing selective pressures in gametes and zygotes. Rec8 production pre-fusion, while indispensable for early zygotic development, imposes a cost on unfused gametes, creating a conflict that is minimized through asymmetric parental investment and specialization between closely related partners. The results of our experimentally parametrized model mirror predictions from the classic Parker-Baker-Smith model of anisogamy evolution^7,34^, and provide direct empirical evidence for one of their central assumptions: that asymmetries can emerge through trade-offs in gamete-level and zygote-level fitness. Interestingly, fission yeast parental asymmetries do not require complex differences in gamete size but molecular-level specialization towards zygote investment. Since mutating *mfm* genes alone alters Rec8 expression in gametes, limited changes in pheromone communication may promote parental asymmetry. For example, M-cell specific transcription factor Mc may increase M-pheromone production either directly^35,36^ or by downregulating general signaling factors^36,37^. Likewise, P-cell sensitivity to the M-pheromone might be reduced by the P-cell specific transcription factor Pc^38^. Such limited regulatory asymmetries as a precursor to gamete dimorphism are consistent with mating-type loci in the anisogamous *Eudorina*, which contain only two sex-limited genes and two pairs of gametologs (*i*.*e*., genes with alleles in both mating haplotypes)^39^. Functional complementation between transcription factors residing at mating-type loci of isogamous, anisogamous and oogamous Volvocine species suggests that appearance of dimorphic gametes need not rely on their extensive sequence divergence but on changes in their regulatory networks^40^. Asymmetry in Rec8 production through duplication of the fission yeast *mfm* genes would be consistent with this model. Extensive sex-specific allele differentiation may in fact arise after the dimorphic gametes appear, as proposed for the MAT3 gene of *Volvox carter* ^41^.

Since gametic Rec8 production is regulated by the cascade that originates at the mating-type loci, selection for Rec8 expression dynamics might affect evolution of these loci^42^, which have been proposed to be precursors of sex chromosomes^43–45^ and to promote the evolution of anisogamy in isogamous ancestors^43,46–48^. Furthermore, the mating-type-biased selection pressures we observe might shape composition of populations, which could explain why P-cells may be underrepresented in the majority of fission yeast isolates^49^ and the dominance of one mating type observed in medically relevant fungal species^50,51^. By studying Rec8 physiological roles across biological scales — from molecular and cellular functions to population dynamics — this work bridges cell biology and evolutionary theory, offering a mechanistic explanation for how functional differences between gametes may evolve before the appearance of morphological dimorphism.

## SUPPLEMENTAL INFORMATION

**Supplemental Document S1** contains Supplemental Figures S1-S7 with accompanying Supplemental Figure Legends, Supplemental Movie Legends, Methods and Supplemental References sections

**Supplemental Notes S2** contains details of the numerical and statistical methods relevant to the computational model, theory and fits to experimental data.

**Supplemental Movies 1-12**.

**Supplemental Tables 1-4**.

Supplemental Table S1. Excel file contains the composition of media used in this study.

Supplemental Table S2. Excel file contains the list of strains used in this study.

Supplemental Table S3. Excel file contains the list of genetic markers used in this study and plasmids used to generate them.

Supplemental Table S4. Excel file contains the list of differentially expressed genes between mated and unmated fission yeast populations.

**Supplemental Sequences** of plasmids used in this study are available at Figshare doi.org/10.6084/m9.figshare.31555474 .

## RESOURCE AVAILABILITY

The code related to the theoretical analyses is available at the GitHub repository https://github.com/harrybooth/Fission-Yeast-Rec8. All plasmids and fission yeast strains are available upon reasonable request. All data is available upon reasonable request. All custom scripts used in experimental quantifications are available upon reasonable request.

## AUTHOR CONTRIBUTIONS

CM and AV designed the wetlab experiments. CM performed wetlab experimental work. CSC generated scripts for automated quantifications of imaging data. HB and ZH designed theoretical framework. HB implemented the theoretical model. CM, HB, ZH and AV and wrote the initial manuscript. ZH and AV revised the manuscript.

## ACKNOWLEDGMENTS

We thank Sophie Martin and Shiv Grewal for sharing published fission yeast genetic markers and plasmids. We thank Kabir Husain, Nadine Vastenhouw, Stephan Gruber and Richard Benton for critical reading of the manuscript. We thank Amy Bowen, Lewis Mosby, Michele Marconcini, Charles Mullon and Hadjivasiliou and Vjestica lab members for constructive discussions.

## FUNDING

This work has been funded by the Swiss National Science Foundation grant Eccellenza PCEFP3_187004/1, European Research Council grant 949914 ZygoticFate, and University of Lausanne funding obtained by AV. CSC obtained funding from the European Molecular Biology Organisation ALTF 89-2022. HB and ZH were supported by the Francis Crick Institute, which receives its core funding from Cancer Research UK, the UK Medical Research Council, and Wellcome Trust.

## DECLARATION OF INTERESTS

The authors declare no competing interests.

## DECLARATION OF GENERATIVE AI AND AI-ASSISTED TECHNOLOGIES

During the preparation of this manuscript, the authors used ChatGPT in order to improve text clarity and grammar. The authors subsequently reviewed and edited the content as needed and take full responsibility for the content of the manuscript.

## Supplemental Document S1

### The Supplemental Document S1 contains

1. Supplemental Figures S1-S7
2. Supplemental Figure Legends
3. Supplemental Movie Legends Note that Movies are available at: https://doi.org/10.6084/m9.figshare.31564093
4. Methods
5. Supplemental References.

## SUPPLEMENTAL FIGURE LEGENDS

**Figure S1.**
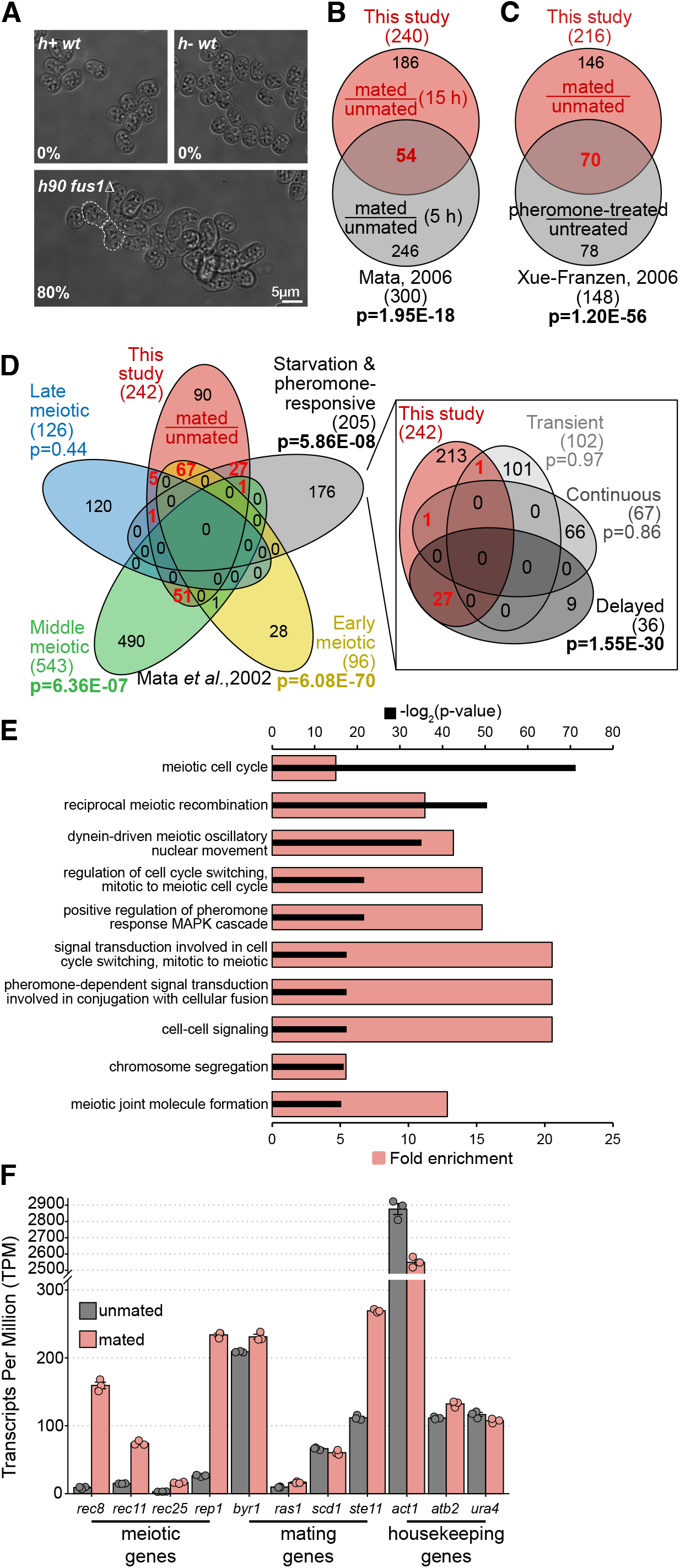
Fission yeast mating pairs upregulate meiotic transcripts ahead of fertilization. **(A)** Micrographs show brightfield images of fission yeast nitrogen-starved populations with indicated genotypes that we analysed by RNA sequencing. Heterothallic *h+* and *h-* strains produced only P- and M-gametes, respectively, and are thus unable to mate individually. Homothallic *h90* strains undergo mating-type switching to produce both gamete types. The *fus1Δ* mutants lack the formin necessary for partner fusion and thus partners arrest as mated pairs (dashed lines). The cells were incubated 15 h without nitrogen source. Percentages indicate mating efficiency. Scale bar is indicated. **(B-D)** We compared gene expression of mated and unmated fission yeast gametes shown in (A) and for significantly upregulated protein coding genes (| log_2_(fold-change)| ≥ 1, adjusted p-value ≤ 0.05) we report the overlap with earlier studies using Venn diagrams. Mated gametes showed 2-fold upregulation of 255 protein coding genes (**Fig.1A**), which included genes induced by partner presence^12^ (B), exogenous pheromone treatment^13^ (C) and meiotic genes^11^ (D). In **(B)** we show significant overlap with genes exposure using the data from Mata *et al*., 2006 (ref^12^, see Materials and methods). In **(C)** we show a strong overlap with genes induced upon treatment of gametes with exogenous pheromones, as reported by Xue-Franzen *et al*. 2006 (ref^13^). In **(D)** we used the data obtained by Mata *et al*., 2002 (ref^11^) from fission yeast diploid cells induced to synchronously enter meiosis. Briefly, diploid cells induced to enter meiosis show four temporal waves of gene inductions that coincided with major biological processes of nutritional adaptation and gamete differentiation (starvation- or pheromone-induced genes; gray areas), premeiotic S-phase and recombination (early meiotic genes; yellow areas), meiotic divisions (middle genes; green areas) and spore formation (late genes; blue areas). Note the highly significant overlap between early meiotic genes and genes we find upregulated during mating. When starvation- or pheromone-induced genes are further categorized according to their dynamics^11^ into “transient”, “continuous”, and “delayed” categories (D, inset), we observe a highly significant overlap between genes induced in our assay and the “delayed” pheromone-responsive genes. We provide hypergeometric test p-values for all analysed comparisons and report the number of genes in each category. Note that we used a variable number of upregulated genes (216-242) between comparisons because each previous study probed different sets of genes. **(E)** The barplot shows the top ten most significantly enriched Gene Ontology (GO) categories for protein coding genes upregulated in mated populations over unmated populations. The statistical significance (top axis, black bars) was obtained with the hypergeometric test using the fission yeast Uniprot protein database as the sample background. The fold-enrichment (bottom axis, red bars) reports over-representation of genes from indicated GO categories as compared to their frequency in the genome. Note the high significance and enrichment for GO categories relating to meiotic processes. **(F)** The graph reports the abundance of transcripts (TPM, transcripts per million) for indicated genes in non-mated (gray) and mated (pink) populations of gametes from RNAseq analyses. Note the comparable transcript levels under mating conditions between genes associated with meiotic recombination and chromosome segregation (*rec8, rec11, rec25* and *rep1*) and genes necessary for mating (*byr1, ras1, scd1* and *ste11*) and several house-keeping genes (*act1, atb2* and *ura4*). Bars report mean values, error bars denote standard deviation, dots indicated individual datapoints.

**Figure S2.**
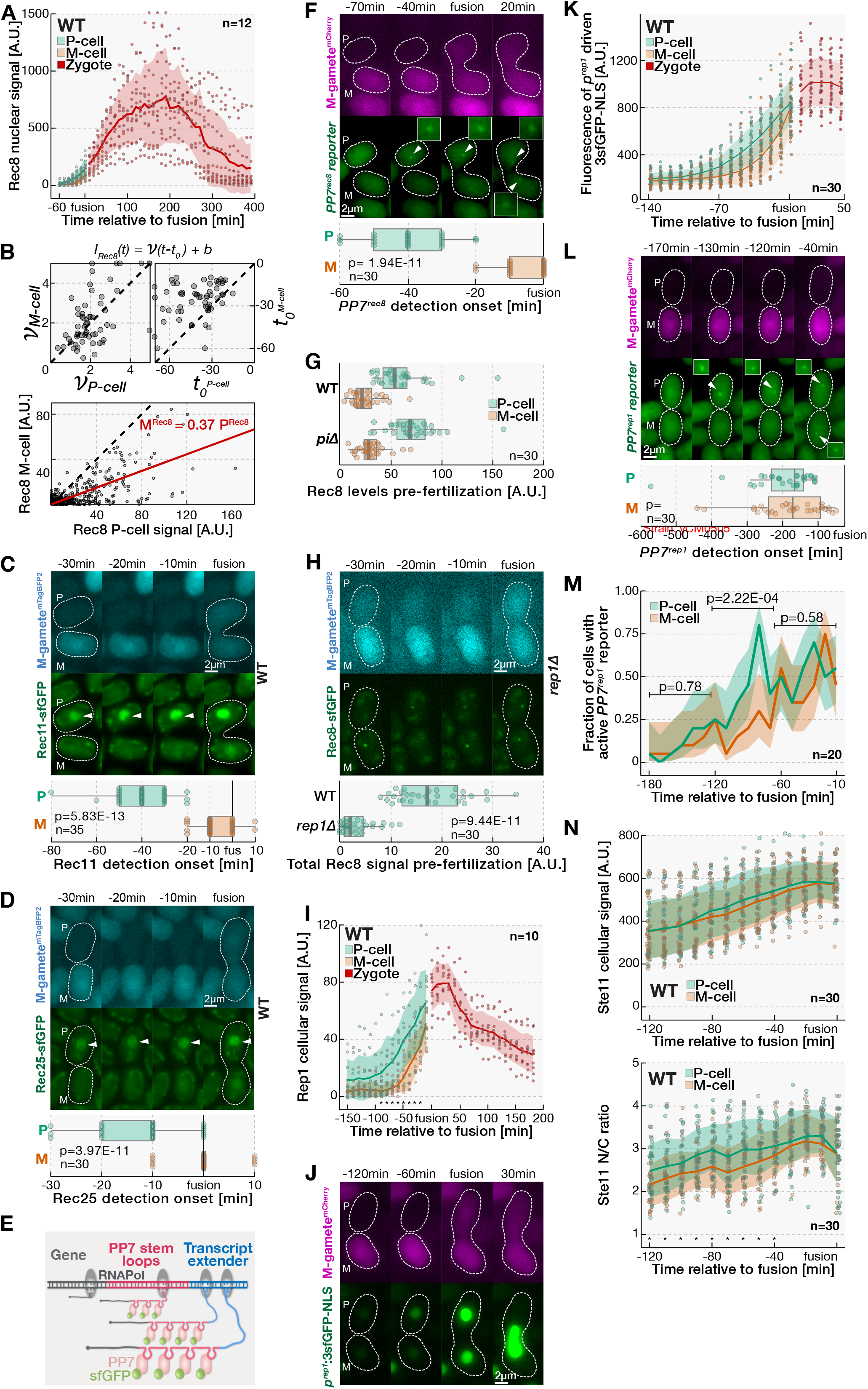
P-gametes are first to increase pheromone signaling and express of multiple meiotic genes. **(A)** The graph reports mean nuclear Rec8-sfGFP intensity throughout sexual reproduction. Note that Rec8-sfGFP is induced in gametes, that production continues in zygotes, with levels peaking approximately 3 h post-fusion, and that the Rec8-sfGFP signal is lost late in development as cells progress through meiotic divisions (**Mov.1**). **(B)** In top panels, dot plots show the rate (*v*) and time of production onset (t_0_) for Rec8-sfGFP production in P- and M-gametes for individual wild-type mating pairs. The Rec8-sfGFP signal intensity during mating were fitted using the indicated functional form where is the signal intensity at time, is the production rate and is the onset of production. The mean value across all fits for the P-gametes was, and for the M-gamete was . In the bottom panel, Rec8-signal in each partner from 60 to 10 min pre-fusion has been plotted and the best orthogonal regression fit, with an, is indicated with the red line. Dashed black lines indicate equivalence of parameters for P- and M-gametes. Note that partner gametes produce Rec8 at similar rates, and that the time of production onset differs. Also note that M-partners produce approximately a third of Rec8 detected in P-partners. **(C-D)** Green channel micrographs show Rec11-sfGFP (C) and Rec25-sfGFP (D) during mating in homothallic wild-type strains. Cells co-express the mTagBFP2 from the *p*^*mam1*^ promoter (blue), which we used to determine the mating type of gametes and the time of partner cells fusion. Boxplots report the onset of detection for the sfGFP tagged protein in P- and M-gametes. Note that Rec11-sfGFP and Rec25-sfGFP accumulate in the centrally positioned nucleus and that the expression is first detectable in P-gametes (arrowheads, boxplots). **(E)** The design of the PP7 reporter used to visualize transcription *in vivo*. We placed 24 PP7 stem-loops (red) after the STOP codon of the gene of interest (gray). We also introduced the 3.2kb fragment of the budding yeast GLT1 ORF and a transcriptional terminator (blue) to extend the transcript and increase its dwell time at the transcription site, which allows better signal detection. The cells co-express the sfGFP (green) fused with the non-aggregating PP7 viral protein variant (pink) that binds the RNA stem-loops. As RNA polymerase (gray ellipses) produces transcripts — each carrying 24 stem-loops — sfGFP-PP7 is concentrated to produce a fluorescent dot at the transcription site as a proxy of active transcription. **(F)** The micrographs show the PP7 reporter (green) detailed in (E) used to monitor *rec8* gene transcription. The M-gamete-specific *p*^*mam1*^ promoter regulates expression of mCherry (magenta), which we used to determine the gametes mating type and time of fusion. Boxplots report the onset of PP7 reporter dot (arrowheads) detection in P- and M-gametes. Note that the *rec8* transcription is detected first in P-gametes and subsequently in M-gametes. **(G)** Boxplots report the Rec8-sfGFP levels in P- and M-gametes 10 min prior to cell fusion for wild-type and *piΔ* mutant mating pairs, as indicated. Kruskal-Wallis test p-values comparing Rec8-sfGFP levels between P- and M-partners are 1.49E-08 and 4.00E-09 for wild-type and *piΔ* mating pairs, respectively. Note that the lack of Pi, which regulates the parental bias in expression of the *mei3* gene in zygotes^4^, does not regulate asymmetric Rec8 expression in gametes. **(H)** Micrographs shows Rec8-sfGFP (green) during mating of homothallic *rep1Δ* mutant strains. The mTagBFP2 (blue) as in (C). Boxplots report the Rec8-sfGFP levels in P-gametes 10 min prior to gamete fusion for wild-type and *rep1Δ* mutant mating pairs, as indicated. Note the severely reduced Rec8 production in *rep1Δ* mutants and that, only in some cells, Rec8-sfGFP forms faint nuclear foci, possibly marking the centromeres. **(I)** The graph reports mean cellular Rep1-sfGFP intensity throughout sexual reproduction. Note that Rep1-sfGFP is induced first in P-gametes and only subsequently in M-gametes. Rep1-sfGFP levels peak approximately 30 min post-fusion and the signal decreases as development progresses (**Mov.1**). **(J)** Micrographs show the 3sfGFP-NLS (green) expressed from the *p*^*rep1*^ promoter during wild-type matings. The mCherry (magenta), as in (F). Note the faint signal before cells mate, which suggests that the *p*^*rep1*^ promoter is “leaky” even before partners grow towards each other. Note that as partners mate, the green fluorescence is upregulated first in P-gametes and only subsequently in M-gametes. **(K)** The graph reports mean cellular intensity for 3sfGFP-NLS expressed from the *p*^*rep1*^ promoter shown in (J) throughout mating. **(L)** The micrographs show the PP7 reporter (green) detailed in (E) that we used to monitor *rep1* gene transcription. The mCherry (magenta) as in (F). Boxplots report the onset of PP7 reporter dot detection in P- and M-gametes. In mating populations, we observed transient bursts of *rep1* PP7 reporter activity in both P- and M-gametes even before mating took place, which is consistent with the observed “leakiness” of the *rep1* promoter (J-K). Importantly, as partner gametes engaged each other, we detected *rep1* transcription bursts more frequently in P-than M-gametes (M). In the final mating steps, *rep1* PP7 reporter activity bursts occurred with similar frequency in both partners (M). Consistent with the observed *rep1* promoter activity (J-K), our results show that the *rep1* transcription occurs more robustly in P-than M-gametes. **(M)** The graph reports mean detection frequency (lines) for the *rep1* transcription reporter shown in (L) in the 3 h before gamete fusion. Note significantly more frequent detection in P-gametes during 60-120 min before fertilization. Shaded areas report the Wilson interval and the p-values are from the exact McNemar’s test. **(N)** The graphs report the mean cellular intensity (top) and nucleocytoplasmic ratio (bottom) for Ste11-sfGFP in mating wild-type P- and M-gametes, as indicated. We used the paired t-test to compare the nucleocytoplasmic ratio in P- and M-gametes and report values lower than 0.05 with an asterisk (*). Note that the Ste11-sfGFP nuclear targeting is significantly higher in P-than M-gametes early in the mating process. All timepoints are relative to partner cell fusion and gamete mating types are indicated (P/M). Scale bars are indicated in micrographs, white dashed lines indicate cell boundaries visualized from brightfield images. Dots represent individual datapoints and we indicate the number of analysed mating events (n). Unless indicated otherwise, we show Kruskal-Wallis test p-values. We show either exact p-values or indicate p-value range with asterisks as 0.05 > *. In boxplots, box edges indicate 25th and 75th percentiles, the central line denotes the median, and whiskers extend 1.5 times the interquartile range. In line graphs, lines connect mean values and shaded areas report standard deviation.

**Figure S3.**
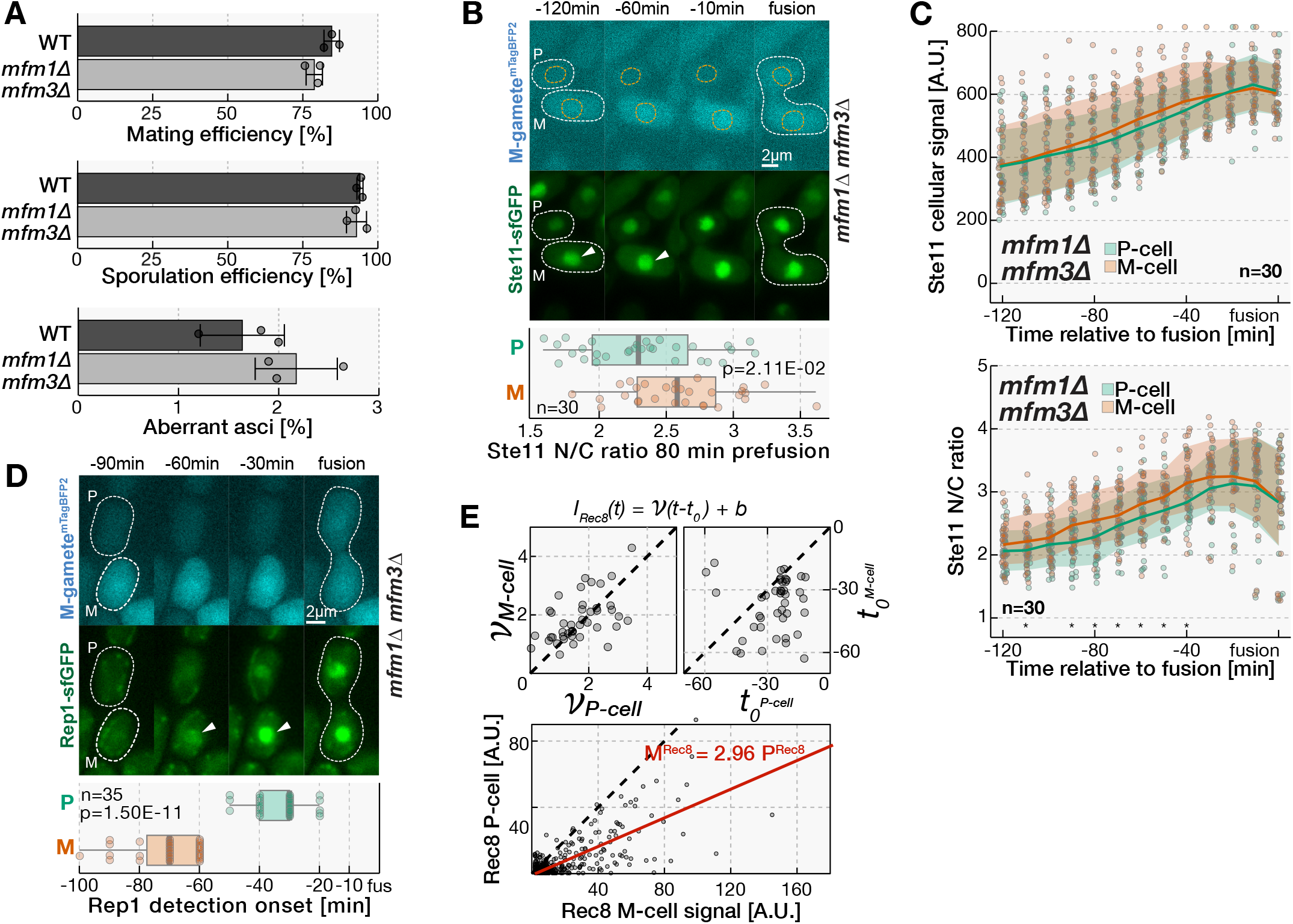
Asymmetries in the early expression of several meiotic genes in P-gametes are regulated by pheromone signalling. **(A)** As indicated, bar charts report mating and sporulation efficiencies, and aberrant sporulation rates in mating mixtures of either wild-type or *mfm1Δ mfm3Δ* mutant strains 48 h upon mating induction. Bars report mean values, error bars denote standard deviation, and dots indicated individual datapoints. Note that the *mfm1Δ mfm3Δ* mutants, which delay the onset of mating, show only minor defects in sexual reproduction at 48 h after mating induction. **(B)** Micrographs show Ste11-sfGFP (green) during mating of *mfm1Δ mfm3Δ* mutant partners. The mTagBFP2 (blue) as in (C). Nuclear boundaries, which we visualized with the nuclear ^NLS^mCherry marker (**Mov.2**), are indicated with orange dashed lines. Boxplots report the Ste11-sfGFP nucleocytoplasmic (N/C) ratio in P- and M-gametes 80 min before fusion. Note that M-gametes shows higher nuclear Ste11 targeting than the P-partners 80 min before partners fuse (arrowhead, boxplots). **(C)** The graphs report the mean cellular intensity (top) and nucleocytoplasmic ratio (bottom) for Ste11-sfGFP in mating *mfm1Δ mfm3Δ* P- and M-gametes, as indicated. We used the paired t-test to compare the nucleocytoplasmic ratio in P- and M-gametes and report values lower than 0.05 with an asterisk (*). Note that the Ste11-sfGFP nuclear targeting is significantly higher in M-than P-gametes early in the mating process. **(D)** Micrographs show Rep1-sfGFP (green) during mating of homothallic strains lacking the *mfm1* and *mfm3* genes. The mTagBFP2 (blue) as in (C). Boxplots report the onset of Rep1-sfGFP signal detection in P- and M-gametes. Note that Rep1-sfGFP signal is produced first in M-gametes (arrowheads, boxplots) during mating of *mfm1Δ mfm3Δ* partners. In top panels, dot plots show the rate (*v*) and time of production onset (t_0_) for Rec8-sfGFP production in P- and M-gametes for individual *mfm1Δ mfm3Δ* mating pairs. The Rec8-sfGFP signal traces were fitted as in (B). The mean value across all fits for the P-gametes was, and for the M-gamete was . **(E)** In the bottom panel, Rec8-signal in each partner from 60 to 10 min pre-fusion has been plotted and the best orthogonal regression fit, with an, is indicated with the red line. Note that partner gametes produce Rec8 at similar rates but first in M-cells. Note that the *mfm1Δ mfm3Δ* M-gametes produce approximately 3-fold more Rec8 than the M-gametes. All timepoints are relative to partner cell fusion and gamete mating types are indicated (P/M). Scale bars are indicated in micrographs, white dashed lines indicate cell boundaries visualized from brightfield images. Dots represent individual datapoints and we indicate the number of analysed mating events (n). Unless indicated otherwise, we show Kruskal-Wallis test p-values. We show either exact p-values or indicate p-value range with asterisks as 0.05 > *. In boxplots, box edges indicate 25th and 75th percentiles, the central line denotes the median, and whiskers extend 1.5 times the interquartile range. In line graphs, lines connect mean values and shaded areas report standard deviation.

**Figure S4.**
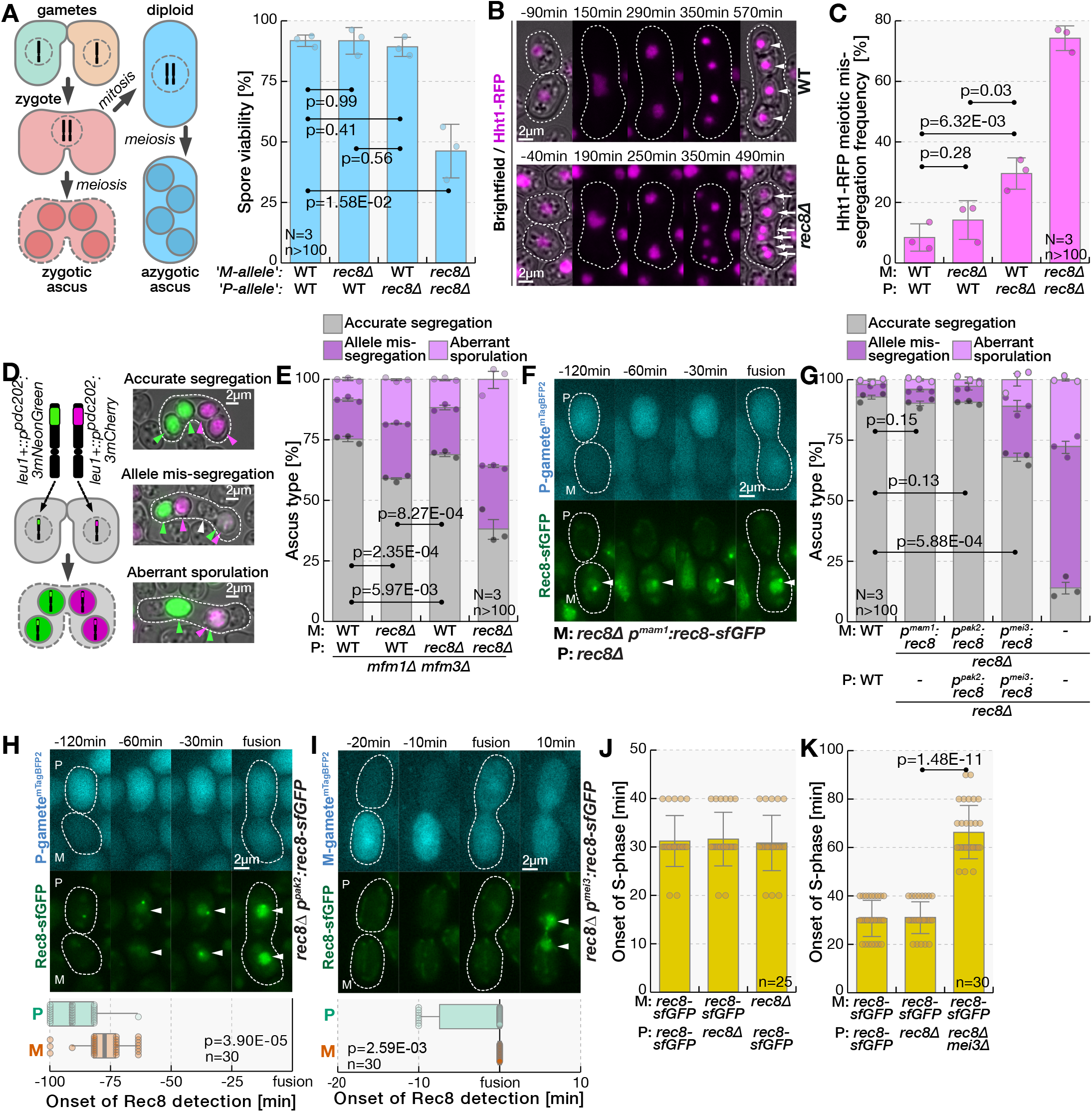
Rec8 is haplo-sufficient in diploids and its expression in either gamete ensures accurate meiosis in zygotes. **(A)** The schematic (left) illustrates zygotic and azygotic asci formation. Haploid gametes fuse to produce a zygote, which enter meiosis to produce zygotic asci (red). Rare zygotes that do not immediately enter meiosis can be selected and propagated mitotically as diploid cells in nitrogen-rich conditions (blue). Upon nitrogen depletion, diploids undergo meiosis and sporulation to produce azygotic asci (see Materials and methods). The bar chart (right) reports the viability of spores produced in azygotic asci^42^ of diploid strains with indicated genotypes. Even though differences between parental alleles have not been reported for diploid cells, or during azygotic meiosis, we indicate the origin of *rec8* alleles in diploid cells. Note that deleting both *rec8* copies severely reduces viability of spores produced by diploid cells but that removal of one *rec8* allele does not significantly reduce spore viability. **(B)** Timelapses show localization of the Hht1 histone H3^23^ fused to the red fluorescent protein RFP (magenta) during mating of either wild-type or *rec8Δ* mutants, as indicated. The brightfield images are overlayed in the first and last timepoints. We did not observe defects in fusion of gametic nuclei in wild-type or zygotes produced by *rec8* mutant gametes (0/100 zygotes for each cross, **Mov.3**). Meiotic divisions produced four equal chromatin masses of Hht1-RFP in majority of zygotes produced by two wild-type parents (arrowheads). Zygotes with two *rec8Δ* parents often segregated Hht1-RFP into aberrant number of uneven chromatin masses (arrows). **(C)** The bar chart reports the frequency of Hht1-RFP meiotic segregation defects described in (B) in zygotes produced by indicated crosses of heterothallic wild-type and *rec8Δ* mutant strains. Note that deleting *rec8* only in P-gametes causes more pronounced chromatin segregation defects than deleting *rec8* only in M-gametes. **(D)** The schematic shows the design of synthetic reporter alleles used to monitor meiotic allele segregation. The *leu1+* genomic locus on chromosome II was used to integrate the sporulation-induced promoter *p*^*pdc202*^ followed by three tandem repeats of fluorescent proteins mCherry, in one parent, or mNeonGreen^7^, in the other parent. Crosses between the two parents produces spores with fluorescence that indicates the inherited alleles. Micrographs show the different types of asci produced by crossing otherwise wild-type strains that carry the two reporter alleles. We show the overlay of brightfield with green and red fluorescent channels (gray, green and magenta, respectively). Green and magenta arrowheads point to spores emitting corresponding fluorescent signals, and white arrowheads indicate spores that do not produce fluorescence. We show asci that accurately segregate alleles between four spores (top; 91±1%), form four spores but mis-segregate reporter alleles (middle, note the spore with red and green fluorescent signal; 6±1%), or form fewer than four spores (bottom; 3±1%). **(E)** Stacked histograms shown the frequency of asci shown in (D) and observed in indicated crosses of heterothallic *mfm1Δ mfm3Δ* mutant strains where the *rec8* gene is either intact or deleted. The strains carry the reporter alleles detailed in (D). Note that in *mfm1Δ mfm3Δ* mutants, deleting *rec8* only in M-gametes causes more pronounced meiotic segregation defects than deleting *rec8* in P-gametes only. Note that deleting *rec8* in both *mfm1Δ mfm3Δ* partners strongly impairs allele segregation during meiosis. Welch test p-values are shown. **(F)** Micrographs show Rec8-sfGFP (green) expressed from the M-cell specific *p*^*mam1*^ promoter during mating of partners that lack the native *rec8* gene. P-gametes utilize the P-cell-specific *p*^*map3*^ promoter to express mTagBFP2 (blue), which we used to determine the mating type of gametes and time of partner fusion. Dashed lines indicate cell boundaries visualized from brightfield images. We did not quantify the time of Rec8-sfGFP induction relative to cell fusion since M-cells activate the *p*^*mam1*^ promoter already at the start of the experiment (arrowheads), which is not correlated to the time of cell fusion. **(G)** Stacked histograms show frequency of asci shown in (D) in mating mixture produced by wild-type gametes and crosses where Rec8 is either lacking or solely expressed from the M-gamete-specific *p*^*mam1*^ promoter, the mating-induced *p*^*pak2*^ promoter or zygotic *p*^*mei3*^ promoter, as indicated. Additionally, strains carry the reporter alleles detailed in (D). Note the largely accurate meiotic allele segregation in crosses where *p*^*mam1*^ and *p*^*pak2*^ promoters drive Rec8 expression in gametes, and high incidence of meiotic defects when Rec8 is expressed at cell fusion from the *p*^*mei3*^ promoter. Welch test p-values are shown. **(H-I)** Micrographs show Rec8-sfGFP (green) expressed from the mating-induced *p*^*pak2*^ promoter (H) and the zygotic *p*^*mei3*^ promoter (I) during mating of homothallic strains where native *rec8* alleles are deleted. The M-gamete-specific *p*^*mam1*^ promoter regulates expression of mTagBFP2 (blue), which we used to determine the gametes mating type and the time of fusion. Boxplots report the onset of Rec8-sfGFP detection in P- and M-gametes. Kruskal-Wallis test p-values are shown. Note that Rec8-sfGFP expression from the *p*^*pak2*^ promoter starts earlier, and the expression from the *p*^*mei3*^ promoter later, than the wild-type Rec8 expression (**Fig.1C**). **(J-K)** Bar charts report the time from partner fusion until the S-phase is triggered in indicated crosses between heterothallic strains where the native *rec8* gene is either deleted or fused to the sfGFP sequence. Additionally, we indicate the strain that carries the *mei3Δ* mutation (K). Kruskal-Wallis test p-value for indicated comparison is shown. Note that mutating *rec8* gene in either parent does not affect the time of pre-meiotic S-phase onset in zygotes (J). Also note that that deleting the *mei3* P-allele significantly delays premeiotic S-phase initiation in zygotes (K). All timepoints are relative to partner cell fusion and gamete mating types are indicated (P/M). Scale bars are indicated in micrographs, white dashed lines indicate cell boundaries visualized from brightfield images. Dots represent individual datapoints and we indicate the number of analysed replicates (N) and relevant cell types (n). In bar charts and stacked histograms, bars report means and error bars denote standard deviation. In boxplots, box edges indicate 25th and 75th percentiles, the central line denotes the median, and whiskers extend 1.5 times the interquartile range. We report exact p-values from statistics tests reported detailed in panel legends.

**Figure S5.**
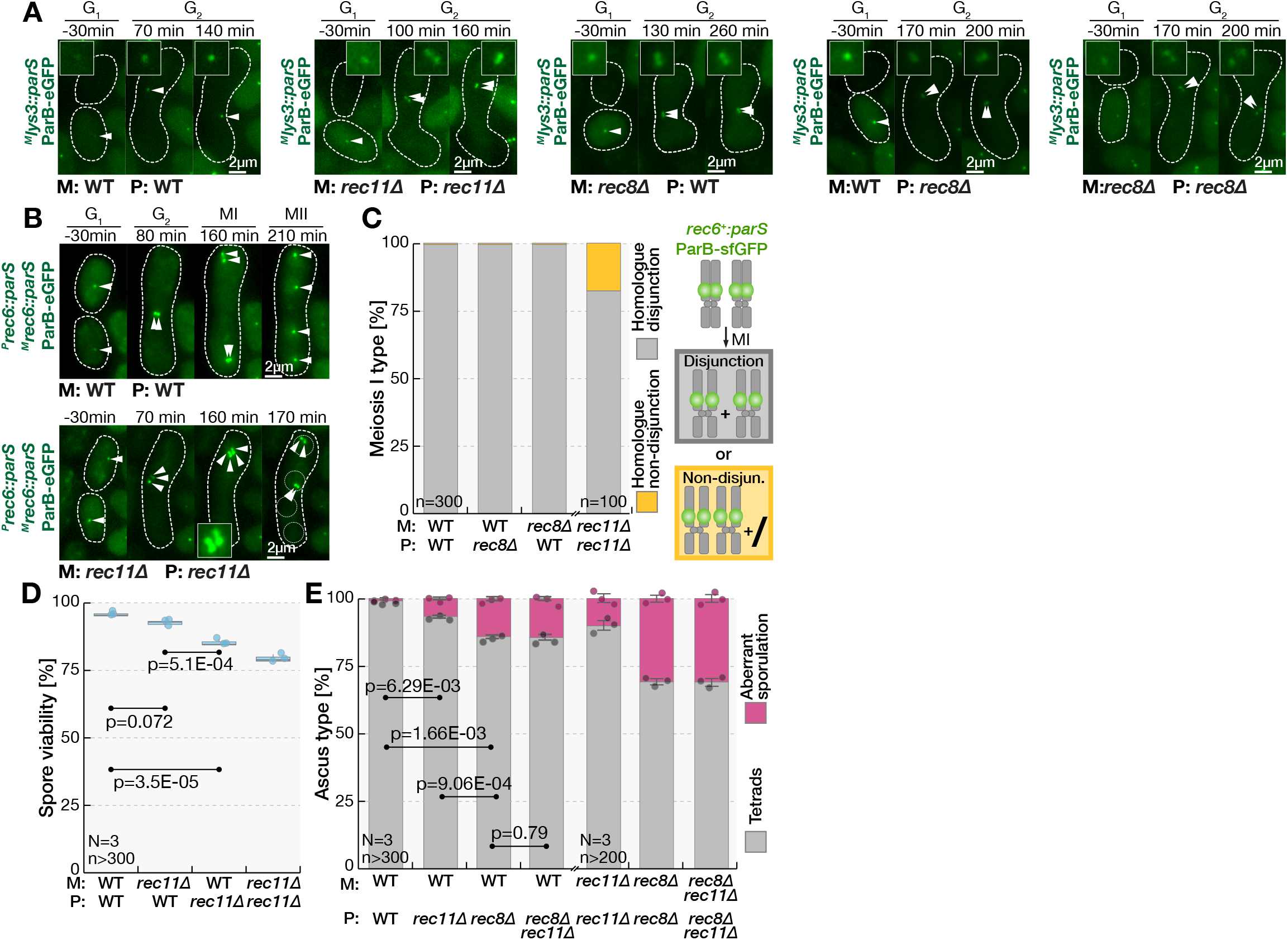
The *rec8* P-allele contributes more to meiotic success than the M-allele primarily by promoting meiotic centromere function and reductional meiosis I. **(A)** Micrographs show mating between indicated wild-type, *rec8Δ* and *rec11Δ* partners where M-gametes carry the *lys3* locus labeled with the *parS*^*c3*^/ParB^c3^-eGFP (green) and P-gametes carry the *lys3* locus labeled with the *parS*^*c2*^/ParB^c2^-mCherry-NLS (see **Mov.7**). Cell cycle phases are indicated (G_1_, G_2_). Note that the *lys3* locus-recruited *parS*^*c3*^/ParB^c3^-eGFP (arrowheads, insets) produces a single dot prior to meiosis in wild-type zygotes but two dots in zygotes that lack either parental *rec8* allele. **(B)** Micrographs show mating between indicated wild-type, *rec11Δ* and *rec8Δ* partners where both gametes carry the *rec6* locus labeled with the *parS*^*c3*^/ParB^c3^-eGFP (green). Cell cycle phases are indicated (G_1_, G_2_, MI and post-MII). Note that during meiosis I, in wild-type zygotes the pairs of *rec6* loci segregate to opposite poles (homologue disjunction) and that in *rec8* mutants the four *rec6* loci co-segregate together (homologue non-disjunction, see insets). **(C)** For indicated crosses between wild-type and *rec8Δ* mutants, stacked histograms report the frequency of phenotypes shown in (B), where during meiosis I parental *rec6* loci segregate as pairs to opposite poles (homologue disjunction) or co-segregate all together (homologue non-disjunction). **(D)** Boxplots report the viability of spores produced in indicated crosses of wild-type and *rec11Δ* mutant gametes. Tukey test p-values are shown. Note that deleting *rec11* in the P-gamete reduces spore viability more than deleting it in the M-gamete. **(E)** Stacked histograms report the frequency of tetrad asci and asci containing aberrant spore number for indicated crosses between wild-type, *rec11Δ* and *rec8Δ* mutant gametes. Welch test p-values are shown. Note that meiotic defects are more pronounced upon deleting *rec8* than *rec11*, and that *rec8Δ* is epistatic to *rec11Δ*. All timepoints are relative to cell fusion. Scale bars are indicated in micrographs, white dashed lines indicate cell boundaries visualized from brightfield images. Dots represent individual datapoints and we indicate the number of analyzed replicates (N) and relevant cell types (n). In stacked histograms, bars report means and error bars denote standard deviation. In boxplots, box edges indicate 25th and 75th percentiles, the central line denotes the median, and whiskers extend 1.5 times the interquartile range.

**Figure S6.**
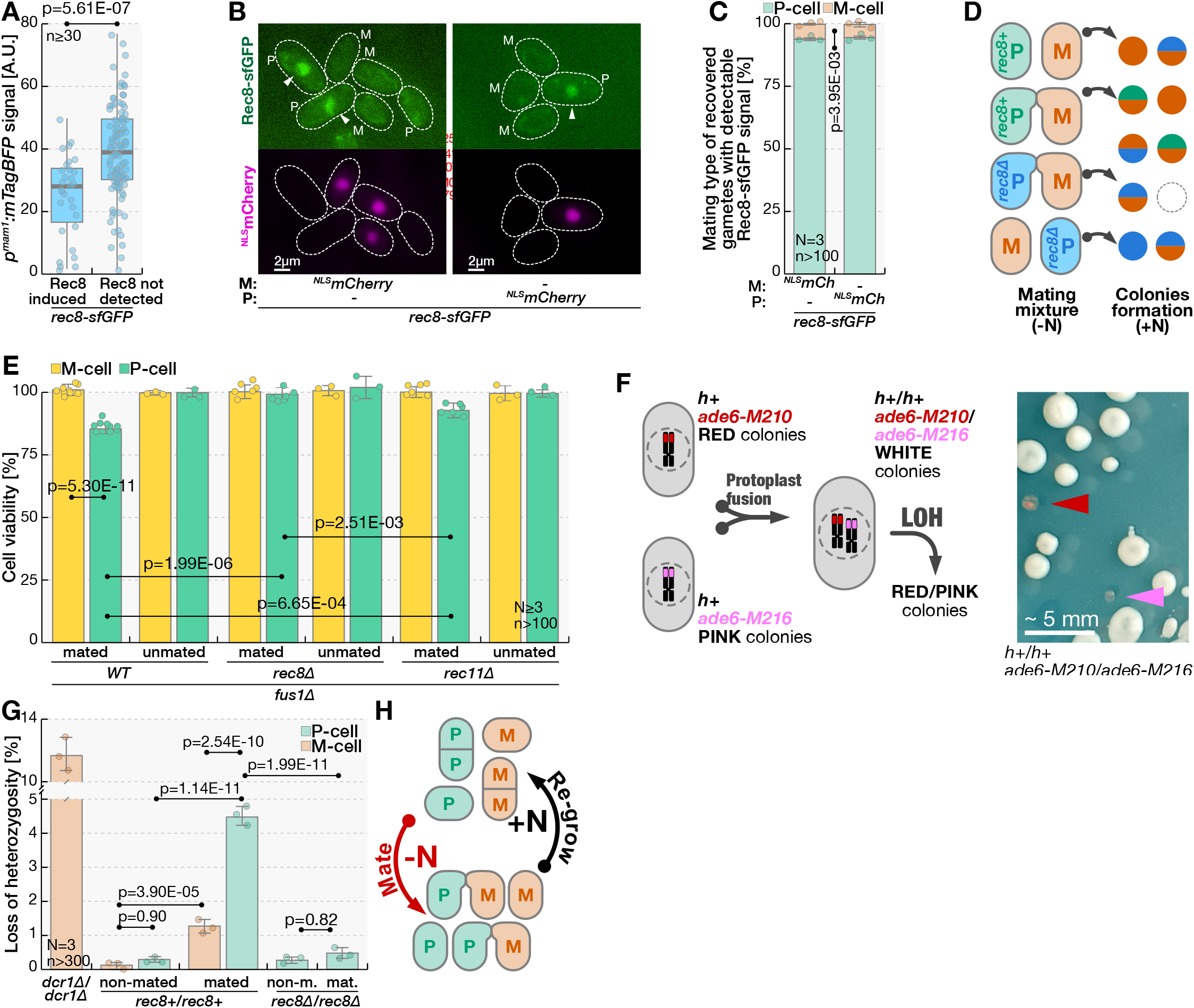
Mating produces a population of Rec8-expressing P-gametes and increases genomic instabilities in P-gametes that return to mitotic proliferation. **(A)** Homothallic cells coding for Rec8-sfGFP and expressing mTagBFP2 from the M-cell-specific *p*^*mam1*^ promoter, were induced to mate and the mating was interrupted by shifting cells to growth media. Boxplots report mean mTagBFP2 cellular fluorescence for gametes that either induce or lack detectable Rec8 signal at the time mating is interrupted. The Kruskal-Wallis test p-value is shown. Note that the gametes with detectable Rec8-sfGFP signal have lower mTagBFP2 levels and are thus likely enriched in P-gametes. Note that mTagBFP2 signal can be reliably used to determine the mating type of gametes only within a mating pair (*e*.*g*. in **Fig.1F** partners show small difference in mTagBFP2 levels). **(B)** Micrographs show Rec8-sfGFP (green) and ^NLS^mCherry (magenta) signals upon transferring the indicated heterothallic mating mixtures to growth media. We used cell morphology to distinguish gametes from zygotes, and the ^NLS^mCherry signal to determine the mating type of gametes (P, M). Note the Rec8-sfGFP in P-gametes (arrowheads). Dashed lines indicate cell boundaries visualized from brightfield images. **(C)** Stacked histogram shows the frequency of P- and M-mating type for gametes that induced Rec8 upon retrieval from indicated heterothallic mating mixtures onto growth media, as shown in (B). The Kruskal-Wallis test p-value reports the significance for the over-representation of the P mating-type in unfused Rec8-expressing gametes. Note that the Rec8-expressing gametes are predominantly the P-mating type. **(D)** The schematic illustrates the viability assay for mated and unmated gametes from mating mixtures. Since fission yeast mating is asynchronous, we used a microscopic needle to isolate individual gametes or mating pairs produced between heterothallic *fus1Δ* mutant P-gametes (green, blue) and M-gametes (orange) and transferred them onto growth media to assess their ability to form colonies. To ensure that *rec8+* and r*ec8Δ* P-gametes (green and blue, respectively) experience identical mating conditions, we pooled the two strains together before introducing the M-cells. Distinct selection markers were used to determine the colony progenitors, and we report gamete viability as the ratio between observed and expected number of colonies forming. We used the same approach to determine gamete viability in mating mixtures of *rec8+* and r*ec8Δ* M-gametes pooled with a single P-partner strain. **(E)** The bar chart reports viability of P- and M-cells from fission yeast mating mixtures with indicated genotypes. Tukey test p-values are shown. Note that mating reduces viability of P-cells, and that this is largely dependent on the *rec8* gene and partially dependent on the *rec11* gene. **(F)** The schematic (left) shows the design of the loss-of-heterozygosity (LOH) assay^8^, which detects both copy number loss and copy number-neutral allele loss. Briefly, because only *h+/h-* diploids acquire zygotic identity, the *h+/h+* and *h-/h-* diploids are stable and able to mate. We obtained the diploids by fusing protoplasts of haploid cells that carry *ade6-M210* (red) and *ade6-M216* (pink) mutations, which are individually auxotrophic but capable of interallelic complementation that restores adenine prototrophy in diploid cells. Loss of either *ade6* allele results in auxotrophic cells. Note that adenine prototrophs produce white colonies whereas adenine auxotrophs results in formation of colonies with red/pink hue that are easily visualized (arrowheads) as shown in the photograph (right). We deleted both alleles of the *fus1* gene to ensure that upon mating *h+/h+* and *h-/h-* diploids are unable to fuse. **(G)** The bar chat shows LOH rates for *ade6* locus in mated and non-mated strains with indicated genotypes, which we measured using the assay detailed in (F). We report the results for *fus1Δ/fus1Δ* mutant *h+/h+* and *h-/h-* diploids that were starved for nitrogen either in the presence or the absence of the mating partner before assessing the LOH rates. The *dcr1Δ/dcr1Δ* mutant was used as a positive control for the LOH assay^8^. Tukey test p-values are shown. Note that LOH rates increase in mating populations, and that the effect is significantly greater in P-than M-gametes. Also note that deletion of *rec8* reduces LOH rates in mated P-gametes. **(H)** The schematic shows the experimental design used to evolve fission yeast populations of P- and M-gametes in **Fig.3E**. We subjected populations of 10^7^ cells initially containing the same number of heterothallic P- and M-gametes to cycles of mating and regrowth. Note that partners could not fuse because we deleted the *fus1* gene. We performed the experiment using P- and M-gametes that either both carried the wild-type *rec8* gene or both lacked the *rec8* gene. Scale bars are shown for micrographs, white dashed lines indicate cell boundaries visualized from brightfield images and gamete mating types are indicated (P/M). Dots represent individual datapoints and we indicate the number of analyzed replicates (N) and relevant cell types (n). In boxplots, the top and bottom box edges indicate 25th and 75th percentiles, and the central line displays the median. The whiskers extend 1.5 times the interquartile range. Dots represent individual datapoints. In bar charts and stacked histograms, bars report means and error bars denote standard deviation.

**Figure S7.**
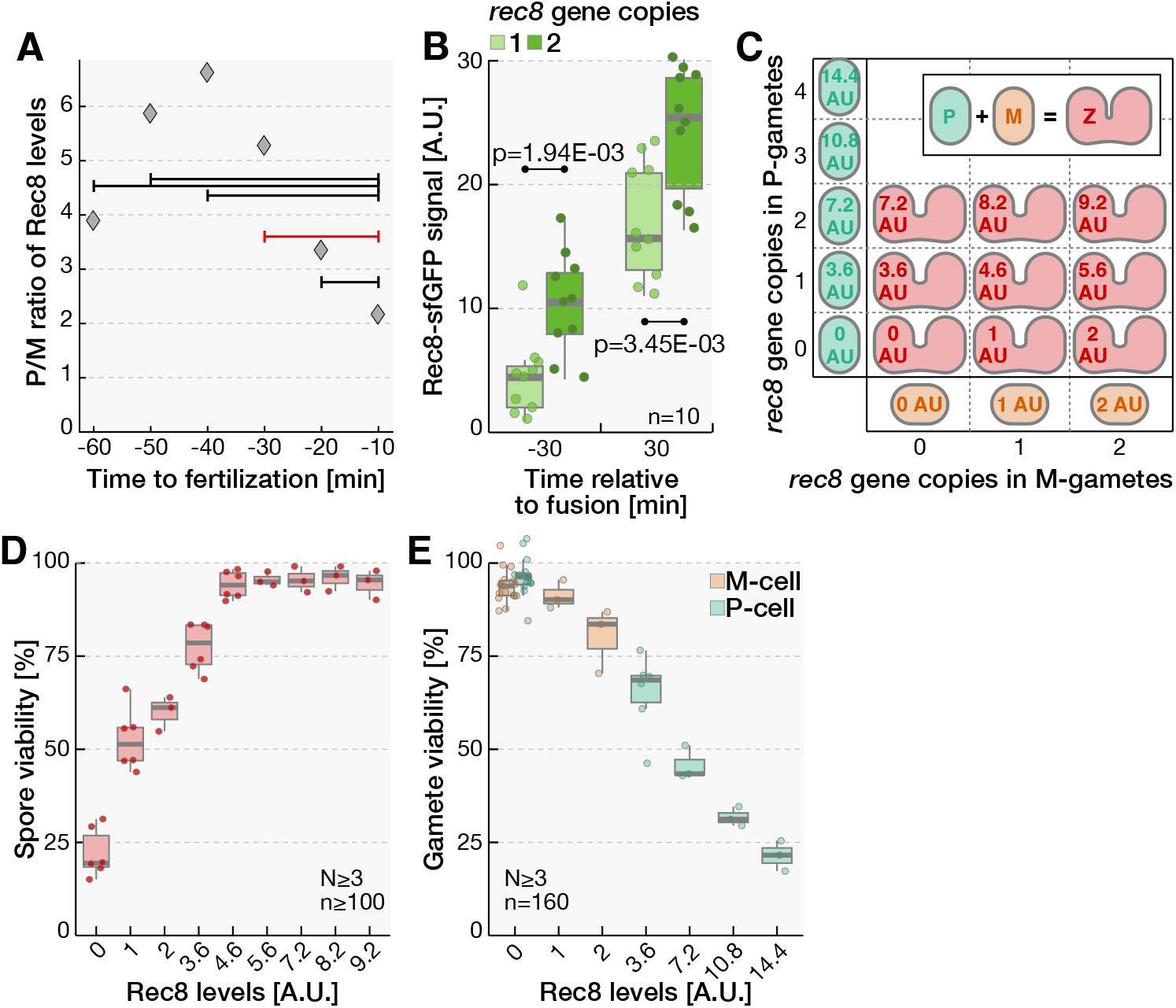
Quantifications and fits of zygotic success and spore viability as a function of Rec8 levels. **(A)** The plot reports the observed Rec8-sfGFP signal ratio between wild-type P- and M-gametes at indicated timepoints (diamonds), with intervals (bars) indicating the average of signal ratios calculated for timepoints in the interval pre-fusion. Each time point ratio was calculated using orthogonal regression using intensity data associated with indicated timepoints. For modeling, we relied on the P/M ratio of 3.6 in Rec8 levels, which is the average of the observed ratios calculated at -10, -20 and -30 minutes (red bar). We do not to include earlier timepoints because very low Rec8 expression in M-cells causes high signal-to-noise ratio and hence inflates the asymmetry. **(B)** Boxplots report mean Rec8-sfGFP fluorescence at indicated timepoints in crosses of gametes that carry the native *rec8* gene tagged with sfGFP (1 copy) or gametes that carry an additional copy of *rec8* gene tagged with sfGFP (2 copies). Note that the presence of two *rec8-sfGFP* gene copies approximately doubles the Rec8-sfGFP levels in gametes and zygotes. **(C)** The schematic illustrates how the levels of Rec8 in gametes and zygotes were manipulated to determine the and dependence on Rec8 levels. We normalized the Rec8 levels detected in wild-type M-gametes ahead of partner fusion to 1 A.U. and used the relative levels of Rec8 in wild-type P-gametes of 3.6 A.U., as detailed in (A). Introducing each additional *rec8* gene copy, as indicated on the axes, was assumed to each provide one fold increase of the Rec8 protein, based on results obtained in (B). Thus, each additional *rec8* gene copy in M- and P-gametes was assumed to increase the Rec8 levels by 1 and 3.6 A.U., respectively. We considered Rec8 expressed in each gamete as functionally equal when evaluating the effect of Rec8 level on gamete survival. Likewise, we considered Rec8 supplied by each parent functionally equal for spore viability, and, as indicated, we mated gametes with different *rec8* gene copy number to evaluate how total parental Rec8 levels affect zygote fitness. **(D)** The experimental data used to fit the probability of successful meiosis as a function of total meiotic factor investment . The boxplot reports viability of spores produced by crosses of heterothallic partners that supply indicated levels of Rec8, which are noted in A.U. as detailed in (C). Heterothallic gamete populations were allowed to mate and sporulate, before collecting a defined number of spores and plating them onto growth media to assess their ability to form colonies. **(E)** The experimental data used to fit the probability of successful return to mitosis as a function of meiotic factor produced in unfused gametes. The boxplot reports viability of mated gametes expressing indicated levels of Rec8, which are noted in A.U. as detailed in (C). Gamete populations were allowed to mate and to form mating pairs but could not fuse due to deletion of the *fus1* gene. Individual mating pairs were then transferred onto growth media and the ability to form colonies was assessed. In boxplots, top and bottom box edges indicate 25th and 75th percentiles, and the central line displays the median. The whiskers extend 1.5 times the interquartile range. Dots represent individual datapoints and we indicate the number of analyzed replicates (N) and relevant cell types (n).

**Figure S1. Fission yeast mating pairs upregulate meiotic transcripts ahead of partner fusion**

**(A)** Micrographs show brightfield images of fission yeast nitrogen-starved populations with indicated genotypes that we analysed by RNA sequencing. Heterothallic *h+* and *h-* strains produced only P- and M-gametes, respectively, and are thus unable to mate individually. Homothallic *h90* strains undergo mating-type switching to produce both gamete types. The *fus1Δ* mutants lack the formin necessary for partner fusion and thus partners arrest as mated pairs (dashed lines). The cells were incubated 15 h without nitrogen source. Percentages indicate mating efficiency. Scale bar is indicated.

**(B-D)** We compared gene expression of mated and unmated fission yeast gametes shown in (A) and for significantly upregulated protein coding genes (|log_2_(fold-change)| ≥ 1, adjusted p-value ≤ 0.05) we report the overlap with earlier studies using Venn diagrams. Mated gametes showed 2-fold upregulation of 255 protein coding genes (**Fig.1A**), which included genes induced by partner presence^1^ (B), exogenous pheromone treatment^2^ (C) and meiotic genes^3^ (D). In **(B)** we show significant overlap with genes upregulated within the first five hours of partner exposure using the data from Mata *et al*., 2006 (ref.^1^, see Methods). In **(C)** we show a strong overlap with genes induced upon treatment of gametes with exogenous pheromones, as reported by Xue-Franzen *et al*. 2006 (ref.^2^). In **(D)** we used the data obtained by Mata *et al*., 2002 (ref.^3^) from fission yeast diploid cells induced to synchronously enter meiosis. Briefly, diploid cells induced to enter meiosis show four temporal waves of gene inductions that coincided with major biological processes of nutritional adaptation and gamete differentiation (starvation- or pheromone-induced genes; gray areas), premeiotic S-phase and recombination (early meiotic genes; yellow areas), meiotic divisions (middle genes; green areas) and spore formation (late genes; blue areas). Note the highly significant overlap between early meiotic genes and genes we find upregulated during mating. When starvation- or pheromone-induced genes are further categorized according to their dynamics^3^ into “transient”, “continuous”, and “delayed” categories (D, inset), we observe a highly significant overlap between genes induced in our assay and the “delayed” pheromone-responsive genes. We provide hypergeometric test p-values for all analysed comparisons and report the number of genes in each category. Note that we used a variable number of upregulated genes (216-242) between comparisons because each previous study probed different sets of genes.

(**E**) The barplot shows the top ten most significantly enriched Gene Ontology (GO) categories for protein coding genes upregulated in mated populations over unmated populations. The statistical significance (top axis, black bars) was obtained with the hypergeometric test using the fission yeast Uniprot protein database as the sample background. The fold-enrichment (bottom axis, red bars) reports over-representation of genes from indicated GO categories as compared to their frequency in the genome. Note the high significance and enrichment for GO categories relating to meiotic processes.

(**F)** The graph reports the abundance of transcripts (TPM, transcripts per million) for indicated genes in non-mated (gray) and mated (pink) populations of gametes from RNAseq analyses. Note the comparable transcript levels under mating conditions between genes associated with meiotic recombination and chromosome segregation (*rec8, rec11, rec25* and *rep1*) and genes necessary for mating (*byr1, ras1, scd1* and *ste11*) and several house-keeping genes (*act1, atb2* and *ura4*). Bars report mean values, error bars denote standard deviation, dots indicated individual datapoints.

**Figure S2. P-gametes are first to increase pheromone signaling and express of multiple meiotic genes**

**(A)** The graph reports mean nuclear Rec8-sfGFP intensity throughout sexual reproduction. Note that Rec8-sfGFP is induced in gametes, that production continues in zygotes, with levels peaking approximately 3 h post-fusion, and that the Rec8-sfGFP signal is lost late in development as cells progress through meiotic divisions (**Mov.1**).

**(B)** In top panels, dot plots show the rate (*v*) and time of production onset (t_0_) for Rec8-sfGFP production in P- and M-gametes for individual wild-type mating pairs. The Rec8-sfGFP signal intensity during mating were fitted using the indicated functional form *I*_*Rec*8_(*t*) = *v* ∗ (*t* − *t*_0_) + *b* where *I*_*Rec*8_(*t*) is the signal intensity at time *t, v* is the production rate and *t*_0_ is the onset of production. The mean 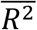, value across all fits for the P-gametes was 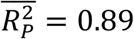, and for the M-gamete was 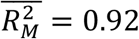. In the bottom panel, Rec8-signal in each partner from 60 to 10 min pre-fusion has been plotted and the best orthogonal regression fit, with an *R*^2^ = 0.42, is indicated with the red line. Dashed black lines indicate equivalence of parameters for P- and M-gametes. Note that partner gametes produce Rec8 at similar rates, and that the time of production onset differs. Also note that M-partners produce approximately a third of Rec8 detected in P-partners.

**(C-D)** Green channel micrographs show Rec11-sfGFP (C) and Rec25-sfGFP (D) during mating in homothallic wild-type strains. Cells co-express the mTagBFP2 from the *p*^*mam1*^ promoter (blue), which we used to determine the mating type of gametes and the time of partner cells fusion. Boxplots report the onset of detection for the sfGFP tagged protein in P- and M-gametes. Note that Rec11-sfGFP and Rec25-sfGFP accumulate in the centrally positioned nucleus and that the expression is first detectable in P-gametes (arrowheads, boxplots).

**(E)** The design of the PP7 reporter used to visualize transcription *in vivo*. We placed 24 PP7 stem-loops (red) after the STOP codon of the gene of interest (gray). We also introduced the 3.2kb fragment of the budding yeast GLT1 ORF and a transcriptional terminator (blue) to extend the transcript and increase its dwell time at the transcription site, which allows better signal detection. The cells co-express the sfGFP (green) fused with the non-aggregating PP7 viral protein variant (pink) that binds the RNA stem-loops. As RNA polymerase (gray ellipses) produces transcripts — each carrying 24 stem-loops — sfGFP-PP7 is concentrated to produce a fluorescent dot at the transcription site as a proxy of active transcription.

**(F)** The micrographs show the PP7 reporter (green) detailed in (E) used to monitor *rec8* gene transcription. The M-gamete-specific *p*^*mam1*^ promoter regulates expression of mCherry (magenta), which we used to determine the gametes mating type and time of fusion. Boxplots report the onset of PP7 reporter dot (arrowheads) detection in P- and M-gametes. Note that the *rec8* transcription is detected first in P-gametes and subsequently in M-gametes.

**(G)** Boxplots report the Rec8-sfGFP levels in P- and M-gametes 10 min prior to cell fusion for wild-type and *piΔ* mutant mating pairs, as indicated. Kruskal-Wallis test p-values comparing Rec8-sfGFP levels between P- and M-partners are 1.49E-08 and 4.00E-09 for wild-type and *piΔ* mating pairs, respectively. Note that the lack of Pi, which regulates the parental bias in expression of the *mei3* gene in zygotes^4^, does not regulate asymmetric Rec8 expression in gametes.

**(H)** Micrographs shows Rec8-sfGFP (green) during mating of homothallic *rep1Δ* mutant strains. The mTagBFP2 (blue) as in (C). Boxplots report the Rec8-sfGFP levels in P-gametes 10 min prior to gamete fusion for wild-type and *rep1Δ* mutant mating pairs, as indicated. Note the severely reduced Rec8 production in *rep1Δ* mutants and that, only in some cells, Rec8-sfGFP forms faint nuclear foci, possibly marking the centromeres.

**(I)** The graph reports mean cellular Rep1-sfGFP intensity throughout sexual reproduction. Note that Rep1-sfGFP is induced first in P-gametes and only subsequently in M-gametes. Rep1-sfGFP levels peak approximately 30 min post-fusion and the signal decreases as development progresses (**Mov.1**).

**(J)** Micrographs show the 3sfGFP-NLS (green) expressed from the *p*^*rep1*^ promoter during wild-type matings. The mCherry (magenta), as in (F). Note the faint signal before cells mate, which suggests that the *p*^*rep1*^ promoter is “leaky” even before partners grow towards each other. Note that as partners mate, the green fluorescence is upregulated first in P-gametes and only subsequently in M-gametes.

**(K)** The graph reports mean cellular intensity for 3sfGFP-NLS expressed from the *p*^*rep1*^ promoter shown in (J) throughout mating.

**(L)** The micrographs show the PP7 reporter (green) detailed in (E) that we used to monitor *rep1* gene transcription. The mCherry (magenta) as in (F). Boxplots report the onset of PP7 reporter dot detection in P- and M-gametes. In mating populations, we observed transient bursts of *rep1* PP7 reporter activity in both P- and M-gametes even before mating took place, which is consistent with the observed “leakiness” of the *rep1* promoter (J-K). Importantly, as partner gametes engaged each other, we detected *rep1* transcription bursts more frequently in P-than M-gametes (M). In the final mating steps, *rep1* PP7 reporter activity bursts occurred with similar frequency in both partners (M). Consistent with the observed *rep1* promoter activity (J-K), our results show that the *rep1* transcription occurs more robustly in P-than M-gametes.

**(M)** The graph reports mean detection frequency (lines) for the *rep1* transcription reporter shown in (L) in the 3 h before gamete fusion. Note significantly more frequent detection in P-gametes during 60-120 min before fertilization. Shaded areas report the Wilson interval and the p-values are from the exact McNemar’s test.

**(N)** The graphs report the mean cellular intensity (top) and nucleocytoplasmic ratio (bottom) for Ste11-sfGFP in mating wild-type P- and M-gametes, as indicated. We used the paired t-test to compare the nucleocytoplasmic ratio in P- and M-gametes and report values lower than 0.05 with an asterisk (*). Note that the Ste11-sfGFP nuclear targeting is significantly higher in P-than M-gametes early in the mating process.

All timepoints are relative to partner cell fusion and gamete mating types are indicated (P/M). Scale bars are indicated in micrographs, white dashed lines indicate cell boundaries visualized from brightfield images. Dots represent individual datapoints and we indicate the number of analysed mating events (n). Unless indicated otherwise, we show Kruskal-Wallis test p-values. We show either exact p-values or indicate p-value range with asterisks as 0.05 > *. In boxplots, box edges indicate 25th and 75th percentiles, the central line denotes the median, and whiskers extend 1.5 times the interquartile range. In line graphs, lines connect mean values and shaded areas report standard deviation.

**Figure S3. Asymmetries in the early expression of several meiotic genes in P-gametes are regulated by pheromone signalling**

**(A)** As indicated, bar charts report mating and sporulation efficiencies, and aberrant sporulation rates in mating mixtures of either wild-type or *mfm1Δ mfm3Δ* mutant strains 48 h upon mating induction. Bars report mean values, error bars denote standard deviation, and dots indicated individual datapoints. Note that the *mfm1Δ mfm3Δ* mutants, which delay the onset of mating, show only minor defects in sexual reproduction at 48 h after mating induction.

**(B)** Micrographs show Ste11-sfGFP (green) during mating of *mfm1Δ mfm3Δ* mutant partners. The mTagBFP2 (blue) as in (C). Nuclear boundaries, which we visualized with the nuclear ^NLS^mCherry marker (**Mov.2**), are indicated with orange dashed lines. Boxplots report the Ste11-sfGFP nucleocytoplasmic (N/C) ratio in P- and M-gametes 80 min before fusion. Note that M-gametes shows higher nuclear Ste11 targeting than the P-partners 80 min before partners fuse (arrowhead, boxplots).

**(C)** The graphs report the mean cellular intensity (top) and nucleocytoplasmic ratio (bottom) for Ste11-sfGFP in mating *mfm1Δ mfm3Δ* P- and M-gametes, as indicated. We used the paired t-test to compare the nucleocytoplasmic ratio in P- and M-gametes and report values lower than 0.05 with an asterisk (*). Note that the Ste11-sfGFP nuclear targeting is significantly higher in M-than P-gametes early in the mating process.

**(D)** Micrographs show Rep1-sfGFP (green) during mating of homothallic strains lacking the *mfm1* and *mfm3* genes. The mTagBFP2 (blue) as in (C). Boxplots report the onset of Rep1-sfGFP signal detection in P- and M-gametes. Note that Rep1-sfGFP signal is produced first in M-gametes (arrowheads, boxplots) during mating of *mfm1Δ mfm3Δ* partners.

**(E)** In top panels, dot plots show the rate (*v*) and time of production onset (t_0_) for Rec8-sfGFP production in P- and M-gametes for individual *mfm1Δ mfm3Δ* mating pairs. The Rec8-sfGFP signal traces were fitted as in (B). The mean 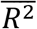, value across all fits for the P-gametes was 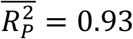, and for the M-gamete was 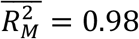. In the bottom panel, Rec8-signal in each partner from 60 to 10 min pre-fusion has been plotted and the best orthogonal regression fit, with an *R*^2^ = 0.11, is indicated with the red line. Note that partner gametes produce Rec8 at similar rates but first in M-cells. Note that the *mfm1Δ mfm3Δ* M-gametes produce approximately 3-fold more Rec8 than the M-gametes.

All timepoints are relative to partner cell fusion and gamete mating types are indicated (P/M). Scale bars are indicated in micrographs, white dashed lines indicate cell boundaries visualized from brightfield images. Dots represent individual datapoints and we indicate the number of analysed mating events (n). Unless indicated otherwise, we show Kruskal-Wallis test p-values. We show either exact p-values or indicate p-value range with asterisks as 0.05 > *. In boxplots, box edges indicate 25th and 75th percentiles, the central line denotes the median, and whiskers extend 1.5 times the interquartile range. In line graphs, lines connect mean values and shaded areas report standard deviation.

**Figure S4. A single *rec8* gene copy supports production of viable spores during azygotic meiosis. Sufficient Rec8 expression by either gamete ensures accurate meiosis in zygotes**.

**(A)** The schematic (left) illustrates zygotic and azygotic asci formation. Haploid gametes

fuse to produce a zygote, which enter meiosis to produce zygotic asci (red). Rare zygotes that do not immediately enter meiosis can be selected and propagated mitotically as diploid cells in nitrogen-rich conditions (blue). Upon nitrogen depletion, diploids undergo meiosis and sporulation to produce azygotic asci (see Methods). The bar chart (right) reports the viability of spores produced in azygotic asci^5^ of diploid strains with indicated genotypes. Even though differences between parental alleles have not been reported for diploid cells, or during azygotic meiosis, we indicate the origin of *rec8* alleles in diploid cells. Note that deleting both *rec8* copies severely reduces viability of spores produced by diploid cells but that removal of one *rec8* allele does not significantly reduce spore viability.

**(B)** Timelapses show localization of the Hht1 histone H3^6^ fused to the red fluorescent protein RFP (magenta) during mating of either wild-type or *rec8Δ* mutants, as indicated. The brightfield images are overlayed in the first and last timepoints. We did not observe defects in fusion of gametic nuclei in wild-type or zygotes produced by *rec8* mutant gametes (0/100 zygotes for each cross, **Mov.3**). Meiotic divisions produced four equal chromatin masses of Hht1-RFP in majority of zygotes produced by two wild-type parents (arrowheads). Zygotes with two *rec8Δ* parents often segregated Hht1-RFP into aberrant number of uneven chromatin masses (arrows).

**(C)** The bar chart reports the frequency of Hht1-RFP meiotic segregation defects described in (B) in zygotes produced by indicated crosses of heterothallic wild-type and *rec8Δ* mutant strains. Note that deleting *rec8* only in P-gametes causes more pronounced chromatin segregation defects than deleting *rec8* only in M-gametes.

**(D)** The schematic shows the design of synthetic reporter alleles used to monitor meiotic allele segregation. The *leu1+* genomic locus on chromosome II was used to integrate the sporulation-induced promoter *p*^*pdc202*^ followed by three tandem repeats of fluorescent proteins mCherry, in one parent, or mNeonGreen^7^, in the other parent. Crosses between the two parents produces spores with fluorescence that indicates the inherited alleles. Micrographs show the different types of asci produced by crossing otherwise wild-type strains that carry the two reporter alleles. We show the overlay of brightfield with green and red fluorescent channels (gray, green and magenta, respectively). Green and magenta arrowheads point to spores emitting corresponding fluorescent signals, and white arrowheads indicate spores that do not produce fluorescence. We show asci that accurately segregate alleles between four spores (top; 91±1%), form four spores but missegregate reporter alleles (middle, note the spore with red and green fluorescent signal; 6±1%), or form fewer than four spores (bottom; 3±1%).

**(E)** Stacked histograms shown the frequency of asci shown in (D) and observed in indicated crosses of heterothallic *mfm1Δ mfm3Δ* mutant strains where the *rec8* gene is either intact or deleted. The strains carry the reporter alleles detailed in (D). Note that in *mfm1Δ mfm3Δ* mutants, deleting *rec8* only in M-gametes causes more pronounced meiotic segregation defects than deleting *rec8* in P-gametes only. Note that deleting *rec8* in both *mfm1Δ mfm3Δ* partners strongly impairs allele segregation during meiosis. Welch test p-values are shown.

**(F)** Micrographs show Rec8-sfGFP (green) expressed from the M-cell specific *p*^*mam1*^ promoter during mating of partners that lack the native *rec8* gene. P-gametes utilize the P-cell-specific *p*^*map3*^ promoter to express mTagBFP2 (blue), which we used to determine the mating type of gametes and time of partner fusion. Dashed lines indicate cell boundaries visualized from brightfield images. We did not quantify the time of Rec8-sfGFP induction relative to cell fusion since M-cells activate the *p*^*mam1*^ promoter already at the start of the experiment (arrowheads), which is not correlated to the time of cell fusion.

**(G)** Stacked histograms show frequency of asci shown in (D) in mating mixture produced by wild-type gametes and crosses where Rec8 is either lacking or solely expressed from the M-gamete-specific *p*^*mam1*^ promoter, the mating-induced *p*^*pak2*^ promoter or zygotic *p*^*mei3*^ promoter, as indicated. Additionally, strains carry the reporter alleles detailed in (D). Note the largely accurate meiotic allele segregation in crosses where *p*^*mam1*^ and *p*^*pak2*^ promoters drive Rec8 expression in gametes, and high incidence of meiotic defects when Rec8 is expressed at cell fusion from the *p*^*mei3*^ promoter. Welch test p-values are shown.

**(H-I)** Micrographs show Rec8-sfGFP (green) expressed from the mating-induced *p*^*pak2*^ promoter (H) and the zygotic *p*^*mei3*^ promoter (I) during mating of homothallic strains where native *rec8* alleles are deleted. The M-gamete-specific *p*^*mam1*^ promoter regulates expression of mTagBFP2 (blue), which we used to determine the gametes mating type and the time of fusion. Boxplots report the onset of Rec8-sfGFP detection in P- and M-gametes. Kruskal-Wallis test p-values are shown. Note that Rec8-sfGFP expression from the *p*^*pak2*^ promoter starts earlier, and the expression from the *p*^*mei3*^ promoter later, than the wild-type Rec8 expression (**Fig.1C**).

**(J-K)** Bar charts report the time from partner fusion until the S-phase is triggered in indicated crosses between heterothallic strains where the native *rec8* gene is either deleted or fused to the sfGFP sequence. Additionally, we indicate the strain that carries the *mei3Δ* mutation (K). Kruskal-Wallis test p-value for indicated comparison is shown. Note that mutating *rec8* gene in either parent does not affect the time of pre-meiotic S-phase onset in zygotes (J). Also note that that deleting the *mei3* P-allele significantly delays premeiotic S-phase initiation in zygotes (K).

All timepoints are relative to partner cell fusion and gamete mating types are indicated (P/M). Scale bars are indicated in micrographs, white dashed lines indicate cell boundaries visualized from brightfield images. Dots represent individual datapoints and we indicate the number of analysed replicates (N) and relevant cell types (n). In bar charts and stacked histograms, bars report means and error bars denote standard deviation. In boxplots, box edges indicate 25th and 75th percentiles, the central line denotes the median, and whiskers extend 1.5 times the interquartile range. We report exact p-values from statistics tests reported detailed in panel legends.

**Figure S5. The *rec8* P-allele contributes more to meiotic success than the M-allele primarily by promoting meiotic centromere function and reductional meiosis I**

**(A)** Micrographs show mating between indicated wild-type, *rec8Δ* and *rec11Δ* partners where M-gametes carry the *lys3* locus labeled with the *parS*^*c3*^/ParB^c3^-eGFP (green) and P-gametes carry the *lys3* locus labeled with the *parS*^*c2*^/ParB^c2^-mCherry-NLS (see **Mov.7**). Cell cycle phases are indicated (G_1_, G_2_). Note that the *lys3* locus-recruited *parS*^*c3*^/ParB^c3^-eGFP (arrowheads, insets) produces a single dot prior to meiosis in wild-type zygotes but two dots in zygotes that lack either parental *rec8* allele.

**(B)** Micrographs show mating between indicated wild-type, *rec11Δ* and *rec8Δ* partners where both gametes carry the *rec6* locus labeled with the *parS*^*c3*^/ParB^c3^-eGFP (green). Cell cycle phases are indicated (G_1_, G_2_, MI and post-MII). Note that during meiosis I, in wild-type zygotes the pairs of *rec6* loci segregate to opposite poles (homologue disjunction) and that in *rec8* mutants the four *rec6* loci co-segregate together (homologue non-disjunction, see insets).

**(C)** For indicated crosses between wild-type and *rec8Δ* mutants, stacked histograms report the frequency of phenotypes shown in (B), where during meiosis I parental *rec6* loci segregate as pairs to opposite poles (homologue disjunction) or co-segregate all together (homologue non-disjunction).

**(D)** Boxplots report the viability of spores produced in indicated crosses of wild-type and *rec11Δ* mutant gametes. Tukey test p-values are shown. Note that deleting *rec11* in the P-gamete reduces spore viability more than deleting it in the M-gamete.

**(E)** Stacked histograms report the frequency of tetrad asci and asci containing aberrant spore number for indicated crosses between wild-type, *rec11Δ* and *rec8Δ* mutant gametes. Welch test p-values are shown. Note that meiotic defects are more pronounced upon deleting *rec8* than *rec11*, and that *rec8Δ* is epistatic to *rec11Δ*. All timepoints are relative to cell fusion. Scale bars are indicated in micrographs, white dashed lines indicate cell boundaries visualized from brightfield images. Dots represent individual datapoints and we indicate the number of analyzed replicates (N) and relevant cell types (n). In stacked histograms, bars report means and error bars denote standard deviation. In boxplots, box edges indicate 25th and 75th percentiles, the central line denotes the median, and whiskers extend 1.5 times the interquartile range.

**Figure S6. Mating produces a population of Rec8-expressing P-gametes and increases genomic instabilities in P-gametes that return to mitotic proliferation**.

**(A)** Homothallic cells coding for Rec8-sfGFP and expressing mTagBFP2 from the M-cell-specific *p*^*mam1*^ promoter, were induced to mate and the mating was interrupted by shifting cells to growth media. Boxplots report mean mTagBFP2 cellular fluorescence for gametes that either induce or lack detectable Rec8 signal at the time mating is interrupted. The Kruskal-Wallis test p-value is shown. Note that the gametes with detectable Rec8-sfGFP signal have lower mTagBFP2 levels and are thus likely enriched in P-gametes. Note that mTagBFP2 signal can be reliably used to determine the mating type of gametes only within a mating pair (*e*.*g*. in **Fig.1F** partners show small difference in mTagBFP2 levels).

**(B)** Micrographs show Rec8-sfGFP (green) and ^NLS^mCherry (magenta) signals upon transferring the indicated heterothallic mating mixtures to growth media. We used cell morphology to distinguish gametes from zygotes, and the ^NLS^mCherry signal to determine the mating type of gametes (P, M). Note the Rec8-sfGFP in P-gametes (arrowheads). Dashed lines indicate cell boundaries visualized from brightfield images.

**(C)** Stacked histogram shows the frequency of P- and M-mating type for gametes that induced Rec8 upon retrieval from indicated heterothallic mating mixtures onto growth media, as shown in (B). The Kruskal-Wallis test p-value reports the significance for the over-representation of the P mating-type in unfused Rec8-expressing gametes. Note that the Rec8-expressing gametes are predominantly the P-mating type.

**(D)** The schematic illustrates the viability assay for mated and unmated gametes from mating mixtures. Since fission yeast mating is asynchronous, we used a microscopic needle to isolate individual gametes or mating pairs produced between heterothallic *fus1Δ* mutant P-gametes (green, blue) and M-gametes (orange) and transferred them onto growth media to assess their ability to form colonies. To ensure that *rec8+* and r*ec8Δ* P-gametes (green and blue, respectively) experience identical mating conditions, we pooled the two strains together before introducing the M-cells. Distinct selection markers were used to determine the colony progenitors, and we report gamete viability as the ratio between observed and expected number of colonies forming. We used the same approach to determine gamete viability in mating mixtures of *rec8+* and r*ec8Δ* M-gametes pooled with a single P-partner strain.

**(E)** The bar chart reports viability of P- and M-cells from fission yeast mating mixtures with indicated genotypes. Tukey test p-values are shown. Note that mating reduces viability of P-cells, and that this is largely dependent on the *rec8* gene and partially dependent on the *rec11* gene.

**(F)** The schematic (left) shows the design of the loss-of-heterozygosity (LOH) assay^8^, which detects both copy number loss and copy number-neutral allele loss. Briefly, because only *h+/h-* diploids acquire zygotic identity, the *h+/h+* and *h-/h-* diploids are stable and able to mate. We obtained the diploids by fusing protoplasts of haploid cells that carry *ade6-M210* (red) and *ade6-M216* (pink) mutations, which are individually auxotrophic but capable of interallelic complementation that restores adenine prototrophy in diploid cells. Loss of either *ade6* allele results in auxotrophic cells. Note that adenine prototrophs produce white colonies whereas adenine auxotrophs results in formation of colonies with red/pink hue that are easily visualized (arrowheads) as shown in the photograph (right). We deleted both alleles of the *fus1* gene to ensure that upon mating *h+/h+* and *h-/h-* diploids are unable to fuse.

**(G)** The bar chat shows LOH rates for *ade6* locus in mated and non-mated strains with indicated genotypes, which we measured using the assay detailed in (F). We report the results for *fus1Δ/fus1Δ* mutant *h+/h+* and *h-/h-* diploids that were starved for nitrogen either in the presence or the absence of the mating partner before assessing the LOH rates. The *dcr1Δ/dcr1Δ* mutant was used as a positive control for the LOH assay^8^. Tukey test p-values are shown. Note that LOH rates increase in mating populations, and that the effect is significantly greater in P-than M-gametes. Also note that deletion of *rec8* reduces LOH rates in mated P-gametes.

**(H)** The schematic shows the experimental design used to evolve fission yeast populations of P- and M-gametes in **Fig.3E**. We subjected populations of 10^7^ cells initially containing the same number of heterothallic P- and M-gametes to cycles of mating and regrowth. Note that partners could not fuse because we deleted the *fus1* gene. We performed the experiment using P- and M-gametes that either both carried the wild-type *rec8* gene or both lacked the *rec8* gene.

Scale bars are shown for micrographs, white dashed lines indicate cell boundaries visualized from brightfield images and gamete mating types are indicated (P/M). Dots represent individual datapoints and we indicate the number of analyzed replicates (N) and relevant cell types (n). In boxplots, the top and bottom box edges indicate 25th and 75th percentiles, and the central line displays the median. The whiskers extend 1.5 times the interquartile range. Dots represent individual datapoints. In bar charts and stacked histograms, bars report means and error bars denote standard deviation.

**Figure S7. Quantifications and fits of zygotic success and spore viability as a function of Rec8 levels**

(**A**) The plot reports the observed Rec8-sfGFP signal ratio between wild-type P- and M-gametes at indicated timepoints (diamonds), with intervals (bars) indicating the average of signal ratios calculated for timepoints in the interval pre-fusion. Each time point ratio was calculated using orthogonal regression using intensity data associated with indicated timepoints. For modeling, we relied on the P/M ratio of 3.6 in Rec8 levels, which is the average of the observed ratios calculated at -10, -20 and -30 minutes (red bar). We do not to include earlier timepoints because very low Rec8 expression in M-cells causes high signal-to-noise ratio and hence inflates the asymmetry.

**(B)** Boxplots report mean Rec8-sfGFP fluorescence at indicated timepoints in crosses of gametes that carry the native *rec8* gene tagged with sfGFP (1 copy) or gametes that carry an additional copy of *rec8* gene tagged with sfGFP (2 copies). Note that the presence of two *rec8-sfGFP* gene copies approximately doubles the Rec8-sfGFP levels in gametes and zygotes.

(**C**) The schematic illustrates how the levels of Rec8 in gametes and zygotes were manipulated to determine the 5(·) and 7(·) dependence on Rec8 levels. We normalized the Rec8 levels detected in wild-type M-gametes ahead of partner fusion to 1 A.U. and used the relative levels of Rec8 in wild-type P-gametes of 3.6 A.U., as detailed in (A). Introducing each additional *rec8* gene copy, as indicated on the axes, was assumed to each provide one fold increase of the Rec8 protein, based on results obtained in (B). Thus, each additional *rec8* gene copy in M- and P-gametes was assumed to increase the Rec8 levels by 1 and 3.6 A.U., respectively. We considered Rec8 expressed in each gamete as functionally equal when evaluating the effect of Rec8 level on gamete survival. Likewise, we considered Rec8 supplied by each parent functionally equal for spore viability, and, as indicated, we mated gametes with different *rec8* gene copy number to evaluate how total parental Rec8 levels affect zygote fitness.

**(D)** The experimental data used to fit the probability 5(8) of successful meiosis as a function of total meiotic factor investment 8. The boxplot reports viability of spores produced by crosses of heterothallic partners that supply indicated levels of Rec8, which are noted in A.U. as detailed in (C). Heterothallic gamete populations were allowed to mate and sporulate, before collecting a defined number of spores and plating them onto growth media to assess their ability to form colonies.

**(E)** The experimental data used to fit the probability *q*(*r*_*p*/*m*_) of successful return to mitosis as a function of meiotic factor *r*_*p*/*m*_ produced in unfused gametes. The boxplot reports viability of mated gametes expressing indicated levels of Rec8, which are noted in A.U. as detailed in (C). Gamete populations were allowed to mate and to form mating pairs but could not fuse due to deletion of the *fus1* gene. Individual mating pairs were then transferred onto growth media and the ability to form colonies was assessed. In boxplots, top and bottom box edges indicate 25th and 75th percentiles, and the central line displays the median. The whiskers extend 1.5 times the interquartile range. Dots represent individual datapoints and we indicate the number of analyzed replicates (N) and relevant cell types (n).

## MOVIE LEGENDS

**Movie 1. P-gametes induce expression of meiotic genes before partner M-gametes**.

**(A)** The timelapse shows endogenous Rec8 tagged with sfGFP (green) during mating of homothallic wild-type cells. The M-gamete-specific *p*^*mam1*^ promoter drives expression of cytosolic mTagBFP2 (blue), which we used to determine the gametes mating type (P, M) and time of fusion, which is observed as cytosolic mixing between partners. Brightfield micrographs are also shown (gray). The yellow dashed lines indicate a mating pair of interest. Note that cells show the faint background fluorescence in the green channel in the form of dots and strings, most likely corresponding to mitochondria. Note that Rec8-sfGFP produces a diffuse nuclear signal in addition to prominent nuclear foci that are consistent with reported Rec8 enrichment at centromeres^9^. Note the Rec8-sfGFP signal appearance prior to gamete fusion and first in the centrally positioned nucleus of the P-gametes, followed by M-gametes. Note the continued Rec8-sfGFP production in zygotes and that the protein is degraded during meiotic divisions and ahead of spore formation.

**(B)** The timelapse shows endogenous Rep1 tagged with sfGFP (green) during mating of homothallic wild-type cells. The mTagBFP2 (blue) and the brightfield (gray) images, as well as yellow dashed lines, as in (A). Note that Rep1-sfGFP produces a diffuse nuclear signal that appears during mating and first in P-partners. Note that Rep1-sfGFP signal peaks shortly after partners fuse, followed by a signal decrease and complete degradation at meiotic divisions.

**(C)** The timelapse shows endogenous Ste11 tagged with sfGFP (green) in homothallic wild-type cells immediately after the shift to nitrogen-free media and throughout sexual reproduction. Cells co-express mCherry (magenta) fused with the nuclear localization signal (NLS) that labels nuclei. The mTagBFP2 (blue) and the brightfield (gray) images, as well as yellow dashed lines, as in (A). Note that Ste11-sfGFP is already induced in cells that did not arrest the mitotic divisions, with comparable signal in the nucleus and the cytosol. Note that during mating, Ste11 levels and nuclear targeting increase, and then rapidly decrease upon gamete fusion as Ste11 is degraded in zygotes. The differences in Ste11 nucleocytoplasmic ratio in P- and M-gametes are reported in **Fig.1F, S2N**.

All timepoints are relative to the start of the timelapse. Scale bars are indicated.

**Movie 2. Pheromone communication regulates the bias in meiotic gene expression between P- and M-gametes**.

**(A)**The timelapse shows endogenous Ste11 tagged with sfGFP (green) in homothallic *mfm1Δ mfm3Δ* mutants immediately after the shift to nitrogen-free media and throughout sexual reproduction. The M-gamete-specific *p*^*mam1*^ promoter drives expression of cytosolic mTagBFP2 (blue), which we used to determine the gametes mating type (P, M) and the time of fusion, which is observed as cytosolic mixing between partners. Cells co-express mCherry (magenta) fused with the nuclear localization signal (NLS) that labels nuclei. Brightfield micrographs are also shown (gray). The yellow dashed lines indicate a mating pair of interest. Note that Ste11-sfGFP is already induced in cells that did not arrest the mitotic divisions, with comparable signal in the nucleus and the cytosol. The differences in Ste11 nucleocytoplasmic ratio in P- and M-gametes are reported in **Fig.S3B-S3C**.

**(B)** The timelapse shows endogenous Rep1 tagged with sfGFP (green) during mating of homothallic *mfm1Δ mfm3Δ* mutants. The mTagBFP2 (blue) and the brightfield (gray) images, as well as yellow dashed lines, as in (A). Note that Rep1-sfGFP produces a diffuse nuclear signal that appears during mating and first in M-partners. **(C)** The timelapse shows endogenous Rec8 tagged with sfGFP (green) during mating of homothallic *mfm1Δ mfm3Δ* mutants. The mTagBFP2 (blue) and the brightfield (gray) images, as well as yellow dashed lines, as in (A). Note the Rec8-sfGFP signal appearance prior to partner fusion and first in M-gametes.

All timepoints are relative to the start of the timelapse. Scale bars are indicated.

**Movie 3. Rec8 is required for accurate meiotic segregation of chromatin**.

Timelapses show localization of the Hht1 histone H3^6^ fused to the red fluorescent protein RFP (magenta) during mating of either wild-type or *rec8Δ* mutants, as indicated. Brightfield channel is shown (gray). The yellow dashed lines indicate a mating pair of interest. Note that zygotes with two wild-type parents (top row), or mutant *rec8Δ* M-parent (second row), produce four equal chromatin masses of Hht1-RFP (arrowheads), whereas meiotic divisions in zygotes with *rec8Δ* P-parent (third row), or two *rec8Δ* parents (bottom row), segregate Hht1-RFP into aberrant number of uneven chromatin masses (arrowheads). Also note that after cells fuse, partner nuclei undergo karyogamy in both wild-type and mutant zygotes. All timepoints are relative to the start of the timelapse. Scale bars are indicated.

**Movie 4. Rec8 expression in gametes promotes accurate sporulation**.

Timelapses show Rec8-sfGFP (green) expressed from the M-gamete-specific *p*^*mam1*^ (**A**), mating-induced *p*^*pak2*^ (**B**), and zygotic *p*^*mei3*^ (**C**) promoter during mating between parents that lack the native *rec8* gene. The cells co-express the mTagBFP from either the P-cell-specific *p*^*map3*^ (**A, B**) or the M-gamete-specific *p*^*mam1*^ (**C**) promoter, which we used to determine gametes mating type (P,M) and the time of cell fusion. Brightfield micrographs are also shown (gray). The yellow dashed lines indicate a mating pair of interest. Note that Rec8-sfGFP is induced from the *p*^*mam1*^ promoter hours before cell fusion and in M-gametes only, from the *p*^*pak2*^ promoter in both gametes during mating, and from the *p*^*mei3*^ promoter at the time of fusion. Note that four spores (white arrowheads) form in zygotes driving expression of Rec8-sfGFP from *p*^*mam1*^ or *p*^*pak2*^ promoters, whereas the zygote driving expression of Rec8-sfGFP from *p*^*mei3*^ promoter produces aberrant number of spores. All timepoints are relative to the start of the timelapse. Scale bars are indicated.

**Movie 5. P- and M-alleles produce distinct amounts of Rec8 at the onset of pre-meiotic S-phase**.

Timelapses show Rec8-sfGFP (green) expressed from either one or both parental alleles, as indicated. Cells co-express mCherry-Pcn1 (magenta), which during DNA-replication forms foci (magenta arrowheads) and a uniform nuclear signal outside S-phase. The M-gamete-specific *p*^*mam1*^ promoter drives expression of mTagBFP2 (blue), which we used to determine gametes mating type (P,M) and time of cell fusion. Brightfield micrographs are also shown (gray). The yellow dashed lines indicate a mating pair of interest. Note that the Rec8-sfGFP signal at the onset of S-phase (green arrowheads) is lower when produced by the M-allele alone (bottom) than that produced by the P-allele alone (middle) or both parental alleles (top). All timepoints are relative to the start of the timelapse. Scale bars are indicated.

**Movie 6. Delaying the pre-meiotic S-phase allows for increased Rec8 production from the M-allele ahead of DNA replication**

Timelapses show Rec8-sfGFP (green) expressed from the M-allele in indicated crosses where P-gametes lack the *rec8* gene, and the *mei3* gene is either intact or deleted. Cells co-express mCherry-Pcn1 (magenta), which during DNA-replication forms foci (magenta arrowheads) and a uniform nuclear signal outside S-phase. The M-gamete-specific *p*^*mam1*^ promoter drives expression of mTagBFP2 (blue), which we used to determine gametes mating type (P,M) and time of cell fusion. In the brightfield channel (gray), the yellow lines indicate a mating pair of interest. Note that deleting the *mei3* in P-gametes delays pre-meiotic S-phase, which allows for increased Rec8-sfGFP production (green arrowheads) from the M-allele by the time DNA replication starts. Note the defective spore number (white arrowheads) in crosses where P-gametes lack the *rec8* gene only, and formation of four spores in zygotes that lack *rec8* and *mei3* genes. All timepoints are relative to the start of the timelapse. Scale bars are indicated.

**Movie 7. Native Rec8 levels are required for tight cohesion at chromosome arms**

The timelapses show mating between indicated wild-type, *rec8Δ* and *rec11Δ* partners where P-gametes carry the *lys3* locus labeled with the *parS*^*c3*^/ParB^c3^-eGFP (green) and M-gametes carry the *lys3* locus labeled with the *parS*^*c2*^/ParB^c2^-mCherry-NLS (magenta). Brightfield micrographs are also shown (gray). Note that the *lys3* locus-recruited *parS*^*c3*^/ParB^c3^-eGFP (arrowheads) produces a single dot prior to meiosis in wild-type zygotes but two dots in zygotes that lack either parental *rec8* allele. All timepoints are relative to the start of the timelapse. Scale bars are indicated.

**Movie 8. Zygotes lacking parental *rec8* alleles undergo aberrant, equational meiosis I**

The timelapses show mating between indicated wild-type, *rec8Δ* and *rec11Δ* partners where P-gametes carry the *rec6* locus labeled with the *parS*^*c3*^/ParB^c3^-eGFP (green) and M-gametes carry the *rec6* locus labeled with the *parS*^*c2*^/ParB^c2^-mCherry-NLS (magenta). Brightfield micrographs are also shown (gray). Note that the *rec6* locus-recruited *parS*^*c3*^/ParB^c3^-eGFP (arrowheads) produces a single dot prior to meiosis in wild-type zygotes but two dots in zygotes that lack either parental *rec8* allele. Also note that at meiosis I, in wild-type zygotes the *rec6* loci co-segregate and in *rec8* mutants the foci separate. All timepoints are relative to the start of the timelapse. Scale bars are indicated.

**Movie 9. Zygotes lacking Rec11 fail to segregate homologues**

The timelapses show mating between wild-type or *rec11Δ* partners where both gametes carry the *rec6* locus labeled with the *parS*^*c3*^/ParB^c3^-eGFP (green). Brightfield micrographs are also shown (gray). Note that during meiosis I, in wild-type zygotes the pairs of *rec6* loci segregate to opposite poles (homologue disjunction) and that in *rec8* mutants the four *rec6* loci co-segregate together (homologue non-disjunction).

**Movie 10. Mating produces a subpopulation of P-gametes that induce Rec8 expression but fail to fuse with a partner**

The timelapse shows endogenous Rec8 tagged with sfGFP (green) during mating of homothallic wild-type cells. The M-gamete-specific *p*^*mam1*^ promoter drives expression of cytosolic mTagBFP2 (blue), which we used to determine the gametes mating type (P, M) and time of fusion, which is observed as cytosolic mixing between partners. Brightfield micrographs are also shown (gray). The yellow dashed lines indicate the three cells that engage in mating. Note that two P-gametes induce Rec8-sfGFP expression during competition for the same M-partner. All timepoints are relative to the start of the timelapse. Scale bars are indicated.

**Movie 11. Rec8 expression during mating causes mitotic and growth defects in gametes that return to proliferation**.

The timelapse shows homothallic cells carrying endogenous *rec8* tagged with sfGFP. Cells were induced to mate but the mating was interrupted by returning cells to growth conditions at the start of the imaging. We show the Rec8-sfGFP (green) upon return to growth media (first timepoint) and brightfield images (gray) for all timepoints. Note that Rec8-sfGFP is expressed in several unfused gametes. Note that the majority of unfused gametes resume growth and mitotic divisions. In the left panel, note that the outlined gamete expresses Rec8-sfGFP and that it lyses at first division. In the right panel, note that the outlined gamete expresses Rec8-sfGFP and that one of its daughter cells arrests growth. All timepoints are relative to the start of the timelapse. Scale bar is indicated.

**Movie 12. Evolution of asymmetric parental investment strategy using theoretical model of the fission yeast lifecycle**

Each magenta dot represents an individual genotype (*r*_*p*_, *r*_*m*_) evolving in the 2-dimensional genotype space according to our theoretical model. We use the colorscale to associates points in the genotype space with the expected number of offspring given by *E*(*r*_*p*_, *r*_*m*_) = 2(1 − *f*)[*q*(*r*_*p*_) + *q*(*r*_*m*_)] + 4*fP*_*m*_(*r*_*p*_ + *r*_*m*_), and hence visualize the “fitness landscape”. The population is initialized with 10,000 individuals with an identical, symmetric genotype (1,1). Each frame represents a step of 100 evolutionary generations.

## METHODS

### Growth conditions

Yeast cultures were maintained and propagated using standard fission yeast media and protocols^10^. The composition of all media is detailed in the **Table S1**. Strain husbandry was carried out in YES, EMM and MSL+N media, with incubations carried out at 18°C, 25°C or 30°C, either on solid plates or in liquid cultures shaking at 200 rpm. Cell density was measured as a proxy of cell number using Ultrospec 10 cell density meter (Biochrom, Holliston, USA), where an O.D._600_ = 0.5 corresponded to ∼1·10^7^ cells/mL.

**Genetic crosses** were performed by combining freshly streaked strains on MSL−N plates. Tetrad dissections were performed using the MSM 400 micromanipulator (Singer Instruments, Somerset, United Kingdom) on mating mixtures incubated for ∼24 hours at 25°C. Following dissection onto YES solid plates, colonies were grown and replica-plated onto antibiotic-containing YES plates to determine genetic markers. Plates were imaged with the BioDoc-It™ 220 system (UVP, Analytik Jena, Jena, Germany).

**Selection** for prototrophs was carried out using EMM media that lacked specific nutrients. To select using antibiotics, we supplemented YES medium with 100 µg/mL G418 (AG11347, Biosynth, Staad, Switzerland), 100 µg/mL nourseothricin (AB-102L, Jena Bioscience, Jena, Germany), 50 µg/mL hygromycin B (HYG5000, Formedium, United Kingdom), 100 µg/mL zeocin (ant-zn-5p, Invivogen, San Diego, USA) and 15 µg/mL blasticidin-S (ant-bl-1, San Diego, USA). Glufosinate selection was performed on EMM supplemented with 400 µg/mL glufosinate (FP13608, Biosynth, Staad, Switzerland).

To **culture cells for experimental procedures**, we used previously reported protocols^10,11^. Briefly, cells from freshly streaked YES plates were first pre-inoculated into liquid MSL+N medium and incubated either overnight at 25°C or for 8 hours at 30°C with continuous shaking at 200 rpm. Cultures were then diluted to O.D._600_ = 0.025 in MSL+N medium and incubated overnight at either 25°C or 30°C until they reached exponential phase. To **induce nitrogen starvation** response, cells were collected by centrifugation at 1,000 g for 1 min and washed three times in 1 mL of nitrogen-free MSL-N medium. To avoid mating, we nitrogen-starved heterothallic strains individually. To **induce mating**, we nitrogen-starved homothallic cell cultures, or we mixed equal cell number of separately cultured compatible heterothallic cells. For **timelapse microscopy of mating and sporulation**, we either immediately mounted MSL-N washed cells into imaging chambers — when monitoring Ste11-sfGFP dynamics and expression of Rec8 from the *p*^*mam1*^ promoter — or we resuspended cells to O.D._600_ = 1 in 3 mL of MSL-N liquid media and incubated cultures at 30°C with shaking at 200 rpm for 3-5 hours before mounting cells for imaging. To prepare **mating mixtures on solid media** we concentrated ∼2·10^7^ of MSL-N starved cells in 20 µL of the media, spotted them onto MSL-N agar plates and allowed samples to air-dry before incubation. To **measure mating and sporulation efficiencies** and to **obtain asci and spores** for downstream analyses, we incubate mating mixtures on agar plates for 24 h at 25°C. To **interrupt mating**, we incubate mating mixtures on agar plates for 6-8 h at 25°C before transferring cells into imaging chambers containing YES media, which were immediately subjected to imaging. To **isolate unmated gametes and mating pairs**, we washed *fus1Δ* mutant partners in MSL-N media, and immediately spotted cells on agar plates for 18 h at 25°C. The mating mixtures were than transferred to YES agar plates and individual gametes and mating pairs isolated using the MSM 400 micromanipulator (Singer Instruments, Somerset, United Kingdom).

### Strains and genetic markers

Fission yeast gene nomenclature, sequences, and functional annotations were obtained from the PomBase database^12^. A complete list of strains utilized in this study is provided in **Table S2**. The descriptions of alleles and genetic markers can be found in **Table S3**. Previously published markers were either sourced from the laboratory stocks, received as gifts from other research groups, or acquired from the National BioSource Project (yeast.nig.ac.jp/yeast/top.jsf).

We employed **homothallic** *h90* strains that undergo mating-type switching to produce both P- and M-gametes, or **heterothallic** *h+* and *h-* strains, which generate only P- or M-gametes, respectively. Unless otherwise noted, all strains used in this study are prototrophs.

To obtain the ***h+/h-* diploids**, which we used to produce azygotic asci, we performed crosses between haploid heterothallic strains that are unable to undergo mating type switching due to the H1Δ17 mutation that we introduced at the *mat1* mating-type locus^4^. Additionally, the *bleMX* and *natMX* antibiotic-resistance cassettes, which confer resistance to zeocin and nourseothricin, were integrated at the *mat1* locus in *h+* and *h-* strains, respectively. The two haploid strains were grown separately in MSL+N liquid media until exponential growth, equal number of cells pooled together and washed with MSL-N media and ∼2·10^7^ cells spotted for 6-8 h at 25°C on MSL-N solid media to induce mating and fusion. The mating mixture was then plated onto growth YES media supplemented with zeocin and nourseothricin to select for the rare *h+/h-* diploid colonies. After colonies developed, the sporulation proficient clones were selected for downstream experiments.

To obtain the ***h+/h+* and *h-/h-* diploids**, which we used in the loss-of-heterozygosity (LOH) assay detailed below, we performed protoplast fusion between haploid cells of the same mating type. We used either two haploid *h-* strains or two haploid strains with *mat1-P* locus (*h+*) unable to undergo mating type switching due to the H1Δ17 mutation^4^. Fusing haploids of the same mating type produced diploids that did not sporulate upon nitrogen starvation and instead mated with gametes of the opposite mating type. Additionally, the parental haploid strains carried either the *ade6-M210* or the *ade6-M216* mutation which cause adenine auxotrophy. Since the two *ade6* mutations show interallelic complementation in diploid cells, we selected for *ade6-M210/ade6-M216* diploids using EMM media lacking adenine. Specifically, the two haploid strains were cultivated individually in 25 mL of YES medium to exponential phase (O.D._600_=0.5), cells pooled together and harvested by centrifugation at 1,000 g for 3 min. The pellet was washed twice with 50 mL of sterile water, and once with 50 mL of 0.65 M solution of KCl (19677-0010, Acros Organics, Waltham, USA). Cells were transferred to 15 mL conical tubes and resuspended in 5 mL of Lallzyme solution consisting of 0.65 M KCl and 100 mg/mL Lallzyme MMX (69.316.10, Baldinger AG, Rümikon, Switzerland). The cell suspension was incubated 10-30 min at 30°C with nutation until cell wall digestion was complete and protoplasts clearly visible in the brightfield microscope. Protoplasts were collected by centrifugation at 150 g for 10 min, the supernatant was decanted, and cell integrity visually assessed by brightfield microscopy. The protoplast pellet was resuspended by gently pipetting with nipped 1 mL tips and washed three times with 10 mL of 1 M sorbitol using centrifugation at 150 g for 10 min between washes. Protoplasts were then resuspended in 1 mL of fresh PEG/CaCl_2_ solution prepared by mixing 0.9 mL of 30% w/v PEG4000 (0156.1, Roth, Karlsruhe, Germany) and 0.1 mL of 100 mM CaCl_2_ (0556-500G, VWR, Radnor, USA). Following a 30 min incubation at room temperature, protoplasts were pelleted by centrifugation at 150 g for 10 min, and 700 μL of the supernatant was carefully removed. The remaining cell suspension was gently mixed, and cell integrity was visually confirmed by microscopy. Finally, 150 μL of the cell suspension was plated onto selective EMM media plates lacking adenine and supplemented with 1 M sorbitol (35231.01, SERVA, Heidelberg, Germany). Plates were incubated at 30°C until diploid colonies ormed.

To introduce **genetic modifications** into haploid fission yeast, we performed standard lithium acetate transformation protocol^10^ . We transformed ∼700 ng of linear DNA fragments which carried homology arms at their 5’ and 3’ termini to target integration to the desired genomic locus. Typically, we obtained the transformation fragments by linearizing plasmid with restriction enzymes as detailed in **Table S3**. Transformants were selected based on prototrophic or antibiotic-resistance markers, and correct genomic integration was confirmed using genotyping PCRs or selection marker switching.

For alleles generated in this study and listed in **Table S3**, sequences of the **plasmids** used for their construction are available in the Supplemental Sequences on Figshare (dx.doi.org/10.6084/m9.figshare.29436455.v1). To integrate constructs at ectopic genomic loci, we obtained plasmids derived from the SIV plasmid series^13^, which integrate artificial constructs at the *ade6, his5, lys3*, or *ura4* loci and allow selection for prototrophs. Additionally, we developed plasmids to integrate constructs at the *leu1* genomic locus. To modify native genomic loci, we used the pFA6a backbone plasmid with antibiotic resistance cassettes *bleMX, bsdMX, hphMX, kanMX, natMX* or *patMX*^*14*^ that confer resistance to bleomycin/zeocin, blasticidin-S, hygromycin, G418, nourseothricin or glufosinate, respectively. We used standard molecular cloning techniques for plasmid construction using PCR-amplified genomic sequences. Sequences of DNA fragments amplified by PCR were validated using Sanger sequencing.

To obtain **gene deletions**, we replaced the entire protein coding regions with selection cassettes.

We used **promoters** of native genes to regulate gene expression. To achieve constitutive gene expression, we utilized the *p*^*pcn1*^, *p*^*act1*^, *p*^*tdh1*^ promoters^13^ . For mating type-specific gene expression, we employed either the P-cell-specific *p*^*map3*^ promoter or the M-cell-specific *p*^*mam1*^ promoter, which are induced already during the nitrogen-starvation response and further activated during mating^13^ . To induce gene expression in gametes of both mating types we used the *p*^*pak2*^ promoter^15^. To induce gene expression at the time of cell fusion we used the *p*^*mei3*^ promoter^4,16^. The *p*^*pdc202*^ promoter was used to induce gene expression during sporulation^17^ . The promoter of the *p*^*rep1*^ gene used to drive consisted of 922bp upstream of the START codon. **Transcriptional terminators** from fission yeast *rec8, nmt1*, and *tdh1* genes, as well as budding yeast ADH1 and CYC1 genes, were included at the 3’ termini of genetic constructs as indicated in plasmid sequences.

To visualize fission yeast proteins, we obtained N- or C-terminal fusions with fluorescent proteins sfGFP^18^ or mCherry^19^. Unless otherwise noted, tagging of the fission yeast proteins was performed at their endogenous loci.

To **visualize Rec11, Rec25, Rep1 and Ste11 expressed from their native loci** we introduced the sfGFP sequence at the C-terminus of the ORF followed by the transcriptional terminator of the budding yeast ADH1 gene. The **RFP tagged Hht1** was previously reported^6^.

To **visualize Rec8 expressed from the native locus**, we introduced the sfGFP sequence at the C-terminus of the native ORF. Since *rec8* mRNA is regulated by the DSR motifs^20,21^ at the C-terminus and the 3’UTR, the STOP codon of the sfGFP was followed by the DSR-containing sequences spanning from 92bp upstream to 346bp downstream of the *rec8* STOP codon.

To **advance Rec8 expression**, we placed the *rec8* ORF under the regulation of either the M-cell-specific *p*^*mam1*^ promoter or the gametic *p*^*pak2*^ promoter, and integrated constructs at the *ade6* genomic locus in cells lacking the native *rec8* gene. To **delay Rec8 expression**, we placed the *rec8* ORF under the zygotic *p*^*mei3*^ promoter and integrated the construct at the *ura4* genomic locus in cells lacking the native *rec8* gene. To monitor the dynamics of Rec8 expression from each promoter, the Rec8 was fused with the C-terminal sfGFP sequence. To **increase Rec8 expression** we introduced additional copies of the *rec8* gene containing 623bp promoter sequence upstream of the rec8 START codon, the *rec8* coding region containing introns and fused with sfGFP sequence at the C-terminus, and followed by the region carrying the DSR motifs^20,21^ from 92bp upstream to 156bp downstream of the *rec8* STOP codon. The construct was integrated at *ade6, ura4* and *leu1* loci to produce strains with additional 1-3 copies of the *rec8* gene.

We exploited the viral PP7 protein binding to RNA stem-loops to **visualize the nascent transcription of *rec8* and *rep1 in vivo***^22^. Specifically, we placed 24 PP7 stem-loops after the STOP codon of either gene, followed by the 3229bp of the budding yeast GLT1 ORF and the transcriptional terminator of the CYC1 gene that extend the transcript and thus increase its dwell time at the transcription site, which allows better signal detection. We fused the N-terminl sfGFP to the non-aggregating PP7ΔFG variant tag and expressed it from the *p*^*act1*^ promoter. In the same construct, we used the *p*^*act1*^ promoter to express the N-terminl sfGFP fused to the non-aggregating PP7ΔFG variant, followed by the transcriptional terminator from the budding yeast ADH1 gene. We noted that neither PP7 reporter caused gross defects in sporulation efficiency and production of four spores.

To monitor *p*^*rep1*^ promoter activity, we placed it upstream of the 3sfGFP fused with the SV40 nuclear localization signal (PKKKRKV) followed by the transcriptional terminator from the budding yeast ADH1 gene. This construct was integarated at the ura4 genomic locus.

To **visualize genomic loci** we employed two orthologous *parS*/ParB systems from *Burkholderia cenocepacia*^23^ . We placed the *parS*^*c2*^ or *parS*^*c3*^ sequences at the 3’ regions of genes *lys3* (∼3Mb from chromosome I centromere) and *rec6* (∼4kb from chromosome II centromere). At the same loci, we introduced cassetes that express fluorescently tagged ParB^c2^ or ParB^c3^ proteins that are loaded onto corresponding *parS* sequences. Specifically, we used the *p*^*tdh1*^ promoter to express the ParB^c3^ protein fused to eGFP and followed by the transcriptional terminator of the budding yeast CYC1 gene, and we employed the *p*^*act1*^ promoter to express the ParB^c2^ protein fused to mCherry and SV40 nuclear localization signal, followed by the transcriptional terminator of the budding yeast CYC1 gene. The obtained strains did not show gross defects in growth, mating and spore production. We relied on the clear signal of ParB^c3^-eGFP for subsequent quantifications because *parS*^*c2*^/ParB^c2^-mCherry-NLS markers were difficult to reliably discern from the diffuse nuclear staining. We noted that without the NLS, the ParBc2-mCherry was largely cytosolic.

To visualize **the nuclear and S-phase marker Pcn1**, we introduced a second copy of the gene at the *ura4* locus and fused the ORF with the N-terminal mCherry tag^13^ .

The **nuclear marker** ^**NLS**^**mCherry** consisted of the N-terminal SV40 nuclear localization signal^24^ PKKKRKV fused to the mCherry fluorescent protein and expressed from the *p*^*act1*^ promoter. The construct was integrated at the *ura4* locus.

The **mating type fluorescent markers** consisted of the fluorescent protein mTagBFP2 under the regulation of either the P-cell-specific *p*^*map3*^ promoter or the M-cell-specific *p*^*mam1*^ promoter, which we integrated at either *lys3* or *ade6* loci. We noted that both constructs showed high variability in mTagBFP2 production in response to nitrogen-starvation and mating. Additionally, homothallic cells occasionally induced mTagBFP2 expression and subsequently divided to produce daughters of opposite mating types, which both inherited mTagBFP2 from the mother. For these reasons, the mating type of homothallic cells could be reliably determined only by timelapse imaging of the mating process when the increase in mTagBFP2 occurs in only one partner.

To **label spores with fluorescent markers**, we used the *p*^*pdc202*^ promoter to drive expression of three tandem copies of either mCherry or mNeonGreen^7^ fluorescent proteins during sporulation. The constructs were integrated at the *leu1* genomic locus.

To **label cells with fluorescent markers**, we used the *p*^*tdh1*^ promoter to drive expression of either sfGFP or mCherry. The constructs were integrated at the *lys3* locus. The high levels of sfGFP and mCherry expression allowed us to detect fluorescence in cells, using fluorescence microscopy, and in colonies on solid media, using the transilluminator Safe Imager (Invitrogen, Waltham, USA) and the M165FC stereomicroscope (Leica, Wetzlar, Germany).

### Spore viability assay

To obtain **spores from zygotic asci**, we prepared mating mixtures of ∼2·10^7^ haploid cells on MSL-N solid media and incubated samples for 24 h at 25°C to allow mating and formation of spores. To obtain **spores from azygotic asci**, we spotted ∼2·10^7^ diploid cells on MSL-N solid media and incubated samples overnight at 25°C to allow sporulation. We transferred the samples to 1 mL of sterile water containing 2.5 μL of β-Glucuronidase/Arylsulfatase (10127060001, Roche, Switzerland) and incubated samples overnight at 30°C with agitation. We used the transmitted light microscope to confirm that the enzymatic treatment led to lysis of vegetative cells. We collected the spores by centrifugation at 1,000 g for 1 min and washed cells once in 1 mL of sterile water. We used the Neubauer chamber to count the number of spores in each sample and we plated a defined number of spores (500 to 1,000 spores per replicate) onto YES growth media. The plates were incubated at 30°C until colonies developed, and we report the spore viability as the percentage of observed colonies relative to the plated number of spores.

### Gamete viability assays

When assessing the viability of mated and unmated gametes, we used asynchronous mating mixtures of the fusion deficient *fus1Δ* strains, which form unfused partner pairs and also contain unmated cells. We measured either the viability of gametes in individual mating pairs and viability of gametes in mating mixtures.

We measured the **viability of gametes in individual mating pairs** to directly compare viability between P-gametes that carry either *rec8+* or *rec8Δ* alleles. We mixed an equal number of exponentially growing *h+ rec8+* and *h+ rec8Δ* cells before introducing twice the number of *h-rec8Δ* mating partners. The three strains carried the *fus1Δ* allele and distinct selection and cell fluorescent markers. We split this initial cell mixture into two fractions and used the first fraction to verify the composition of the initial cell mixture; Specifically, we immediately plated 100-500 cells onto non-selective YES plates and proceeded with incubation at 30°C until individual colonies developed. We determined the genotype of each colony using distinct antibiotic selection and fluorescent cell markers. The observed ratio in the numbers of colonies of each genotype was used to calculate the ratio of the three strains in the initial cell mixture. The second fraction of the initial cell mixture was washed three times in MSL-N media and we immediately spotted ∼2·10^7^ cells onto MSL-N solid media for 18 h at 25°C to induce mating. Next, we resuspended these mating mixtures in sterile water and transferred cells onto non-selective YES media plates. We immediately used the MSM 400 micromanipulator (Singer Instruments, Somerset, United Kingdom) to visually distinguish and isolate 160 of mating pairs, or 160 individual gametes, per replicate. Isolated cells were incubated at 30°C until colonies developed, and we used the distinct selection and fluorescent markers to determine each colony’s founder cells. Since the expected fraction of each genotype recovered from the mating mixture should be determined by the initial cell mixture composition, we report the **gamete viability** as percentage of observed relative to the expected number of colonies recovered from mating mixtures. To directly compare viability between M-gametes that either carry wild-type or lack the *rec8* gene, we used the same experimental approach but initially mixed an equal number of exponentially growing *h-rec8+* and *h-rec8Δ* cells before introducing twice the number of *h+ rec8Δ* mating partners. Since viabilities for genotypes used in both experiments showed no significant differences (p^Welch^>0.7), we show results of both experiments together.

To measure **viability of cells in mated and unmated populations**, we used exponentially growing *h+* and *h-* strains lacking the *fus1Δ* allele and carrying distinct selection and cell fluorescent markers. We used either individual strains to test viability of unmated cells, or we mixed an equal number of partner cells to test cell viability in mating populations. For these cell mixtures, we split it into two fractions and used the first fraction to verify the composition of the initial cell mixture; Specifically, we immediately plated 100-500 cells onto non-selective YES plates and proceeded with incubation at 30°C until individual colonies developed. We determined the genotype of each colony using distinct antibiotic selection and fluorescent cell markers. The observed ratio in the numbers of colonies of each genotype was used to calculate the ratio of the two strains in the initial cell mixture. The second fraction of the initial cell mixture was washed three times in MSL-N media and we immediately spotted ∼2·10^7^ cells onto MSL-N solid media for 24 h at 25°C to induce mating. Next, we resuspended these mating mixtures in sterile water, used the Neubauer chamber to count the number of cells in each sample and we plated a defined number of cells (500 to 1,000 spores per replicate) onto YES growth media. The plates were incubated at 30°C until colonies developed, and we report the spore viability as the percentage of observed colonies relative to the plated number of cells taking into account the ratio of the two strains in the initial mating mixture for mated gameets.

### Loss of Heterozygosity (LOH) assay

The loss-of heterozygosity (LOH) assay, which detects both copy number loss and copy number neutral loss of alleles, was adapted from previously published work^25^. As detailed above, we used protoplast fusion between haploid cells to obtain the *h+/h+* and *h-/h-* diploids that show gametic cell fate. Both copies of the *fus1* gene were replaced by the *bsdMX* selection cassette, which prevented cell fusion and conferred resistance to blasticidin-S. Additionally, the cells carried the *ade6-M210* and *ade6-M216* mutant alleles of the *ade6* gene, which are individually non-functional but capable of inter-allelic complementation. Thus, the heterozygous *ade6-M210*/*ade6-M216* diploids are adenine prototrophs and form white colonies. If either *ade6* allele is lost, clones become auxotrophs for adenine and accumulate adenine biosynthesis intermediates that give red/pink hue to colonies. We report **LOH rates** as the percentage of *ade6* auxotrophic clones relative to all clones.

To measure **LOH rates upon nitrogen-starvation but without mating induction**, we cultured *h+/h+* and *h-/h-* heterozygous diploids separately in MSL+N media to exponential phase, washed cells three times with 1ml of MSL-N media, resuspended samples in 3 ml MSL-N for 3-5 hours to final O.D._600_ = 1, and spotted ∼2·10^7^ cells of either strain on solid MSL-N media. After 24 h, the spotted cells were collected, resuspended in sterile water and we plated 10^-3^, 10^-4^ and 10^-5^ serial dilutions on solid YES medium containing blasticidin-S, which diploid strains are resistant to. We incubated the plates at 30°C until colonies formed, performed replica platting onto EMM medium lacking adenine and scored the number of adenine auxotrophic (red/pink) and prototrophic (white) colonies. To measure **LOH rates upon mating induction**, we processed the samples in the same manner with the exception that at the point of washing cells in MSL-N, we introduced the equal number of blasticidin-S-sensitive haploid *fus1Δ* (*patMX* or *natMX*) mating partner cells. After the 24 h incubation on MSL-N plates, the mating efficiency between diploid cells and haploid partners was similar in all crosses (p^Welch^ >0.4, average mating efficiency 81±2%). As previously, cells were resuspended in sterile water and we plated serial dilutions on solid YES medium containing blasticidin-S, which killed the haploid cells used to induce mating. The plates were incubated at 30°C until colonies formed, and we performed replica plating onto EMM medium lacking adenine before scoring for adenine auxotrophic and prototrophic colonies.

### Experimental evolution

We performed experimental evolution experiments as biological triplicates using heterothallic *h+* and *h-* strains that lacked the *fus1* gene, which prevented cell fusion. Cells also carried distinct antibiotic-resistance markers and expressed fluorescent cell markers, which allowed us to distinguish the two cell types in mixed populations. As indicated, the cells carried either wild-type or lacked the *rec8* gene.

The initial populations for evolution experiments were prepared by mixing equal number of heterothallic P- and M-cells. Each experimental cycle started by collecting exponentially growing cells, which we washed three times in MSL-N media and resuspended in 3 mL of MSL-N liquid media to O.D._600_ = 1, followed by incubation at 30°C 3 h with shaking at 200 rpm. We concentrated ∼2·10^7^ cells into 20 µL of media and spotted samples onto MSL-N agar plates. The cells mixtures were incubated 24 h at 25ºC, and entire populations resuspended in YES growth liquid media to final cell density O.D._600_ = 0.05. We incubated recovered populations at 30ºC until cells reached exponential growth, which took approximately 24 h. To initiate the next mating cycle, we collected mitotically growing cells, washed, pre-starved and spotted mixtures onto MSL-N media. Simultaneously, we assessed the composition of the evolving population by collecting 2·10^6^ of mitotically dividing cells and plating 10^-3^, 10^-4^ and 10^-5^ serial dilutions onto non-selective YES solid medium. These plates were incubated at 30°C until colonies developed. We used the transilluminator Safe Imager (Invitrogen, Waltham, USA) and the M165FC stereomicroscope (Leica, Wetzlar, Germany) to distinguish the fluorescent protein present in each colony and thus determine the strain they derived from. We validate these results by replica plating colonies on selective media containing distinct antibiotics. We report population composition as the percentage of colonies derived from each strain in the mixture relative to the total number of observed colonies.

### RNA extraction and high-throughput RNA sequencing

RNA-Seq experiments were performed as biological triplicates. We nitrogen-starved either homothallic *fus1Δ* mutants, which arrested as mated pairs, or heterothallic wild-type strains that produce only one gamete type and thus arrest as unmated cells. Cells growing exponentially in liquid MSL+N media were transferred to MSL-N liquid medium, diluted to density of O.D._600_ = 1 and incubated at 30°C with 200 rpm agitation for 5 hours. Samples were collected by centrifugation, ∼2·10^7^ cells concentrated into 20 μL of media and spotted onto MSL-N solid media. After 10 hours of incubation at 25°C, mating efficiency reached approximately 80% and we harvested 6 mating spots (∼1.2·10^8^ cells) for RNA extraction using inoculation loops.

RNA extraction was based on the previously published protocol^26^ with indicated modifications. Cells recovered from mating spots were resuspended in ice-cold DEPC-treated water, centrifuged at 1,000 g for 1 minute, the supernatant removed and cells snap-frozen in liquid nitrogen. Pellets were resuspended in 750 μL of extraction buffer (10 mM Tris-HCl, pH 7.5; 10 mM EDTA, pH 8.0; 0.5% SDS), mixed with an equal volume of acid-equilibrated phenol, and incubated at 65°C for 1 hour. The samples were cooled on ice and centrifuged at 16,000 g for 15 minutes at 4°C. The aqueous phase was sequentially extracted with phenol:chloroform:isoamyl alcohol (125:24:1; 77617-100ML, Merck, Darmstadt, Germany) and chloroform:isoamyl alcohol (24:1; 25666-100ML, Merck, Darmstadt, Germany), each using centrifugation at 16,000 g for 5 min at 4°C. RNA was precipitated at -20°C overnight using three volumes of absolute ethanol and 0.1 volumes of a 3M sodium acetate pH 5.2 solution, and recovered by centrifugation at 16,000 g for 10 min at 4°C. The RNA pellet was washed with 70% ethanol pre-chilled to 4°C and re-suspended in DEPC-treated water. The RNA samples were further purified and subjected to DNAse treatment using the Monarch RNA purification kit (T2010S, New England Biolabs, Ipswich, USA) following manufacturer instructions. RNA samples were stored at -70°C until use. For library preparation, we used the Illumina TruSeq Stranded mRNA kit. Sequencing was performed on a HiSeq 4000 platform, generating 150 bp single-end reads with a depth of approximately 10 million reads per sample, resulting in over 100X coverage. Sequencing results are available under GEO accession identifier GSE293837.

### RNA-Seq data processing and gene expression analyses

The raw FastQ files obtained from RNA-Seq passed the FastQC quality-check and underwent read trimming with Trimmomatic^27^ and alignment to the *S. pombe* 972 *h-* reference genome (ASM294v2.54) using HISAT2 v2.2.1^28^. Over 93% of reads aligned to the genome in all samples. The gene annotations were obtained from Ensembl Fungi and the differential expression analysis was performed using DESeq2 v1.24.0^29^. We considered as differentially expressed genes exhibiting a |log2(fold-change)| ≥1 with an adjusted p-value <0.05. Transcript abundance was quantified in transcripts per million

To **compare upregulated gene sets** reported in our and previously published studies^15,30,31^, we first curated all datasets. Specifically, we removed and updated obsolete protein-coding gene identifiers using the Pombase database as reference. For comparison with each study, we further curated the datasets to include only genes that were probed in both our and the study used for comparison. The curated lists contained more than 4365 of probed genes. We used the curated lists and applied the custom R scripts, built on the *tidyverse* package, to identify the overlap in upregulated protein-coding genes, and we report the hypergeometric test p-values for indicated overlaps between datasets.

**Gene ontology (GO) enrichment analysis** was performed using the FunRich^32^ (v.3.1.3) software for the protein-coding genes upregulated in mated relative to unmated gametes. The entire upregulated gene set was present in the UniProt^33^ *S. pombe* database, which served as the sample background for enrichment analyses. We report the 10 biological process GO terms with the most significant enrichment as calculated using hypergeometric test. The overrepresentation of genes from the specific GO category among upregulated genes is indicated by the fold-enrichment, which reports the fraction of GO category genes in the upregulated gene set relative to their fraction in the background gene set.

### Microscopy and image processing

We **mounted cells for microscopy** using previously reported protocol^11^ with indicated modifications. To monitor mating, exponentially growing cells were washed in MSL-N and, unless otherwise indicated, pre-starved in MSL-N liquid media for 3-5 h before being immobilized on MSL-N agar pads. To monitor gametes returning to mitotic proliferation, mating mixtures spotted on MSL-N agar plates were resuspended in sterile water and cells transferred onto YES agar pads. The samples were enclosed between a glass coverslip and either a standard glass slide, as previously reported^11^, or a custom PDMS mold, which will be reported separately. We note that cells placed in PDMS chambers yielded stronger fluorescence with reduced bleaching during timelapse microscopy, likely due to improved gas exchange compared to glass chambers. The chambers were sealed with VALAP, prepared as a 1:1:1 mixture of lanolin (A16902.30, Alfa Aesar, Haverhill, USA), Vaseline (16415, Merck, Darmstadt, Germany), and paraffin (26157.291, VWR, Radnor, USA). Imaging was performed in a temperature-controlled environment maintained at 25±2°C.

We used **transmitted light microscopy** to directly visualize and quantify frequency of visually distinct cell types including gametes, mating pairs, zygotes and spores. Observations were made on a Nikon Eclipse Si microscope (Nikon, Minato, Japan) equipped with 40X, 60X, and 100X objectives.

We performed **fluorescence microscopy** using two Nikon imaging systems, both built on the Nikon Eclipse Ti2-E inverted microscope with a TI2-S-SE-E motorized stage and a Z100-N piezo controller for Z-axis control. Samples were imaged with CFI Plan Apochromat Lambda 60X Oil 1.4 NA or CFI Plan Apochromat Lambda 100X Oil 1.45 NA objectives, and captured using a BSI Express camera (Teledyne Photometrics, Tucson, USA). Illumination for transmitted light images was provided by a pE-100 LED (CoolLED, Andover, UK), while fluorescence excitation was achieved using the Lumencore Spectra III 8-NIE-NA system (Beaverton, USA) with excitation lines at 390/22 nm, 440/20 nm, 510/25 nm, 555/28 nm, 575/25 nm, 637/12 nm, and 748/12 nm. Fluorescence filters and mirrors were sourced from Semrock (Rochester, USA), Chroma (Bellows Falls, USA), and Analysentechnik AG (Tübingen, Germany). Filter switching was achieved through either the Ti2-E integrated turret or an LB10-3 filter wheel (Sutter Instrument, Novato, USA). Imaging hardware was synchronized using a PCIe-6323 card with a 6323 trigger breakout box (National Instruments, Austin, USA). The Nikon Perfect Focus System PFS4 was employed for autofocusing. Image acquisition was controlled via Nikon NIS Elements AR software (v5.30.05). Micrographs within a single figure panel were captured with identical microscope settings for a given channel.

**Image processing** was conducted using ImageJ/Fiji (NIH, Bethesda, USA)^34^ and custom Python scripts utilizing *numpy* (https://github.com/numpy/numpy), *nd2* (https://github.com/tlambert03/nd2), and *tifffile* (https://github.com/cgohlke/tifffile) packages. Custom scripts are compiled into a GUI-based tool, *VLabApp*, available at https://github.com/vjesticalab/VLabApp. Brightfield micrographs represent a single focal plane, while fluorescence micrographs show either single z-sections or z-projections. Drift correction during timelapse imaging was performed by aligning samples along x,y axes using integer pixel shifts to maintain pixel values. Z-drift correction was based on transmitted light images to identify the central focal plane. Except in **Fig.3B**, green fluorescence micrographs were displayed with identical contrast within a figure panel.

### Quantifications of microscopy images

**Quantifications of fluorescence intensities** were performed using ImageJ/Fiji^34^, Cellpose^19^ and custom Python scripts and packages detailed bellow. Custom scripts are compiled into a GUI-based tool, *VLabApp*, available at https://github.com/vjesticalab/VLabApp.

We used transmitted light micrographs to **segment cells** either manually or using Cellpose^19^ segmentation tool, which we previously trained on our datasets. We used nuclear markers ^NLS^mCherry or mCherry-tagged Pcn1 to **segment nuclei** either manually or using custom Python scripts built on the *skimage* package (https://scikit-image.org), which employed absolute and adaptive contrasting to identify nuclear boundaries. The **cytosol regions** were determined as cellular regions excluding the nuclear regions. We **determined the sample background areas** either manually as regions unoccupied by cells, or automatically as the least intense 50 pixels in the microscopy channel and the timepoint of interest. We note that **we visually curated the results of computational segmentation** to ensure quality of subsequent quantification. We used segmented regions (e.g. cells, nuclei and cytosols) to quantify **mean fluorescence intensities** and report values after sample background was subtracted.

To determine the **time of cell fusion** we visualized cytosolic mixing of mTagBFP2, which we expressed in only one partner gamete. We used timelapses and determined the time of fusion either manually or using custom Python scripts built on *numpy* package. For the later, we calculated the standard deviation of all pixel intensities in paired gametes and zygote regions at each timepoint. We then defined the fusion as the point where the change in the standard deviation of pixel intensities between consecutive timepoints is largest, which coincides with the spread of the cytoplasmatic fluorophore from one gamete to its partner. We used the fusion time to **align timecourses** from different mating events. The onset of detection for proteins of interest, temporal dynamics of fluorescence intensities and onset of S-phase are all reported relative to cell fusion.

To determine the **time of S-phase onset** relative to gamate fusion, we used timelapse microscopy and monitored the mCherry-tagged DNA replication fork component Pcn1, which forms prominent foci only during DNA replication. We considered zygotes as undergoing replication when we observed multiple Pcn1 foci for at least 20 consecutive minutes. We report the mean time from fusion until first mCherry-Pcn1 foci form.

To quantify ***rep1 and rec8* transcription dynamics**, we manually scored when the PP7 reporter fluorescence dot first appeared in P- and M partners that fuse and report the mean onset of reporter detection. In case of the *rep1* reporter, we additionally scored whether the PP7 reporter produced a fluorescence dot in each partner for 10 min intervals in the 3 h preceeding fusion, and we report the average frequency of reporter detection for each partner at all timepoints.

To quantify **3sfGP-NLS expression from the *p***^***rep1***^ **promoter**, we we used custom python scripts to automatically measure the mean cellular prep1-3sfGFP-NLS fluorescence. We used the timepoints prior to rep1 promoter induction to quantify the cellular background signal and aligned measurements from multiple mating events by the time of gamete fusion.

To quantify the **time of induction** for the sfGFP tagged endogenous Rec8, Rep1, Rec11 and Rec25, we manually determined the earliest timepoint that gametes produced nuclear fluorescence above the background signal and report it relative to the time of partner fusion.

To quantify **Rec8 expression dynamics**, we used custom scripts to segment the mCherry-Pcn1 labeled nuclei and we report the mean nuclear Rec8-sfGFP fluorescence for the indicated number of mating events aligned by the time of fusion. We chose to quantify the nuclear signal because Rec8-sfGFP is strongly targeted to the nucleus, and because the low background of the nucleus allows more reliable signal detection. We used the timepoints prior to Rec8 induction to quantify the nuclear background signal. To quantify the Rec8-sfGFP production dynamics, we calculated the Rec8 production rate *v* and onset of production (*t*_0_) by fitting the Rec8-sfGFP intensity curves from mating events to the model *I*_*Rec*8_(*t*) = maximum(*b, v* ∗ (*t* − *t*_0_) + *b*) where *I*_*Rec*_8(*t*) is the signal intensity at time *t* and *v* is the baseline intensity. We used the “Interior Point Newton” method from the “Optimization.jl” Julia package to minimize a mean squared error loss function.

To quantify **Rec8 levels at the onset of S-phase**, we used the mCherry-Pcn1 marker to visualize the onset of S-phase. We manually segmented the zygotes using brightfield images and report the mean cellular Rec8-sfGFP fluorescence intensity above sample background.

To quantify the **frequency of gametes that induce Rec8 but fail to fuse with a partner**, we used homothallic strains with endogenous *rec8* fused to sfGFP. We used timelapse microscopy to visualize **uninterrupted matings** over 24 h. We manually scored for Rec8-expressing gametes that aborted mating, and report their percentage relative to all gametes that mated. We note that gametes that commenced mating in the last 40 minutes of the timelapse were excluded from the analyses. Alternatively, **we interrupted mating** — whereby mating mixtures residing on MSL-N agar plates for 6-8 h at 25°C were transferred to imaging chambers with YES growth media — and we used brightfield images to visually distinguished gametes from zygotes based on cell size and morphology. We report the frequency of unfused gametes with detectable Rec8-sfGFP as percentage of all cells. Furthermore, we report the **levels of mTagBFP2 expressed from the *p***^***mam1***^ **promoter** in homothallic gametes whose mating was interrupted. Here, we used 30 gametes with detectable Rec8-sfGFP as the Rec8-positive population. The control Rec8-negative population of 120 gametes included four gametes that were closest to each Rec8-positive gamete and did not show Rec8-sfGFP fluorescence. We report the mean cellular mTagBFP2 fluorescence above media background for both Rec8-positive and Rec8-negative populations.

To determine the **mating type of gametes that induce Rec8 but fail to fuse with a partner**, we interrupted mating between heterothallic strains where only one partner expresses the red fluorescent marker ^NLS^mCherry. We mixed equal number of partner cells and induced mating on MSL-N agar plates for 6-8 h at 25°C before transferring samples to imaging chambers with YES growth media. We used brightfield images to visually distinguished gametes from zygotes based on cell size and morphology. Both strains carried endogenous *rec8* fused to sfGFP, which allowed us to manually identify gametes that induce Rec8-sfGFP and to determine their mating type using the _NLS_mCherry marker. The percentage of P- and M-cells in Rec8-expressing gamete population is reported separately for crosses where *h+* or *h-* strain carried the ^NLS^mCherry marker, while the statistical significance for the overrepresentation of P-cells in the Rec8-expressing gametes population was calculated using data from both crosses.

To assess the **viability of gametes that either induce or lack Rec8 expression** we interrupted the mating of homothallic cells that carry endogenous *rec8* fused to sfGFP. Specifically, cells mating on MSL-N agar plates for 6-8 h at 25°C were transferred to imaging chambers with YES growth media and imaged over 24 hours. We used brightfield images to visually distinguish gametes from zygotes based on cell size and morphology, and for each gamete we manually determined whether Rec8-sfGFP signal is induced or not at the start of imaging. Each gamete was then manually tracked for up to two mitotic divisions and we scored instances of cell lysis and growth arrest. For the Rec8-positive and Rec8-negative gamete subpopulations, we report the percentage of gametes that themselves or their daughters lysed or arrested growth.

To quantify **Rep1 protein expression dynamics**, we used custom python scripts to automatically measure the mean cellular Rep1-sfGFP fluorescence. We used the timepoints prior to Rep1 induction to quantify the cellular background signal and aligned measurements from multiple mating events by the time of gamete fusion.

To quantify **Ste11 protein expression dynamics**, we manually segmented cells and we report the mean cellular Ste11-sfGFP fluorescence above background fluorescence. To quantify the **nucleocytoplasmic ratio of Ste11**, we manually segmented the nuclei using the signal of red fluorescent marker ^NLS^mCherry, measured the Ste11-sfGFP in the nucleus and the cytosol and we report their ratio. Measurements from multiple mating events were aligned by the time of cell fusion.

To quantify the accuracy of **meiotic chromatin segregation**, we used timelapse microscopy and monitored dynamics of the Hht1 histone H3 tagged with RFP. We consider as accurate meiotic segregation instances where the zygotic Hht1-RFP signal divided into two visually similar masses at meiosis I, and each of those then divided into two visually similar masses at meiosis II. Any deviation from this pattern was considered as a chromatin mis-segregation defect. Specifically, we observed zygotes where at least one meiotic division failed and instances of meiosis I or II that divided chromatin into two or more clearly uneven chromatin masses. These defects generally resulted in production of fewer or more than four chromatin masses at the end of meiosis. We report the percentage of zygotes showing chromatin segregation defects relative to all analyzed zygotes.

To quantify the accuracy of **meiotic allele segregation**, we monitored the segregation of two synthetic alleles of the same locus on chromosome II between spores. Specifically, we imaged sporulated zygotes obtained by crossing heterothallic strains where the *leu1* locus was linked to the sporulation-induced *p*^*pdc202*^ promoter that regulates 3mNeonGreen in the *h-* strain, and 3mCherry in the *h+* strain. Mendelian segregation of the two alleles results in two spores that express 3mNeonGreen and two spores that express 3mCherry, which we report as accurate allele segregation. We classified instances where at least one spore displays both fluorescent signals as meiotic mis-segregation of alleles. We report zygotes that produced aberrant spore number as a separate class because they may arise from meiotic chromosome mis-segregation or other defects. We report the percentage of the three types of zygotes relative to all zygotes visualized in the cross.

To quantify **sister chromatid cohesion** we, visualized either the P- or the M-parental centromere-distal lys3 or the centromere-proximal rec6 loci with the green fluorescent *parS*/ParB reporter, as indicated. From the time of partner fusion until meiotic divisions, in 10 min intervals we searched for instances of two labeled foci and report the maximum distance between them. We additionally report instances where *rec6* labled loci on sister chromatids co-segregated together or separated to different zygote poles as **frequencies of reductional and equational meiosis I**, respectively. Additionally, we monitored zygotes with both parental *rec6* alleles fluorescently labeled and we report instances of four loci co-segregating at meiosis I as **frequency of homologue non-disjunction**.

We quantified **mating and sporulation efficiencies** from mating mixtures spotted on solid MSL-N media and incubated at 25°C for approximately 24 hours, unless otherwise indicated. We used transmitted light microscopy to visually distinguish and score the number of gametes, unsporulated zygotes and sporulated zygotes. We report results as percentages calculated using the following formulas:

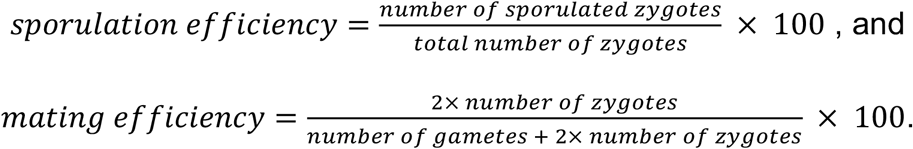

## Statistical analyses

Statistical analyses were performed in Microsoft Excel, R, Julia and Python software packages. We report the sample size (n) and number of biological replicates (N) in figures and figure legends and note that sample sizes were not predetermined. To pool data from multiple mating timelapses, we used the time of cell fusion to align measurements. No randomization or blinding was used. Any sample exclusion is detailed in the result.

As indicated in figure legends for graphs, we report **mean values** and **standard deviation** intervals or, in case of binomial data, the **Wilson interval**.

**Error propagation** for z±σ_z_ calculations from a±σ_a_ and b±σ_b_ was performed for addition and subtraction as σ_z_=sqrt(σ_a_^2^+σ_b_^2^) and for multiplication and division as σ_z_=z·sqrt((σ_a_/a)^2^+σ_b_/b)^2^).

**Orthogonal regression** coefficients were obtained by computing the slope of the first principle component, using the PCA method from the Julia package “MultivariateStats.jl”.

p-values from the **Welch test** — an unequal variances t-test for two-tailed distribution — were calculated in MS Excel using TTEST function. p-values from the **Kruskal-Wallis test** — a non-parametric test for two or more samples — were calculated using *scipy*.*stats* Python package function *kruskal*. p-values from the **Dunn test** — multiple comparisons, post hoc, non-parametric test — were calculated following an initial Kruskal-Wallis test for significant differences and using *scikit_posthocs* Python package function *posthoc_dunn* with Bonferroni adjustment implemented using the *holm* parameter. p-values from the **Tukey’s Honest Significant Difference test** — a multiple comparisons, post hoc test based on the studentized range distribution that inherently includes correction for multiple comparisons — were calculated following initial ANOVA test for significant differences and using *scipy*.*stats* Python package function *tukey_hsd*. p-values from the exact McNemar test — a test for paired binomial data — were calculated using *statsmodels* Python package function *mcnemar*. We report the p-values from the **hypergeometric test**, which tests whether sub-populations are over- or under-represented in a sample, for the overlap of upregulated genes reported in this and previous studies, and for the Gene Ontology (GO) enrichment analyses.

**Boxplots** were obtained from *matplotlib*.*pyplot* Python package with the *boxplot* function. **Stacked histograms** and barplots were obtained from *ggplot2* R package.

## Supplemental Note S2

### Summary

In the isogamous fission yeast *Schizosaccharomyces pombe*, partner P- and M-gametes differentially invest into zygotic development by making unequal amounts of the conserved meiotic cohesin Rec8. Driven by asymmetric partner communication, P-gametes produce Rec8 before fertilization to promote meiotic fidelity and production of viable spores in zygotes. However, this early Rec8 expression also causes genomic instabilities and imposes a fitness cost on P-gametes [1], revealing a trade-off between sexual success and gamete survival (see main text for details).

Our model setup aims to explore how partner gametes distribute production of meiotic factors during sexual reproduction. In particular, we are interested in cases where the meiotic factor benefits zygotes - by promoting successful meiosis and hence production of viable spores - but introduces a cost for gametes that do not successfully find a partner and re-enter a vegetative, mitotic growth phase upon nutrient re-supply. We aim to explore how the costs to gametes and benefits for zygotes of such a meiotic factor quantitatively determine the optimal evolutionary strategy for the production of the meiotic factor in partner gametes.

Here, we 1) provide details of the simulations performed in the main text with regards to the questions posed above, 2) offer further explanation and analysis on the bifurcation arguments presented in the main text, and 3) describe the statistical approach used to fit model parameters to experimental data.

#### 1 Evolutionary model, simulations and theory

##### 1.1 Evolutionary model

Populations of fission yeast cells reproduce asexually through mitosis when nutrients are readily available. However, upon environmental induction, such as nitrogen depletion, cells differentiate into gametes that undergo sexual reproduction. When a pair of gametes successfully fuses, they produce a zygote that enters meiotic divisions and sporulation to produce 4 spores. Meiotic success - and consequentially the viability of spores - depend on the total Rec8 levels provided by both parental gametes to the zygotes at the time of fusion.

In some instances, gamete fusion is unsuccessful. Such unfused gametes may attempt to return to mitosis, however, in anticipation of fertilization, they have produced a quantity of Rec8. Rec8 expression is associated with genomic instability in mitotic cells [1], and hence introduces a fitness cost for gametes returning to asexual reproduction.

The described effects of Rec8 on gametes and zygotes introduce two opposing Rec8-dependent selection pressures on populations of cells undergoing cycles of sexual and asexual reproduction. In what follows, we describe an agent-based model that implements these selection pressures whilst allowing the amount of Rec8 produced by each gamete to evolve.

A population of *L* individuals is initialized with an identical genotype 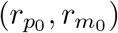. As reported in the main text, when reproducing sexually, mating partners produce Rec8 prior to fusion, and the genotype (*r*_*p*_, *r*_*m*_) defines the quantity of meiotic factor provided at the point of fusion by the P- and M-gametes, respectively. We assume that *r*_*p*_ and *r*_*m*_ are bounded individually between 0 and *r*_max_, from which it follows that 0 ≤ *r*_*p*_ + *r*_*m*_ ≤ 2*r*_max_. As a cell *i* undergoes mitosis its genotype 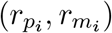 is mutated through the following rule: 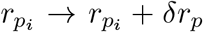 and 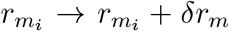, where 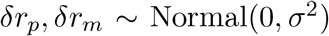. If through mutation *r*_*p/m*_ > *r*_max_ then we set *r*_*p/m*_ = *r*_max_ and apply the same logic to enforce non-negativity, *r*_*p/m*_ > 0. During mitosis a cell gives birth to two daughter cells, where we simplify the mating-type switching process [2] and assume that one of the sisters is assigned the P-gamete fate whilst the other has the M-gamete fate.

We define a maximum population size *L*_max_, and hence set the initial population size to be *L* = *L*_max_. It is assumed that the population undergoes *N* rounds of mitosis before individuals are induced to undergo gametogenesis. To avoid the population size growing exponentially, we force the size of the population to be equal to *L*_max_ after every round of mitosis: Suppose the size of the population before mitosis is *L*. In the case where the population size 2*L* after undergoing mitosis is smaller than the maximum population size *L*_max_, we duplicate (sample with replacement) some proportion of the population. In the case where it is larger than the maximum population size *L*_max_, we select *L*_max_ random individuals (sampling without replacement).

During gametogenesis, individual cells differentiate into P- or M-gametes depending on the mating-type assigned at birth. Once gametes are produced, an amount of the factor given by *r*_*p*_ and *r*_*m*_ is active in the P- and M-gamete, respectively. After gametogenesis, we have a population of *L*_max_ gametes (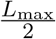 sister pairs) which we denote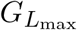 . In the simulations presented in this work, it is assumed that an individual always fuses with the sister. We introduce a parameter *f* ∈ [0, 1], which represents the probability that a given **pair** of sisters fuse successfully. The number of sister pairs which do not fuse is therefore distributed as *k* ≥ Binomial 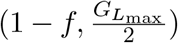. This determines a set of individuals 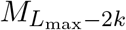 that successfully fuse and a set of individuals *G*_2*k*_ that remain as gametes and return to mitosis.

Rec8 is not the sole factor that regulates meiotic success and spore viability, and thus its quantitative effects might be modulated by non-specified, variable factors. For example, Rec8 requires other cohesin complex components and loading factors. To account for such complexity, we assume that there are preconditions for Rec8-mediated roles during meiosis. We assume that these conditions are met with some probability *κ*_*m*_. Given that these conditions are met and Rec8 is operational, we assume that the likelihood of meiotic success is then an increasing function of the total Rec8 concentration *r*_*p*_ + *r*_*m*_, and is given by a Hill function 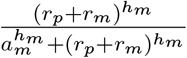 where we have assumed without loss of generality that gamete *i* is of type P and gamete *j* is of type M. Here, the parameter *a*_*m*_ acts as a location parameter, adjusting the concentration for which the probability saturates at 0. *h*_*m*_ is the Hill coefficient, which determines how quickly saturation is achieved at intermediate values.

Rec8-mediated meiotic regulation is non-operational with probability 1 − *κ*_*m*_, and in these instances we assume there is some background probability of successful meiosis Δ:_*m*_, which thus occurs in absence of Rec8. Consistent with this notion, meiosis in *rec8* Δ:/*rec8* Δ: zygotes produces some viable spores (Fig. 2A in the main text).

Given these definitions, a pair of sister gametes (*i, j*) from 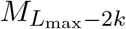 with genotypes 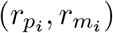 and (*r*_*p*_, *r*_*m*_) succeed in meiosis with a probability *P*_*i,j*_, which takes the following form assuming w.l.o.g that gamete *i* is a P-gamete and gamete *j* is a M-gamete:

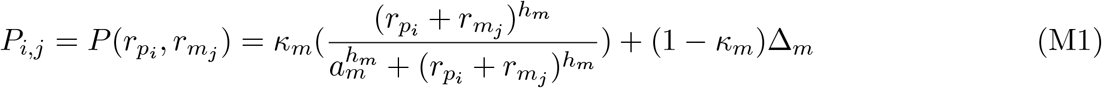

If meiosis is not successful, the progeny is assumed to die. If meiosis is successful, the zygote produces 4 spores that are assumed to inherit the genotype 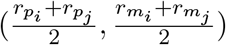, which is consistent with polygenic regulation of Rec8 expression levels.

For individuals in *G*_2*k*_, which remain as gametes and may return to mitosis, we assume that the conditions for Rec8-induced mitotic chromosome missegregation are met with some probability *κ*_*q*_. In these cases, the likelihood that a mitotic daughter cell of a gamete is viable is a decreasing function of *r*_*p*_ or *r*_*m*_, and is given by 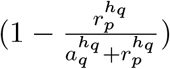. As previously, in cases where Rec8 does not influence viability of the offspring, we assume there is a background probability of survival Δ:_*q*_.

A gamete *i* from *G*_2*k*_ with amount of Rec8 equal to 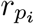 or 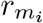 therefore produces viable daughter cells with probability *q*_*i*_, where *q*_*i*_ is given by the modified Hill function:

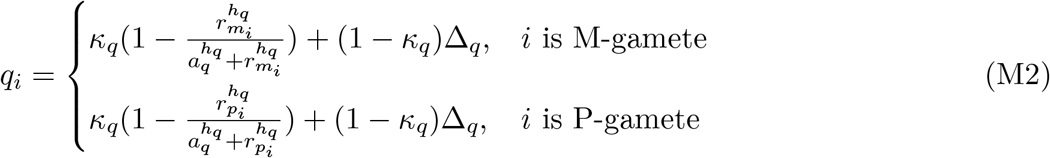

**Table M1:**
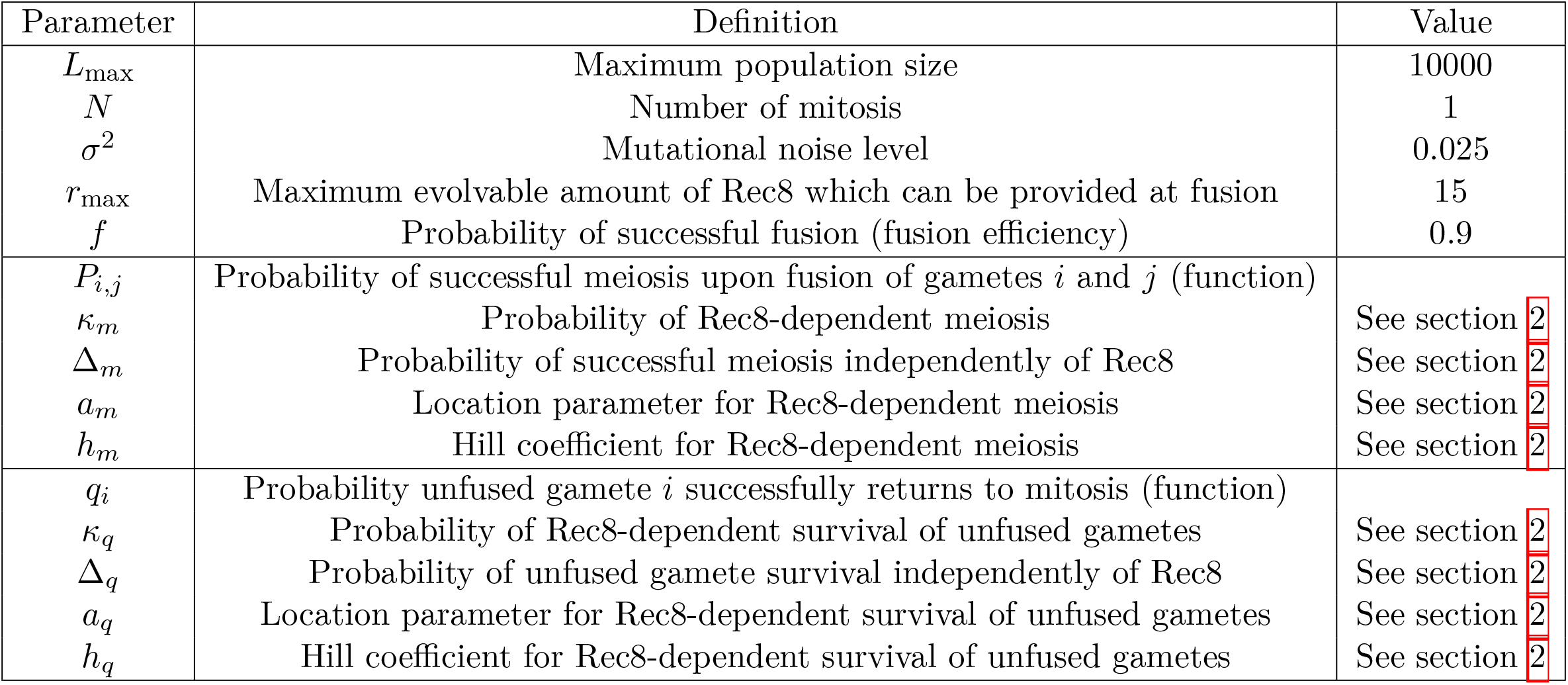
Model Parameters. A table specifying key parameters in the evolutionary model.

The surviving gamete daughters and spores then form the population that re-enters mitosis, and a single generation in evolutionary time has passed. The population then evolves for a number of generations until the population reaches a mutation-selection balance, where the average (*r*_*m*_, *r*_*p*_) no longer changes over evolution time.

Above, we have assumed that *P* and *q* follow sigmoidal profiles defined by Hill functions. We later verify this assumption by quantifying *P* and *q* experimentally. Key model parameters are given in table M1 and pseudo-code for the population-based life cycle model is given in algorithm 1.

In our analysis, we use both population-based simulations (as defined in Section M1, 1) and an analytical function to investigate the evolutionary dynamics associated with the fission yeast life cycle (as defined in Section 1.2).

**Figure M1:**
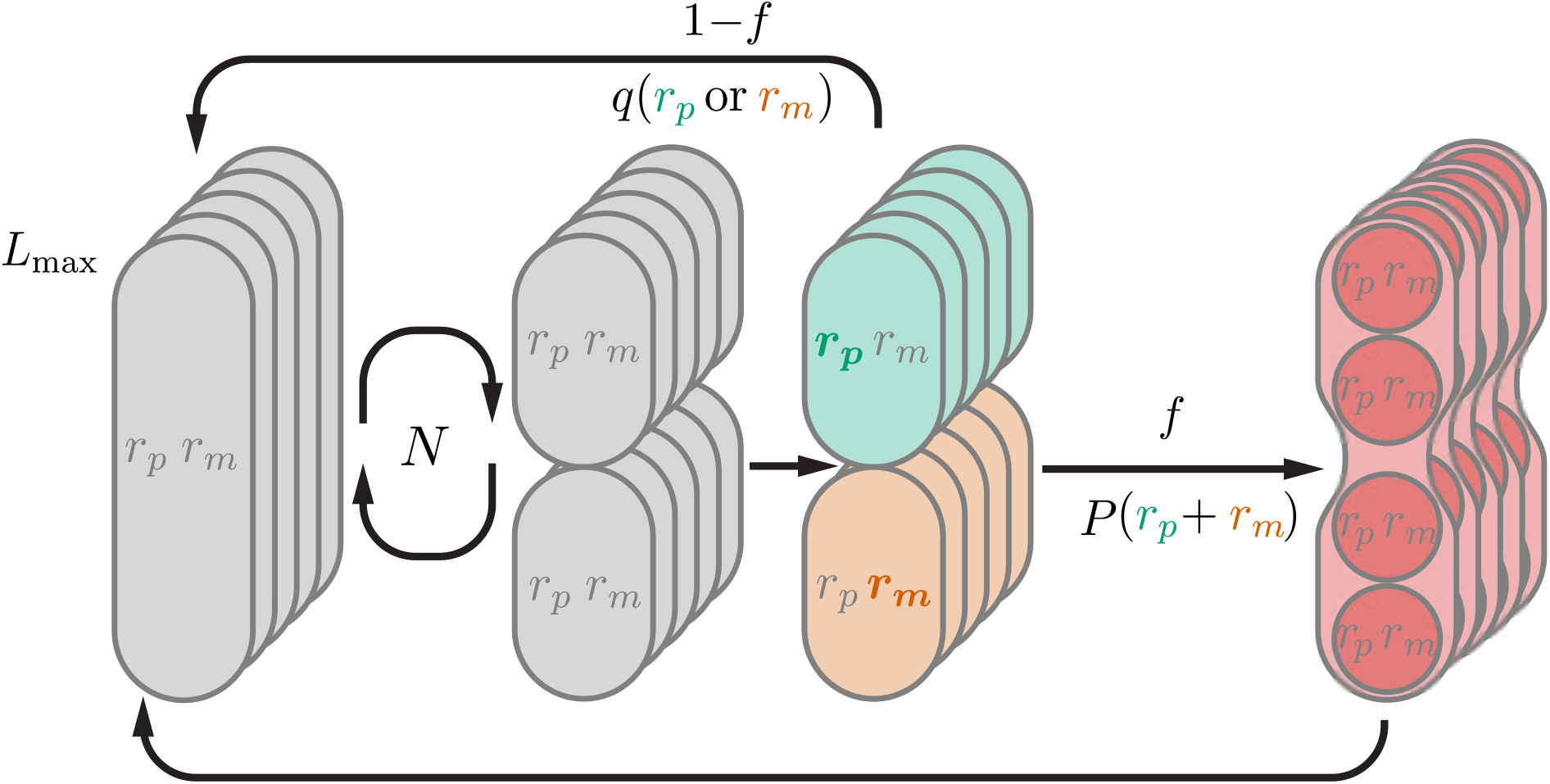
Fission Yeast Life Cycle. The schematic shows the life cycle stages represented in the model. There is a population of *L*_max_ cells. Each cell is associated with an evolving genotype (*r*_*p*_, *r*_*m*_) that defines the production of the meiotic factor in the P- and M-parent. Cells proliferate mitotically for *N* generations followed by a sister cells mating step. We introduce the probability *f* that partners fuse during mating. If fusion is successful, the zygotes receive *n* = *r*_*p*_ + *r*_*m*_ investment and sporulate with probability *P* (*n*). Non-fused gametes return to mitosis with probability *q*(*r*_*p/m*_) and die otherwise.

##### 1.2 Numerical analysis of evolutionary dynamics

We ran simulations based on the life cycle and evolutionary model described in the previous section. We assign an initial genotype of 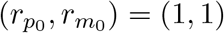 to each individual in a population of size *L*_max_ = 10, 000. We set the mutation size parameter *σ*^2^ = 0.025 and have *N* = 1 of mitosis before meiosis. We ran the evolutionary simulations for max_gen_ = 20, 000 generations. At each generation, we recorded each genotype 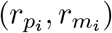 within the population. This allows us to compute the average genotype 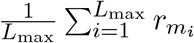 and 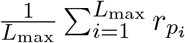. These population-based simulations and quantifications were used to generate Fig. 4B in the main text.

Note that the model does not have any features that intrinsically bias asymmetry in Rec8 production evolving in P-cells in preference to M-cells. However, without loss of generality, we denote the partner that evolves higher levels of Rec8 at the end of the simulation as *r*_*p*_.

###### Algorithm 1

Evolutionary model

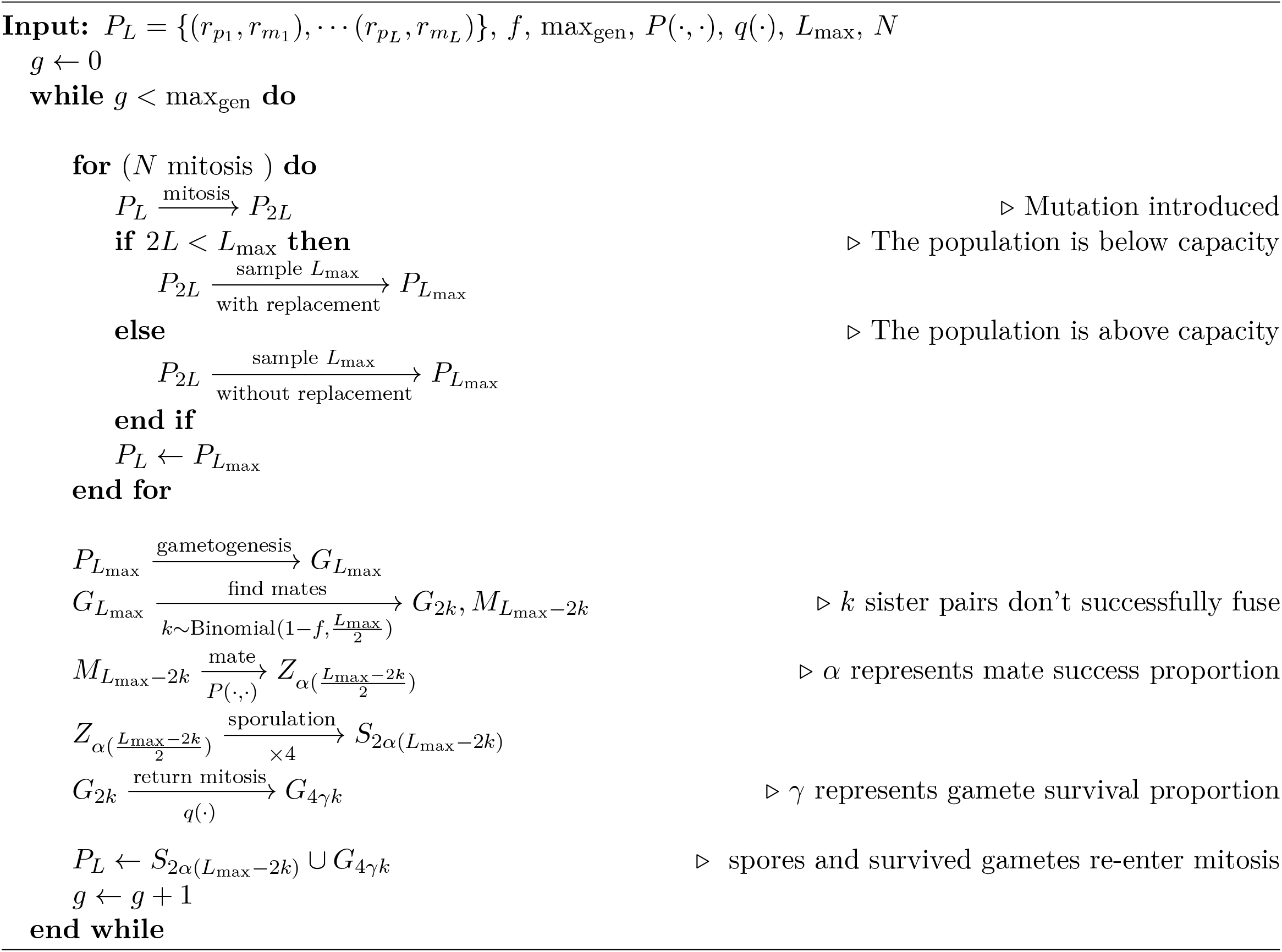

##### 1.3 Expected number of offspring upon sister mating

To further understand the parameters that determine outcomes of our numerical simulations, we derived an analytical expression for the expected fitness of a given genotype (*r*_*p*_, *r*_*m*_). When mating occurs exclusively between sister cells, it is possible to derive an analytical expression for the expected number of offspring that originates from a genotype (*r*_*p*_, *r*_*m*_) after one life cycle in our model. Note first that sisters *i* and *j* are assumed to be genetically identical and thus have identical genotypes, 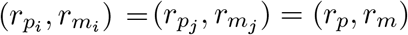. Furthermore, in our model, sisters always represent opposite mating types. Hence, we can write the expected number of offspring which will inherit the genotype (*r*_*p*_, *r*_*m*_) carried by two sisters given that they attempt to mate:

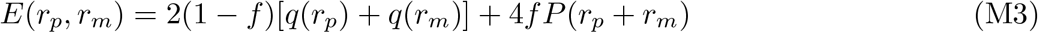

This expression is a proxy for the fitness associated with a genotype. Fitness is therefore dependent on two distinct contributions; the expected payoff if fusion is unsuccessful 2[*q*(*r*_*p*_)+ *q*(*r*_*m*_)] and the expected payoff if mating occurs, 4*P* (*r*_*p*_ + *r*_*m*_), with the fusion efficiency *f* determining which payoff is expected to dominate the other. Note that here the pre-factors reflect that mitotic division gives rise to two cells and meiosis gives rise to four cells.

###### 1.3.1 Analysis of system optima and bifurcations

Since the population size in our simulation is assumed to be large, the effects of drift are negligible, and we expect the solutions obtained at mutation-selection balance to correspond to those that maximize fitness Eq.M3. We use this correspondence to obtain further insight into the evolutionary dynamics leading to asymmetry, by determining numerically the values of (*r*_*p*_, *r*_*m*_) which maximize the fitness Eq.M3. For a specific choice of parameters defining *P* and *q* and for a given value of *f* we obtain a fitness function which assigns a fitness *E*(*r*_*p*_, *r*_*m*_) to a genotype (*r*_*p*_, *r*_*m*_). We then obtain an optimum genotype 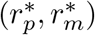 by optimizing this function numerically over the region [0, *r*_max_] *θ* [0, *r*_max_] using the Broyden-Fletcher-Goldfarb-Shanno (BFGS) algorithm, which is implemented in the Julia package `Optimization.jl` in the “BFGS” routine [3].

Additionally, we are able to study the bifurcation behavior of Eq.M3. In the main text we gain insight into the evolutionary dynamics by analyzing Eq.M3 for fixed total investment *n*. We can denote the fixed parental investment *n* = *r*_*p*_ + *r*_*m*_ and then consider

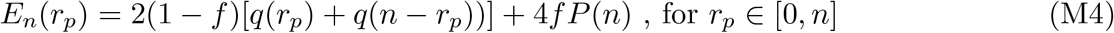

We can find the value 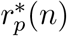 which maximizes *E*_*n*_ over the interval [0, *n*], and then 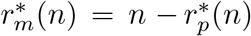. To find the maxima of *E*_*n*_(*r*_*p*_) over [0, *n*] we use the Broyden-Fletcher-Goldfarb-Shanno (BFGS) algorithm which is implemented in the Julia package `Optimiz ation.jl` in the “BFGS” routine [3]. This decomposition is shown in Fig.4D in the main text for varying *f* .

Crucially, we found that there exists a critical value *n*_*c*_, such that for *n*> *n*_*c*_ the optima for the function 2[*q*(*r*_*p*_)+ *q*(*n* − *r*_*p*_)] on the interval [0, *n*] transitions from symmetric optimum 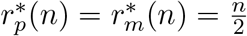 to an asymmetric optimum 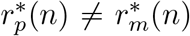 (see main text and Fig. 4E). To numerically identify this critical point, we optimize 2[*q*(*r*_*p*_) + *q*(*n*_*i*_ − *r*_*p*_)] subject to *r*_*p*_ ∈ [0, *n*_*i*_] for 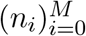 where 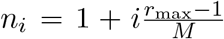. In our analysis, we set the number of discretization points *M* = 280. For each *n*_*i*_ we therefore obtain an optimum 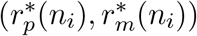. We define the critical value 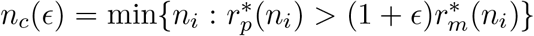. In our analysis, we set *ϵ* = 0.5.

The emergence of this bifurcation is a consequence of the non-linearity of the function *q*. First, if we assume *q* is linear in the range [0, 2*r*_max_] then there is no fitness advantage associated with asymmetry:

Assume *q*(*r*) = *a* + *br*, with *b*< 0,*a* > 0 so that *q* is a decreasing function of *r*. Then:

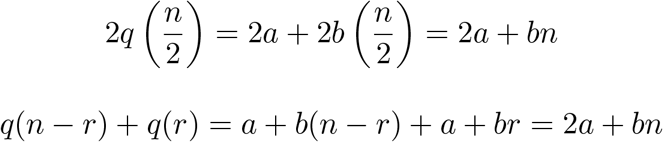

Thus:

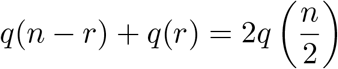

In other words, 2[*q*(*r*_*p*_)+*q*(*n*−*r*_*p*_)] is constant ∀ *n* ∈ [0, 2*r*_max_]. Hence, in the case of a linear relationship between *q*(*r*) and *r*, there is no selective advantage associated with asymmetry.

Suppose alternatively a Hill coefficient of 2, so that 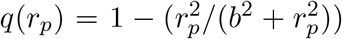. Let *g*(*r*) = *q*(*r*_*p*_) + *q*(*n* − *r*_*p*_). The function *g* is symmetric around *r*_*p*_ = *n/*2, and hence *r*_*p*_ = *n/*2 corresponds to a local minima or local maxima of *g* on the interval [0, *n*]. If *r*_*p*_ = *n/*2 is a local minima, then as a consequence of the extreme value theorem, there exists an alternative point *r*_*p*_ ∈ [0, *n*] with *r*_*p*_ = ≠ *n/*2 which is a global maxima for *g* on [0, *n*]. We can show there exists values of *b, n* such that *r*_*p*_ = *n/*2 is a local minima by showing that *g*^*”*^ (*n/*2) > 0 for some *n, b* > 0. We have that

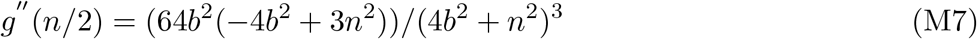

For *n, b* > 0, this is positive for 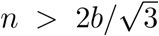. Hence, if this condition on the total investment and Hill function location parameter *b* holds, we have shown that *r*_*p*_ = *n/*2 is a local minima and therefore cannot be a global maxima on [0, *n*]. Therefore, there exists some 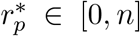 which is a global maxima with 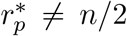, and consequently there exists optimal allocation strategies which are asymmetric for specific values of *b, n* > 0. Note that this argument holds for survival functions *q* of the form 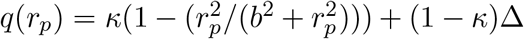 (as in Eq.M2) since the additional terms do not affect the symmetry of *g* or the positivity condition derived for its second derivative. Alternative Hill coefficients would yield different positivity conditions, yet the argument for the existence of non-symmetric allocation strategies through this non-linearity would remain the same.

We have shown that the bifurcation in the optimal strategy 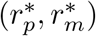 from a symmetric to asymmetric allocation is dictated by the total investment *n*^*\**^ and the shape of the survival function *q*. What dictates the optimal investment *n*^*\**^?

We can re-frame maximizing our fitness function (M3) as finding the optimal total investment *n*^*\**^ for the function 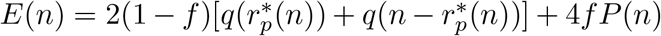, where 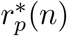 maximizes *E*_*n*_(*r*_*p*_) over the interval [0, *n*] as described above.

We note that the mating payoff *P* (*n*) selects for an increase in the total investment of meiotic factor *n* (Fig.4A, Fig.4G). Furthermore, we see that the maximal payoff 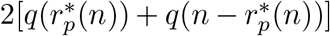 selects for a decrease in the total investment *n* (maxima decrease as *n* increases; main text and Fig.4E). Hence, there exists an optimum *n*^*\**^(*f*) ∈ [0, 2*r*_max_] which represents the optimal trade-off between these two competing selection pressures. In cases where continuing to increase the total investment *n* supports gains in fitness (since the benefit associated to meiosis outweighs the penalization to gamete survival), the system can in principle be driven to a high level of asymmetry if *n*^*\**^ >> *n*_*c*_. Therefore, the degree of asymmetry that evolves primarily depends on the limiting value of *n*, for which the increase in the expected meiosis success 4*P* (*n*) outweighs the decrease in the gamete mitotic success 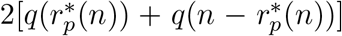. This limiting value depends specifically on the shapes of the functions *P* and *q* and the mating efficiency *f*, as well as the maximum possible Rec8 expression, *r*_max_ (since *n* ≤ 2*r*_max_).

### 2 Estimation of model parameters

This section outlines the inference framework we use to estimate the parameters for the functions *P*_*m*_ and *q*.

#### 2.1 Bayesian framework for parameter inference

We adopt a Bayesian framework for inference of parameters *θ* = (*h*_*m*_, *a*_*m*_, Δ:_*m*_, *κ*_*m*_, *h*_*q*_, *a*_*q*_, Δ:_*q*_, *κ*_*q*_). Our aim is to produce a posterior parameter distribution ℙ (*θ* |*D*) given experimental observations *D* and an initial prior distribution ℙ (*θ*), using Bayes formula:

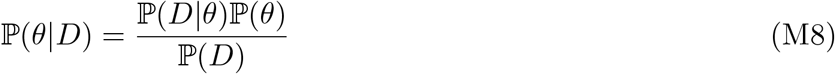

The quantity ℙ(*D*| *θ*) is known as the likelihood function in frequentist frameworks. The most common inference procedure in the frequentist framework is maximum likelihood estimation, in which

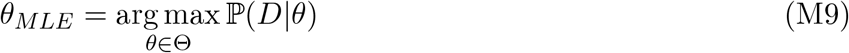

is the chosen parameter inference. This is known as a point estimate of the true, and unknown parameter set *θ*, and is itself a random quantity since it is derived from observations *D*, which are assumed to have a random component (e.g. measurement error or physical noise). In other words, a repeat experiment *D* ^*′*^ would likely yield an alternative point estimate 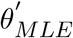. Similarly, confidence intervals constructed from observations *D* are also random quantities, and are constructed in a way which aims to encapsulate the true parameter *θ* with some probability (or “confidence”).

In Bayesian inference it is possible to maintain a more explicit record of our uncertainty in the parameter value *θ* by working with the posterior distribution directly. The posterior reflects how our initial uncertainty of the parameters - our prior, ℙ(*θ*) - has changed after witnessing the experimental data *D*. Bayes rule provides the precise rule for such an update of beliefs.

We now describe the technical details of the inference procedure performed using experimental data. We obtained experimental datasets corresponding to 1) viability of spores produced upon successful fusion between two gamete, and 2) gamete survival upon failed fusion.

When measuring the Rec8 effects on meiotic success and viability of spores produced upon gamete fusion, we genetically manipulated the Rec8 supply by varying the copy number of the *rec8* gene in P- and M-gametes (Fig.S5B). Specifically, heterothallic gamete populations carrying 0-4 copies of the *rec8* gene were induced to mate and sporulate, before plating a defined number of spores onto growth media to assess their ability to form colonies (Fig. S5C). Similarly, when measuring the Rec8 effects on survival of gametes that failed to fuse, we manipulated the copy number of the *rec8* gene in P- and M-gametes to control the Rec8 levels (Fig. S5B). Next, heterothallic gamete populations were induced to form mating pairs, but could not fuse due to deletion of the *fus1* gene. Individual mating pairs were then transferred onto growth media and the ability of each gamete to form colonies was assessed (Fig. S5D, S4D).

The set of model parameters, *θ*, is partitioned into *θ* = *θ*_*m*_ ∪ *θ*_*q*_, since their estimation is independent of each other due to experimental design and the biological processes under study. We therefore obtain two posterior distributions ℙ(*θ*_*m*_|*D*_*m*_) and ℙ(*θ*_*q*_|*D*_*q*_), where *D*_*m*_ and *D*_*q*_ are the biological measurements used to estimate *θ*_*m*_ and *θ*_*q*_ respectively. As detailed in the following sections, for both experiments we establish the probabilistic model underpinning the experimental procedure, and use it to determine the likelihood of observing the data conditioned on a parameter set ℙ(*D*|*θ*). We then define our prior beliefs about parameter values, ℙ(*θ*).

#### 2.2 Derivation of likelihood and priors for the measurements of Rec8 effects on spore viability

Suppose we have *N*_*m*_ spores that are viable and form colonies (*y* = 1) with some probability *P* . This is therefore a Bernoulli process:

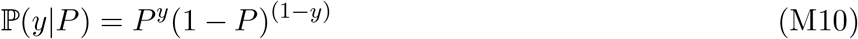

As proposed in 1.1, we model the probability of viable spores as some function of the total Rec8 levels *n* provided by both gametes to the zygote and some vector of parameters *θ*_*m*_ so that *P* = *P* (*n*; *θ*_*m*_). Given a dataset 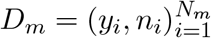 we can write down the likelihood of the data given parameters *θ*_*m*_:

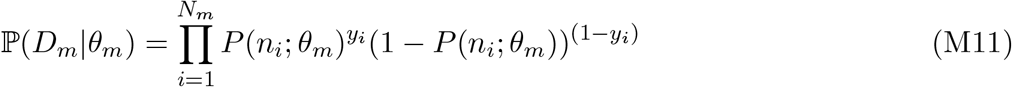

We then obtain the log likelihood

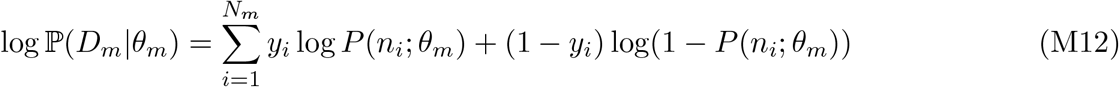

which we use in numerical procedures to estimate the posterior. As detailed in 1.1, our functional form for *P* (*n*; *θ*_*m*_) is

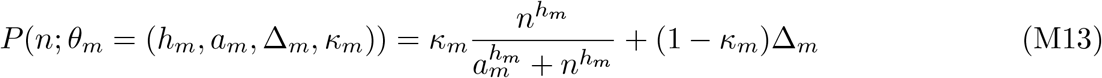

With this we obtain an explicit expression for ℙ(*D*_*m*_|*θ*_*m*_) and log ℙ(*D*_*m*_|*θ*_*m*_) through substitution into and M12, the probabilistic details of our setup unchanged.

For our priors we assume the following distributions

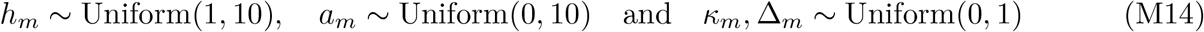

#### 2.3 Derivation of likelihood and priors for the measurements of Rec8 effects on unfused gamete survival

Suppose we have *N*_*q*_ gametes that re-enter mitosis to form colonies (*y* = 1) with some probability *q*. This is also a Bernoulli process:

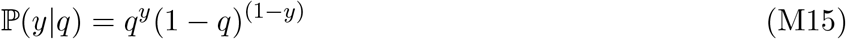

As proposed in 1.1, we model the probability that the unfused gamete is viable as some function of the Rec8 levels *r* present in the gamete and some parameters *θ*_*q*_, so that *q* = *q*(*r*; *θ*_*q*_). Given a dataset 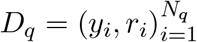 we can write down the likelihood of the data given parameters *θ*:

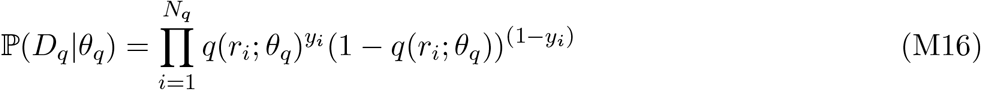

We then obtain the log likelihood

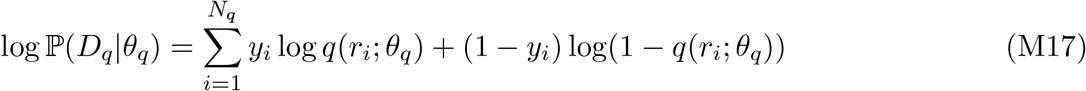

which we use in numerical procedures to estimate the posterior. As detailed in 1.1, our functional form for *q*(*r*; *θ*_*q*_) is

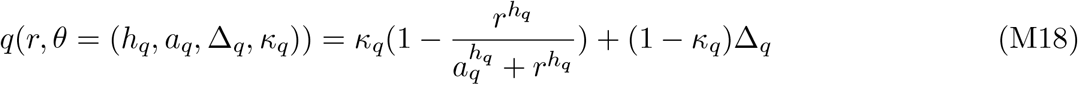

With this we obtain an explicit expression for ℙ(*D*_*q*_|*θ*_*q*_) and log ℙ(*D*_*q*_|*θ*_*q*_) through substitution into and M17, the probabilistic details of our setup unchanged.

For our priors we assume the following distributions

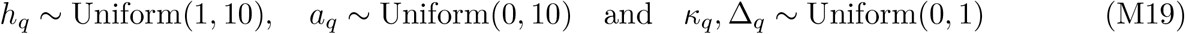

#### 2.4 Numerical procedure for estimation of posterior parameter distributions

In general, the posterior distribution cannot be obtained analytically due to the normalization constant ℙ(*D*) in the Bayes rule. However, there is a well-established and theoretically rigorous set of techniques known as Markov Chain Monte-Carlo (MCMC) that are routinely used in practical implementations of Bayesian inference to numerically draw samples from the posterior [4]. In many variants of MCMC, one is able to draw samples from any distribution given a function which is proportional to its density. We can therefore sample from our parameter posterior ℙ(*θ*|*D*) if we know a function *f* (*θ*) such that ℙ(*θ*|*D*) ∝ *f* (*θ*). We know from Bayes rule that the posterior is proportional to the product of the likelihood and the prior density. Hence, we take *f* (*θ*) = ℙ(*D*|*θ*)ℙ(*θ*), for which analytical expressions have been derived before 2.2, 2.3. Given this formalism, we use the Julia package DynamicHMC.jl^1^ to draw samples from the posterior using the “No-U-Turn Sampler” (NUTS) [5].

##### 2.4.1 Results

By drawing a 50’000 parameter sets from the posterior distributions ℙ(*θ*_*m*_|*D*_*m*_) and ℙ(*θ*_*q*_|*D*_*q*_). The marginal posterior distributions associated with each parameter are shown in fig. M2.

**Figure M2.**
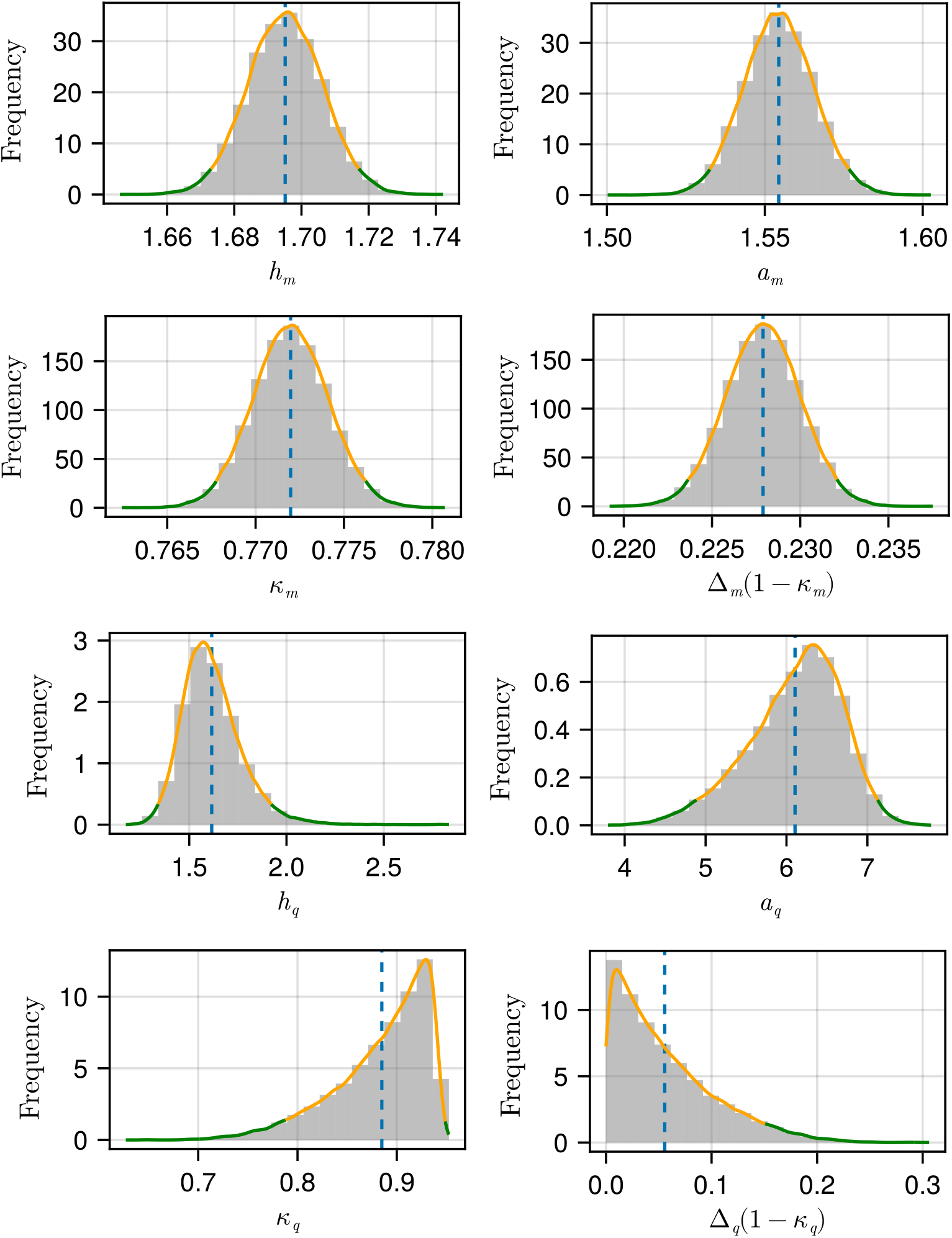
Marginal posterior distributions of parameters. Histograms show the distribution of parameter values within 50,000 posterior samples. The blue dashed lines represent the mean parameter value. The smooth green lines represent a kernel density estimate of the marginal distribution for each parameter, with the 95% credible region as determined by the highest posterior density procedure indicated by the orange outline.

The 95% credible regions as determined by the highest posterior density procedure are shown for each parameter in table M2

#### 2.5 Derivation of the posterior distribution of *n*_*c*_

To investigate whether the posterior distributions for the parameters that define *P* (.) and *q*(.) are consistent with the evolution of asymmetric provisioning in Rec8, we sought to calculate the posterior distribution of *n*_*c*_, the critical value of *n* above which asymmetries evolve (see Section 1.3.1). To achieve this, we sampled 10,000 parameter sets from the posterior distributions we obtained (see Section 2.4.1). For each parameter set sampled from the posterior distribution we determine the critical value *n*_*c*_(*ϵ*) using the procedure described in 1.3.1.

**Table M2:**
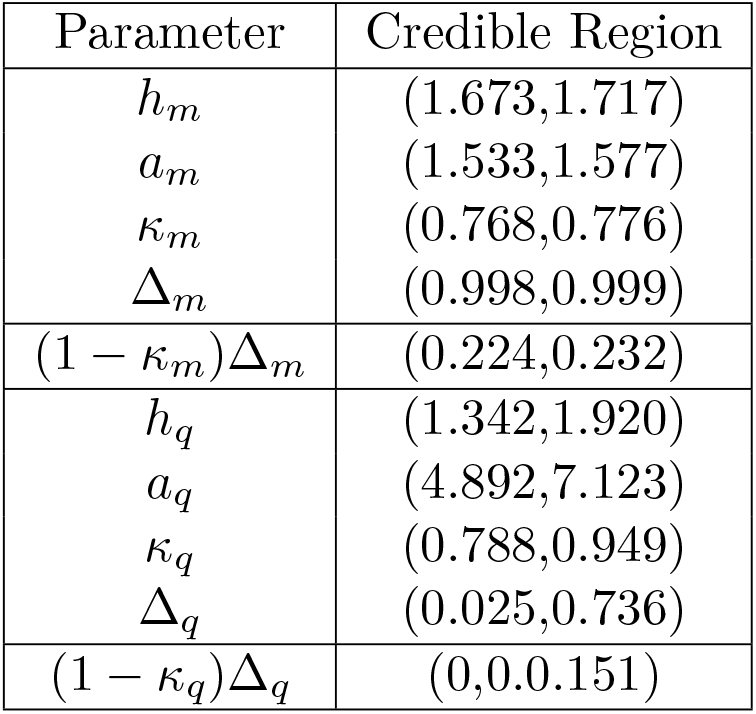
95% credible regions associated with posterior marginal distributions. A table specifying the 95% credible region obtained for each marginal distribution obtained for inferred parameters

| Parameter | Credible Region |
| --- | --- |
| $h_m$ | (1.673,1.717) |
| $a_m$ | (1.533,1.577) |
| $\kappa_m$ | (0.768,0.776) |
| $\Delta_m$ | (0.998,0.999) |
| $(1 - \kappa_m)\Delta_m$ | (0.224,0.232) |
| $h_q$ | (1.342,1.920) |
| $a_q$ | (4.892,7.123) |
| $\kappa_q$ | (0.788,0.949) |
| $\Delta_q$ | (0.025,0.736) |
| $(1 - \kappa_q)\Delta_q$ | (0,0.0.151) |

This resulted in a distribution of critical values *n*_*c*_(*ϵ*) (shown in Fig. 4H in the main text), reflecting uncertainty in the parametrization of the survival function *q* and hence the functional 2[*q*(*r*_*p*_)+ *q*(*n* − *r*_*p*_)] from which *n*_*c*_(*ϵ*) is determined. The critical value *n*_*c*_(*ϵ*) tells us: what is the minimum total investment which implies an asymmetric investment is an evolutionary optimal strategy. Given that we observe asymmetric investment in WT, we should therefore expect that the total investment *n*_WT_ is at least as large as the *n*_*c*_(*ϵ*) implied by the inferred parameters for our model.

#### 2.6 Other factors that may impact gamete and zygote survival

The theoretical model we have developed is set up to represent gene expression that induces a fitness cost for gametes and a fitness benefit for zygotes. Therefore, our model presents a generalizable framework that we then parametrized using our experimental quantification of the fitness effects of Rec8 on gametes and zygotes. We do not rule out that additional meiotic factors have similar effects as Rec8 on gamete and zygote fitness. In fact, we have shown that another cohesin, Rec11, also has a detrimental effect on gametes and is beneficial for zygotes - albeit to a lesser extent than Rec8 (fig. M2). We therefore expect our parameter quantification and conclusions to represent a conservative lower limit for the degree of asymmetry in a broader family of meiotic factor expression in P-versus M-gametes. In this way, our analysis offers a conservative and cautious assessment of whether the observed fitness effects of Rec8 on gametes and zygotes could drive the evolution of asymmetric Rec8 expression.

**Figure M3.**
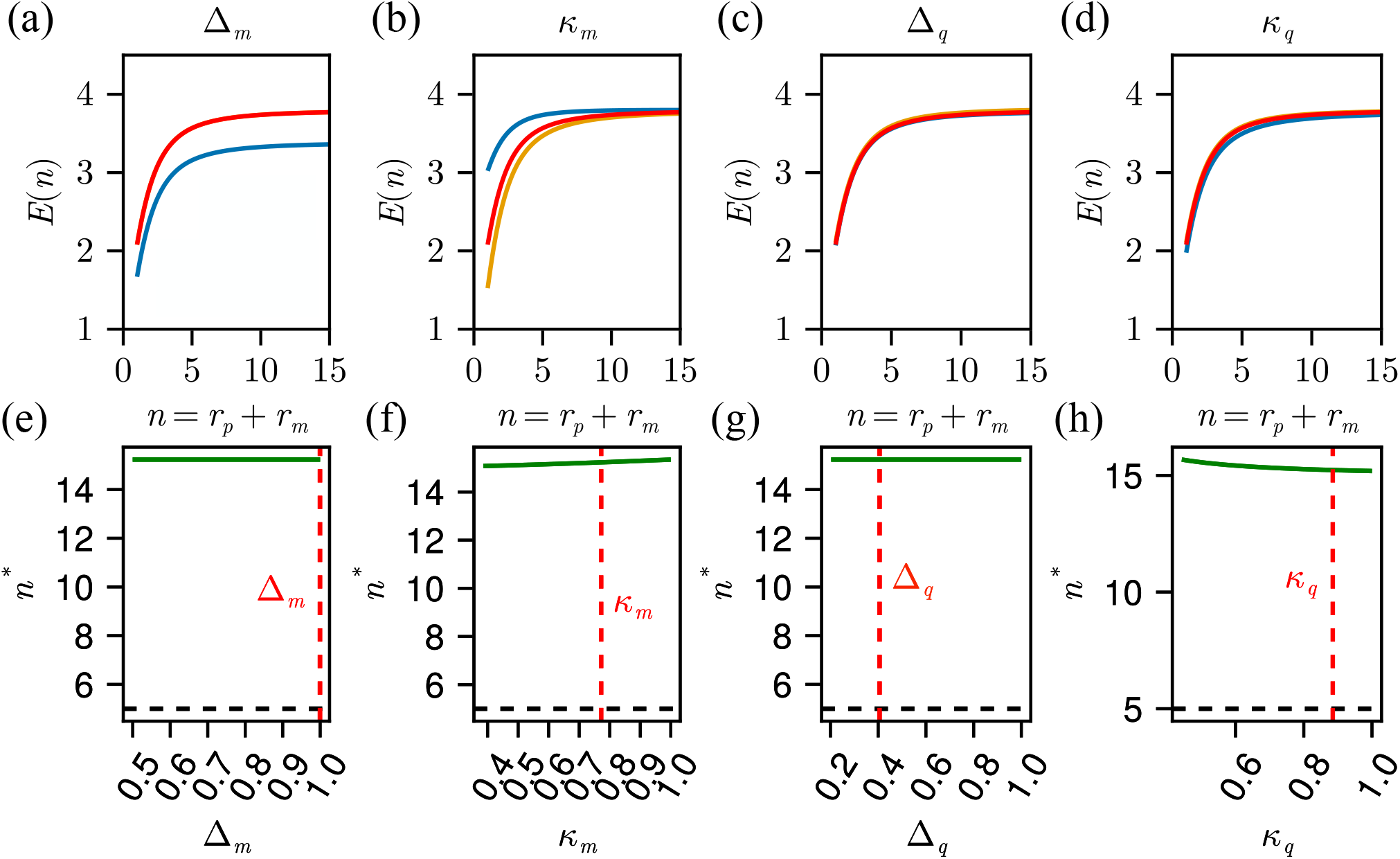
Robustness analysis near the fitted parameters. (a - d): The expected fitness *E*(*n*) as a function of the total investment of meiotic factor to the zygote while varying Δ:_*m*_ (a), *κ*_*m*_ (b), Δ:_*q*_ (c), *κ*_*q*_ (d). Red represents the fitness function *E*(*n*) for the parameters fitted from Rec8 data, orange represents the fitness function when each parameter is set to its maximal value of 1 and blue is the fitness function when we halved the fitted parameter. (e)-(h): the optimal value of the total zygotic investment while varying Δ:_*m*_ (e), *κ*_*m*_ (f), Δ:_*q*_ (g), *κ*_*q*_ (h).

Another consideration is other factors that may contribute to zygote and gamete fitness that are independent of Rec8/Rec11. For example, removing both parental *rec8* alleles does not result in complete elimination of zygotes (Fig.S7D), and high levels of *rec8* does not always lead to gamete death. To reflect other biological processes that may contribute to gamete and zygote viability, we have set up our theoretical model so that gametes and zygote fitness have some dependency on other genes (see Section 1.1).

In particular, Rec8-mediated meiotic regulation in our model is non-operational with probability 1 − *κ*_*m*_, and in these instances we assume there is a background probability of successful meiosis Δ:_*m*_ - which therefore can capture the role of alternative factors that contribute to meiosis (see section 1.1). In a similar manner, *κ*_*q*_, Δ:_*q*_ represent alternative mechanisms through which alternative factors can impact gamete survival.

These parameters are directly inferred by our experimental data (fig. M2). To directly assess how changes in these parameters, for example — reflecting the role of other genes — affects our conclusions, we performed a perturbation analysis near the fitted values of these parameters (fig. M3).

As discussed in Section 1.3.1, the critical value *n*_*c*_ is fully determined by *q*(). Furthermore, perturbing *κ*_*q*_ and Δ:_*q*_ does not affect the location of the maxima or minima for 2(*q*(*r*_*p*_) + *q*(*n* − *r*_*p*_)). Therefore, these parameters do not affect the critical value for the total investment of rec8 for which asymmetry becomes the evolutionary optimal strategy. However, perturbing these parameters may affect the optimal total investment *n*^*\**^, since they implicitly adjust the weighting between the gamete survival cost and the mating payoff in Eq.M3. This can be analyzed by examining *E*(*n*) for varying parameter perturbations (Figure M3:(a)-(d)).

The value of *n* where *E*(*n*) plateaus does not vary substantially under these perturbations, indicating that the Hill location parameters dominate over the Δ:_*i*_ and *κ*_*i*_ parameters in determining the behavior of the function. Consequently, there are only very small shifts in *n*^*\**^ (Figure M3:(e)-(h)). Therefore, we find that under these perturbations we maintain that *n*^*\**^ > *n*_*c*_, and therefore our conclusion that asymmetric investment of meiotic factors is likely to evolve as a result of the selection pressures identified in our work is robust to changes in these parameters.

#### 2.7 Figure Parameter Tables and Code Availability

Tables M3 and M4 give the model parameters used to generate each panel of Fig. 4 in the main text.

Code to run simulations, parameter estimation and produce figures can be found at this github repository, which forms a reproducible scientific project using DrWatson.jl^6^.

**Table M3:** Parameter choices for Figure 4B-4C in the main text.

| Panel | Parameter | Value |
| --- | --- | --- |
| B: triangle | $h_m$ | 2 |
| | $\Delta_m$ | 1 |
| | $\kappa_m$ | 1 |
| | $h_q$ | 2 |
| | $\Delta_q$ | 1 |
| | $\kappa_q$ | 1 |
| | $a_m$ | 1.5 |
| | $a_q$ | 6 |
| | $L_{\max}$ | 10000 |
| | $N$ | 1 |
| | $r_{\max}$ | 15 |
| B: circle | $h_m$ | 2 |
| | $\Delta_m$ | 1 |
| | $\kappa_m$ | 1 |
| | $h_q$ | 2 |
| | $\Delta_q$ | 1 |
| | $\kappa_q$ | 1 |
| | $a_m$ | 7 |
| | $a_q$ | 7 |
| | $L_{\max}$ | 10000 |
| | $N$ | 1 |
| | $r_{\max}$ | 15 |
| B: star | $h_m$ | 2 |
| | $\Delta_m$ | 1 |
| | $\kappa_m$ | 1 |
| | $h_q$ | 2 |
| | $\Delta_q$ | 1 |
| | $\kappa_q$ | 1 |
| | $a_m$ | 12 |
| | $a_q$ | 12 |
| | $L_{\max}$ | 10000 |
| | $N$ | 1 |
| | $r_{\max}$ | 15 |
| C | $h_m$ | 2 |
| | $\Delta_m$ | 1 |
| | $\kappa_m$ | 1 |
| | $h_q$ | 2 |
| | $\Delta_q$ | 1 |
| | $\kappa_q$ | 1 |

**Table M4:** Parameter choices for Figure 4D-4H in the main text.

| Panel | Parameter | Value |
| --- | --- | --- |
| D | $h_m$ | 2 |
| | $\Delta_m$ | 1 |
| | $\kappa_m$ | 1 |
| | $h_q$ | 2 |
| | $\Delta_q$ | 1 |
| | $\kappa_q$ | 1 |
| | $a_m$ | 1.5 |
| | $a_q$ | 6 |
| E-F | $h_q$ | 2 |
| | $\Delta_q$ | 1 |
| | $\kappa_q$ | 1 |
| | $a_q$ | 6 |
| G-H | $h_m$ | 1.69515 |
| | $\Delta_m$ | 0.99946 |
| | $\kappa_m$ | 0.77198 |
| | $h_q$ | 1.61557 |
| | $\Delta_q$ | 0.40584 |
| | $\kappa_q$ | 0.88496 |
| | $a_m$ | 1.5544 |
| | $a_q$ | 6.10571 |

## Footnotes

1 https://github.com/tpapp/DynamicHMC.jl

## Notes

https://doi.org/10.6084/m9.figshare.31564093

https://doi.org/10.6084/m9.figshare.31555474

### Competing Interest Statement

The authors have declared no competing interest.

